## Supplementary material for "Highly multiplexed selection of RNA aptamers against a small molecule library": All Hits Summary Supplemental

### Hits with fold >= 2.0 (11-Aug-2021 11:20:08)

**470840759 GCTGTC ACTGGA TGGACGCT TCCGGT CTGACGA GTCCTG CAACATACCCAGTTGCCCGCTAGCCC CAGGAC GAAACAGC**

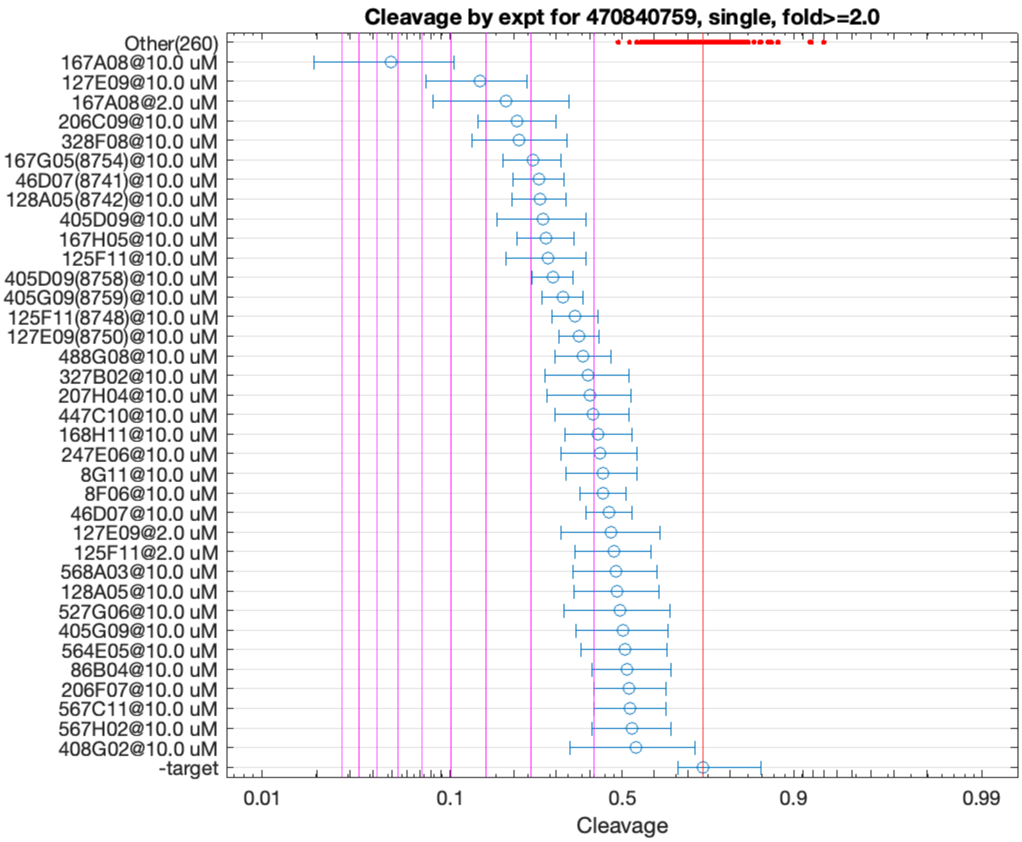


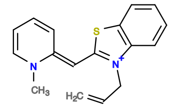

167A08

v>3.1 s=7.5

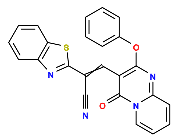

127E09

v>2.1 s=4.1

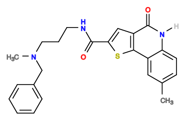

328F08

v>2.2 s=3.2

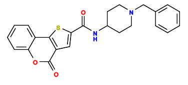

206C09

v>2.6 s=3.2

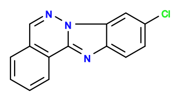

167G05

v>2.2 s=3.0

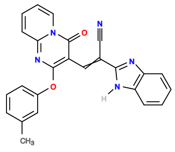

128A05

v>1.8 s=2.9

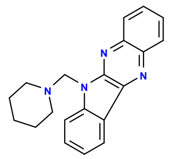

46D07

v>2.2 s=2.9

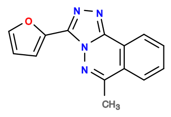

405D09

v>2.5 s=2.8

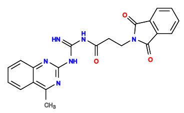

125F11

v>1.9 s=2.7

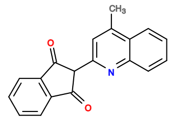

167H05

v>1.7 s=2.7

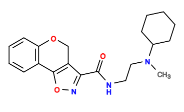

207H09

v>2.6

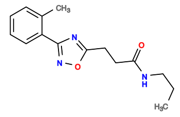

405A07

v>2.6

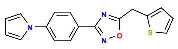

125B05

v>2.6

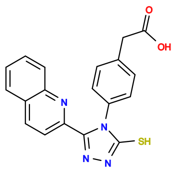

168C07

v>2.6

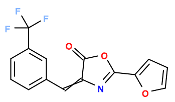

564A05

v>2.6

471146339 GCTGTC ACCGGA GCGGAGA TCTGGT CTGATGA GTCC TGGATACTGAGTAACCACACCGTTACCGCGC GGAC GAAACAGC

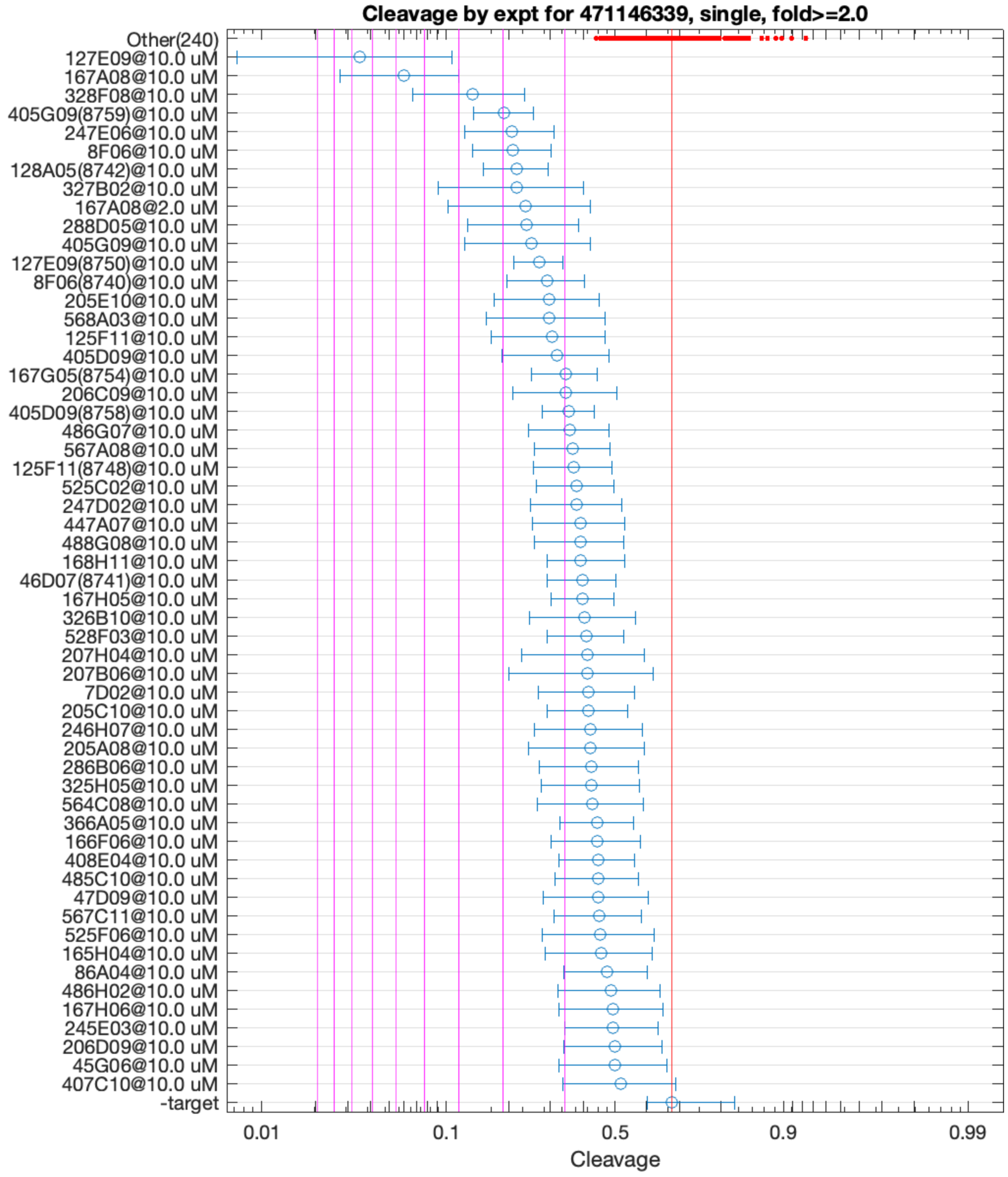


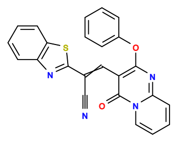

127E09

v>1.7 s=7.5

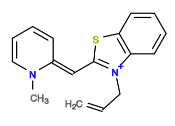

167A08

v>2.3 s=5.6

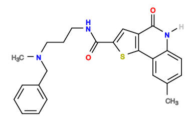

328F08

v>1.6 s=3.6

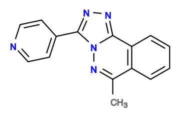

405G09

v>1.9 s=3.0

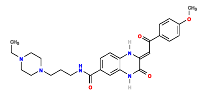

247E06

v>1.8 s=2.8

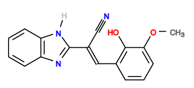

8F06

v>1.4 s=2.8

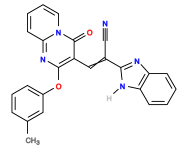

128A05

v>1.1 s=2.8

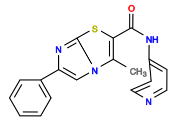

288D05

v>1.3 s=2.6

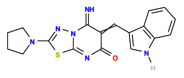

327B02

v>1.0 s=2.6

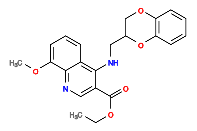

205E10

v>1.7 s=2.2

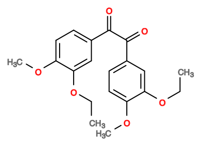

568A03

v>1.2 s=2.2

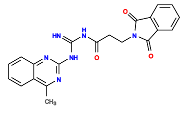

125F11

v>1.2 s=2.1

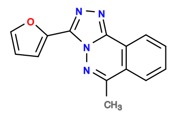

405D09

v>1.6 s=2.1

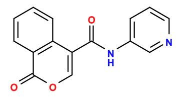

206C11

v>2.0

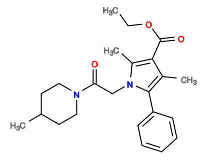

325D07

v>2.0

**472067216 GCTGTC ACTGGA GGGAAAG TCTGGT CTGATGA GTCC TTGTCATCGCCGCCACGGATGATGTTGCCC GGAC GAAACAGC**

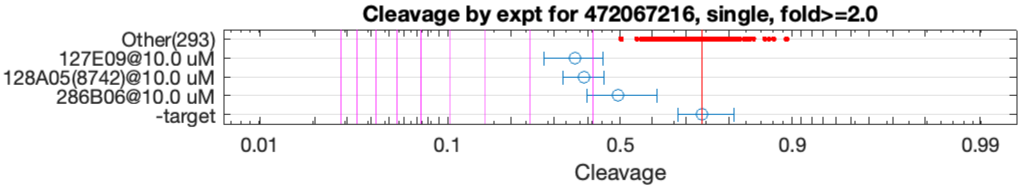


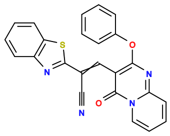

127E09

v>1.2 s=2.3

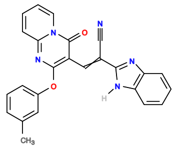

128A05

v>1.2 s=2.2

**472764094 GCTGTC ACAGA ATCGCGAG TCTGT CTGATGA GTCC ACGATGGTGCCCCGATAGCGCCCTCTCCGC GGAC GAAACAGC**

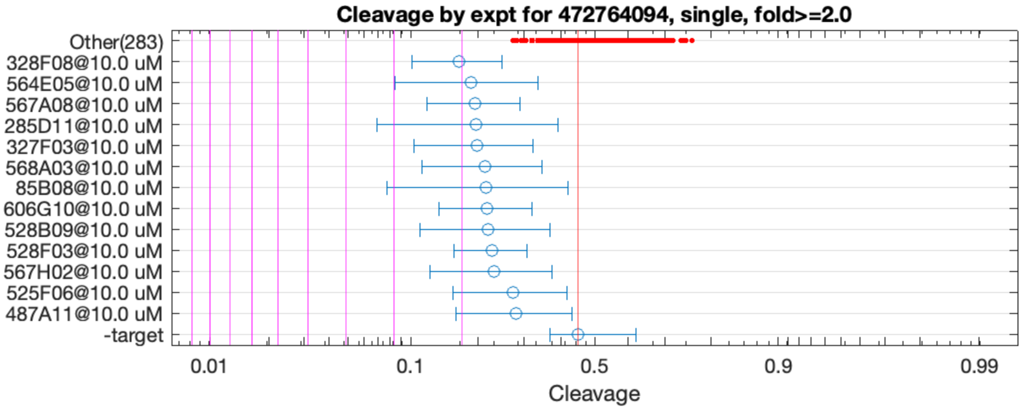


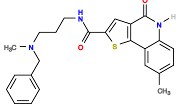

328F08

v>0.8 s=2.0

**472766050 GCTGTC ACCGGA ACGCGGA TCTGGT CTGATGA GTCC GTATACCTGCTCGCCGGTGGGTACACGCGA GGAC GAAACAGC**

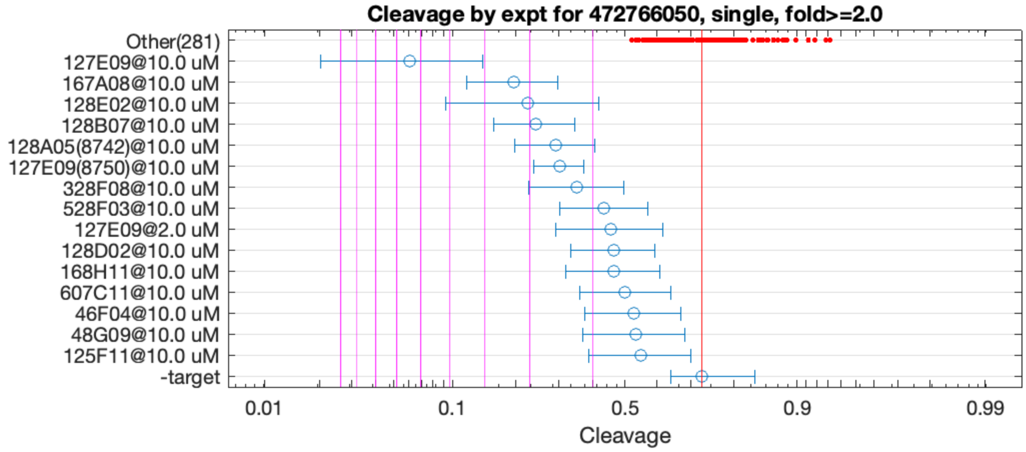


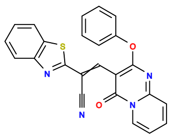

127E09

v>1.2 s=6.5

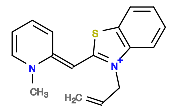

167A08

v>2.0 s=3.4

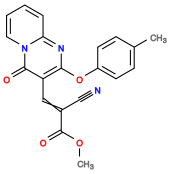

128E02

v>1.5 s=3.0

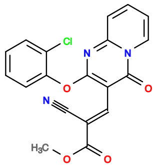

128B07

v>1.6 s=2.9

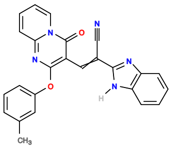

128A05

v>1.8 s=2.6

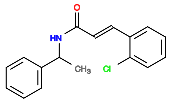

7B04

v>2.5

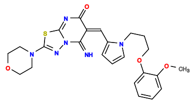

327D09

v>2.5

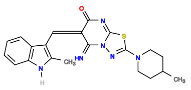

327B10

v>2.3

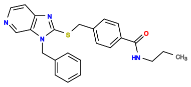

365B03

v>2.3

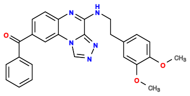

248D10

v>2.3

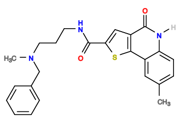

328F08

v>1.8 s=2.2

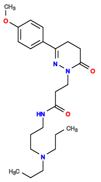

607B11

v>2.2

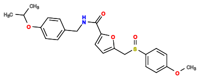

487E05

v>2.2

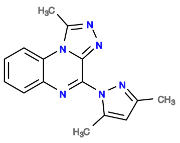

248H10

v>2.2

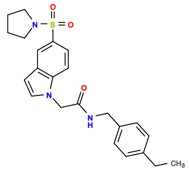

528A04

v>2.1

472786659 GCTGTC ACTGGA TGGAGGCT TCCGGT CTGATGA GTCCTG CAACATACCCAGTTGCCCGCTAGCC CAGGAC GAAACAGC

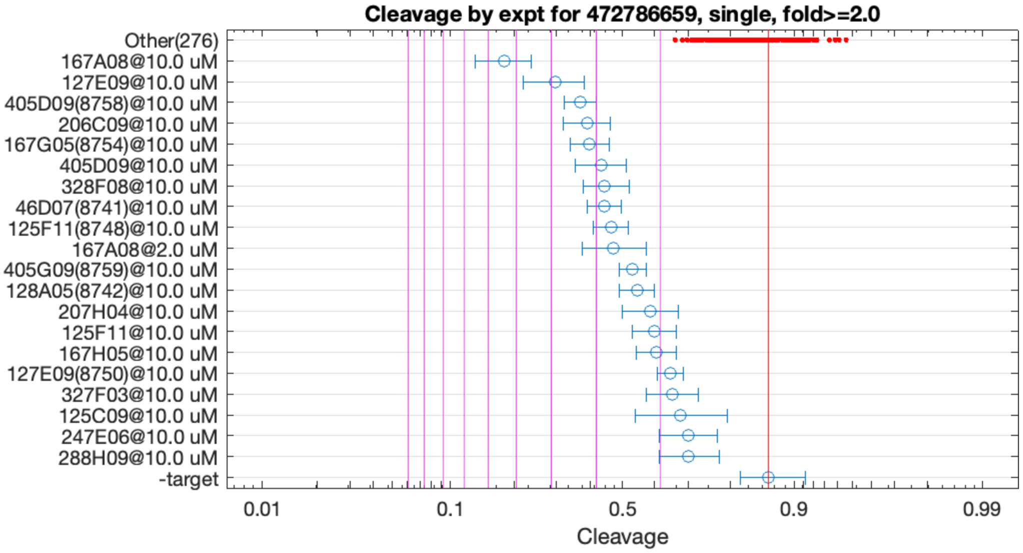


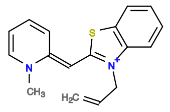

167A08

v>2.2 s=5.4

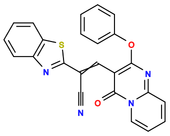

127E09

v>1.7 s=3.9

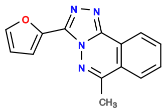

405D09

v>1.9 s=3.3

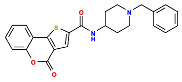

206C09

v>1.8 s=3.2

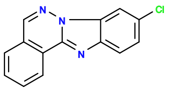

167G05

v>1.9 s=3.1

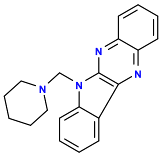

46D07

v>2.0 s=2.9

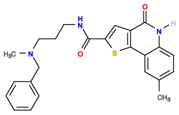

328F08

v>2.2 s=2.8

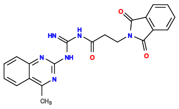

125F11

v>1.6 s=2.7

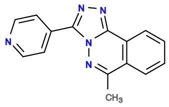

405G09

v>1.8 s=2.4

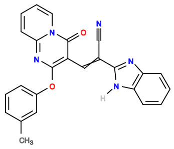

128A05

v>1.5 s=2.3

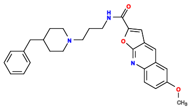

207H04

v>1.8 s=2.2

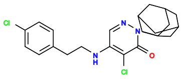

247D09

v>2.1

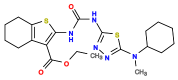

445D06

v>2.0

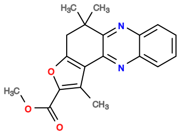

165D05

v>2.0

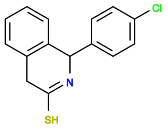

168A06

v>2.0

**472911226 GCTGTC ACAGA ATCTGGAG TCTGT CTGATGA GTCC TGCGATGCCTGATACACATGCCTCTCCGC GGAC GAAACAGC**

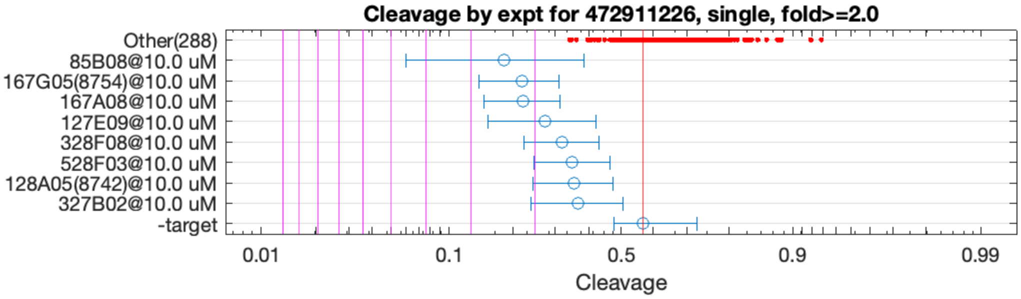


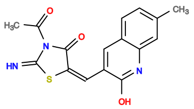

85B08

v>0.9 s=2.5

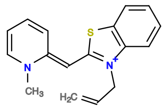

167A08

v>1.1 s=2.2

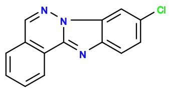

167G05

v>1.0 s=2.1

**472924986 GCTGTC ACAG GATACCGTAAA CTGT CTGATGA ATCCAAATCTGCTCGATTTCGTGTGGGTGCGGGGGAC GAAACAGC**

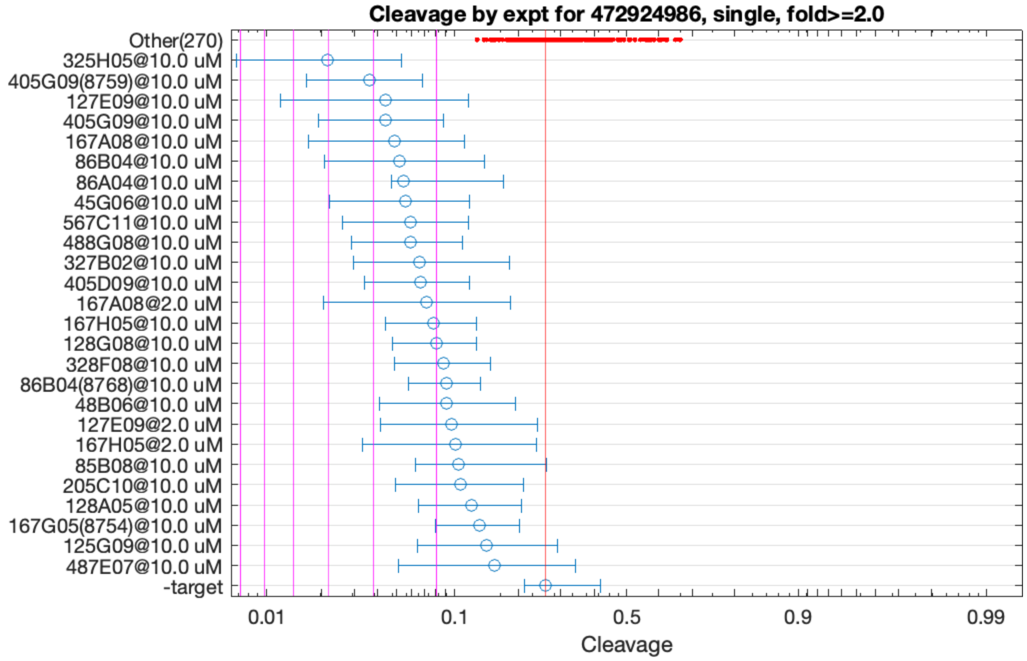


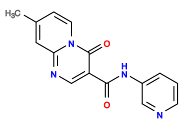

325H05

v>1.3 s=4.0

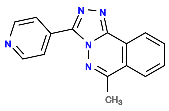

405G09

v>1.3 s=3.0

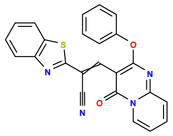

127E09

v>1.2 s=2.8

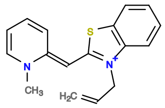

167A08

v>1.2 s=2.6

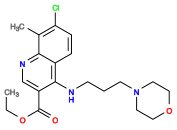

86B04

v>1.4 s=2.5

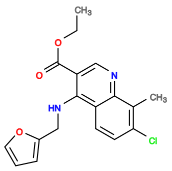

45G06

v>1.4 s=2.5

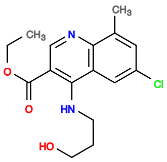

86A04

v>1.0 s=2.5

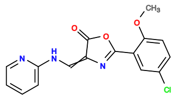

567C11

v>1.2 s=2.4

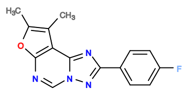

488G08

v>1.2 s=2.4

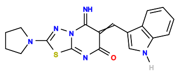

327B02

v>1.1 s=2.2

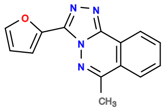

405D09

v>1.1 s=2.2

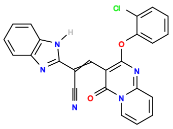

128G08

v>1.3 s=2.0

**472985468 GCTGTC ACCGGA GCGGAGA TCTGGT CTGAAGA GTCC TGGATACTGAGTAACAACACCGTTACCGCGC GGAC GAAACAGC**

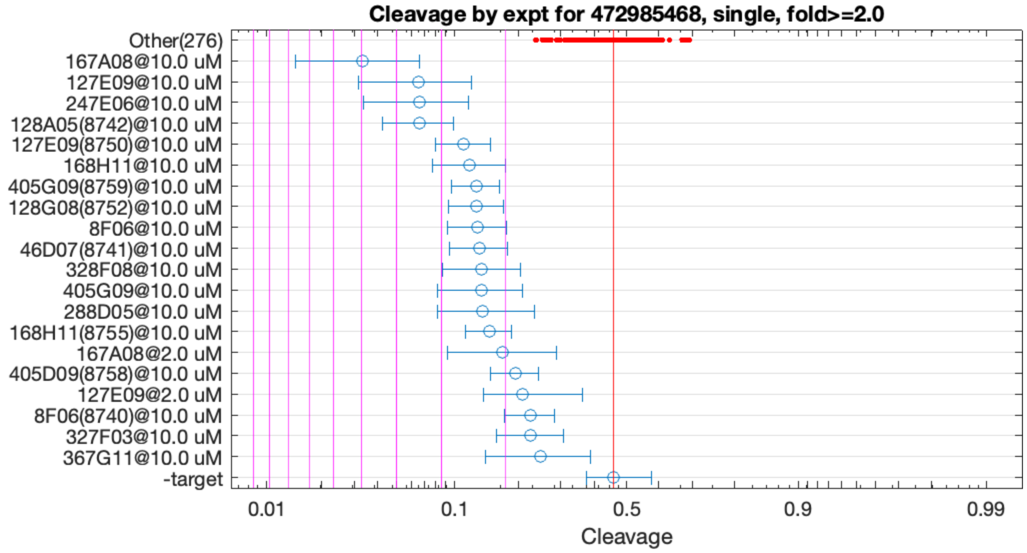


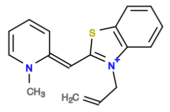

167A08

v>2.9 s=5.2

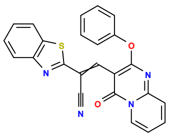

127E09

v>2.0 s=3.5

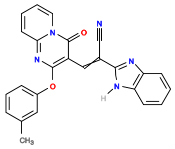

128A05

v>1.8 s=3.5

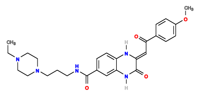

247E06

v>2.2 s=3.4

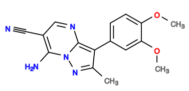

168H11

v>1.8 s=2.5

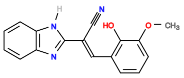

8F06

v>1.6 s=2.4

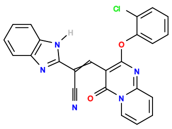

128G08

v>2.2 s=2.4

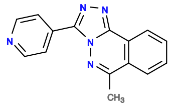

405G09

v>2.1 s=2.4

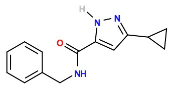

525B07

v>2.3

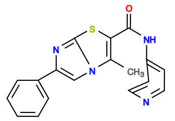

288D05

v>1.9 s=2.3

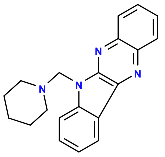

46D07

v>1.8 s=2.3

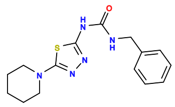

445A03

v>2.3

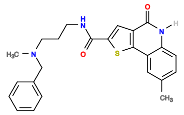

328F08

v>1.7 s=2.3

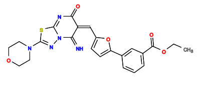

327D08

v>2.3

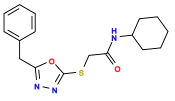

246G05

v>2.3

**472986817 GCTGTC ACCGGA GGGAAAG TCTGGT CTGATGA GTCC TTGTCATCGCCACCTCGGATGATGTTGCCC GGAC GAAACAGC**

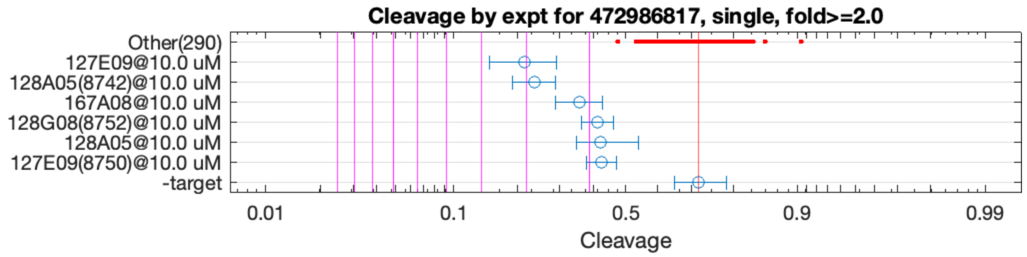


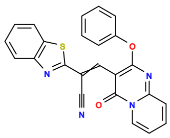

127E09

v>1.3 s=3.0

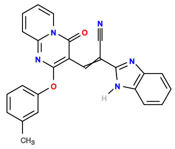

128A05

v>1.6 s=2.9

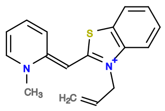

167A08

v>1.3 s=2.1

473025843 GCTGTC ACCGGA GTTCGAGTTGTCGATTCGATTGAGGGACGG TCCGGT CTGACGA GTCCA CCATCC TGGAC GAAACAGC

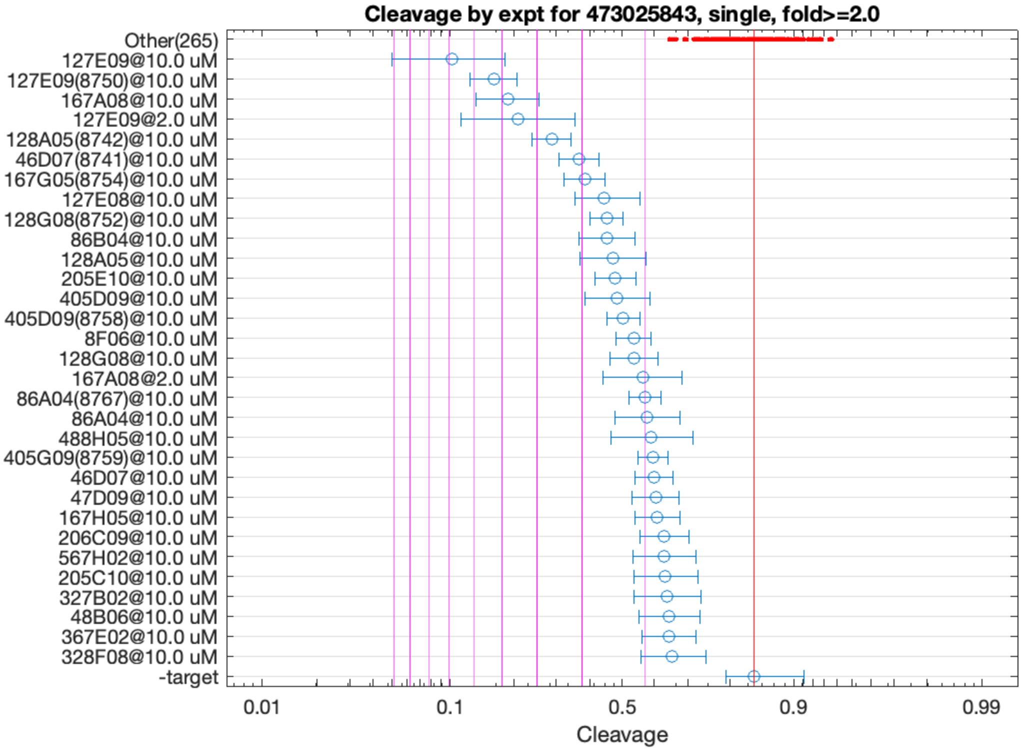


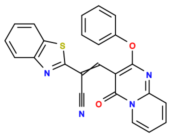

127E09

v>4.0 s=6.9

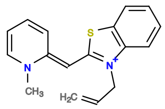

167A08

v>2.4 s=4.8

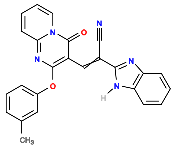

128A05

v>2.0 s=3.6

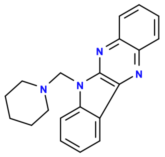

46D07

v>1.9 s=3.1

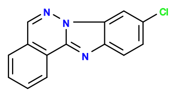

167G05

v>2.0 s=2.9

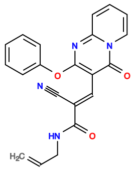

127E08

v>2.1 s=2.6

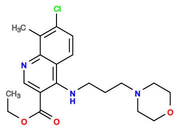

86B04

v>2.2 s=2.6

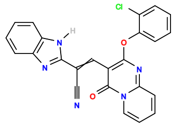

128G08

v>2.1 s=2.6

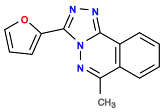

405D09

v>1.7 s=2.4

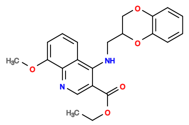

205E10

v>1.7 s=2.4

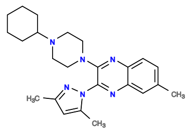

248E02

v>2.2

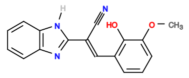

8F06

v>1.4 s=2.2

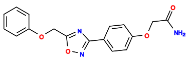

125D05

v>2.2

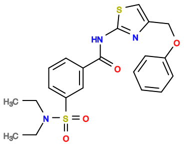

88G05

v>2.1

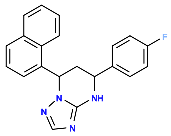

125H02

v>2.1

**473078304 GCTGTC ACTGGA GGGAAAG TCTGGT CTGATGA GTCC TTGTCATCGCCACCATGGATGATGTTGCCC GGAC GAAACAGC**

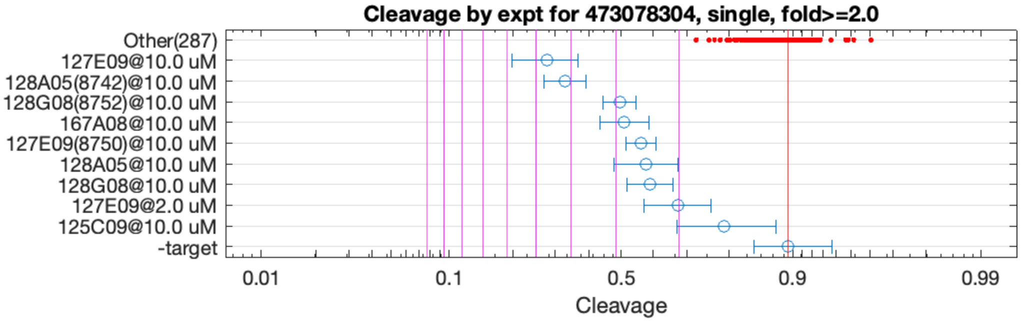


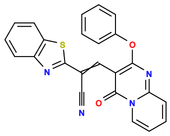

127E09

v>1.7 s=4.6

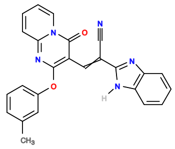

128A05

v>1.8 s=4.2

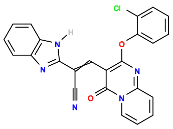

128G08

v>2.1 s=2.9

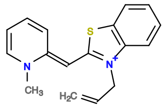

167A08

v>1.5 s=2.8

**473081357 GCTGTC ACTGGA TCCAAGCT TCCGGT CTGATGA GTCCA ATGAGGGTGCTTTCACTTCTCGCAACGG TGGAC GAAACAGC**

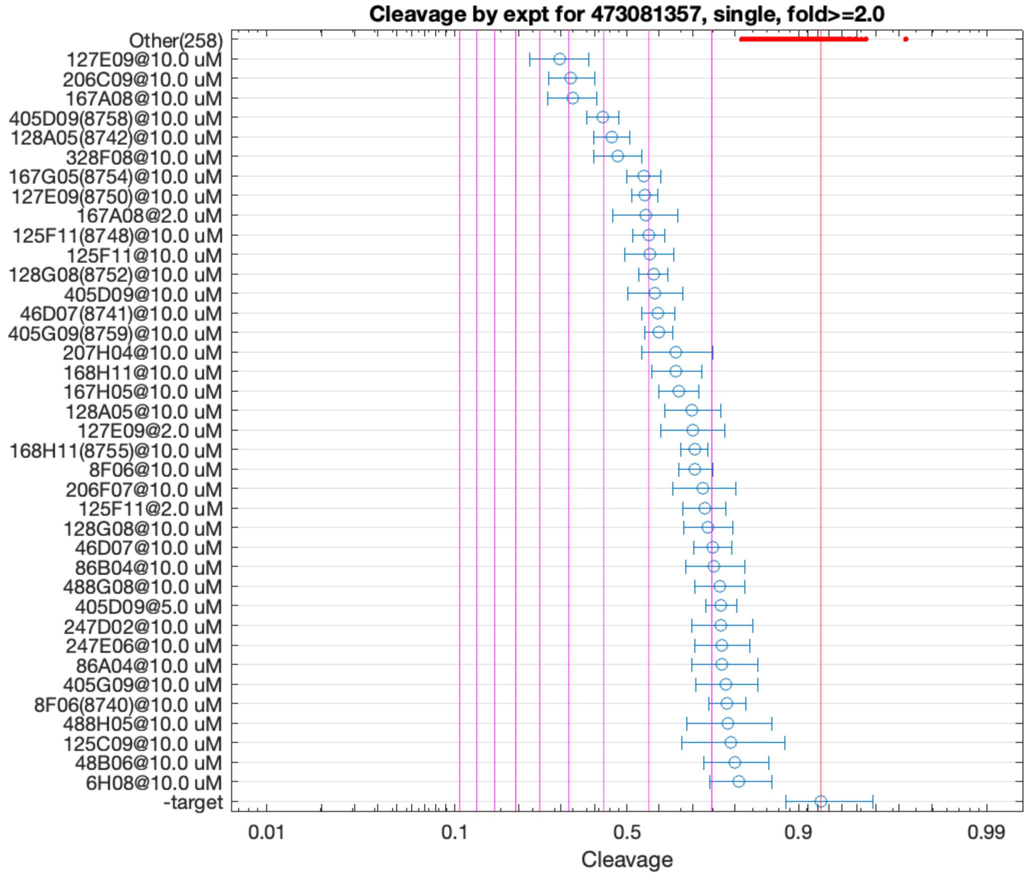


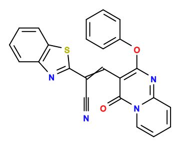

127E09

v>2.5 s=5.3

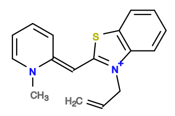

167A08

v>2.9 s=4.9

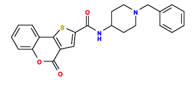

206C09

v>3.1 s=4.9

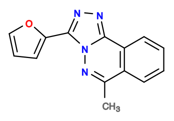

405D09

v>2.5 s=4.1

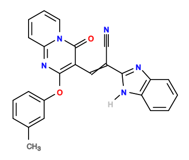

128A05

v>2.3 s=3.8

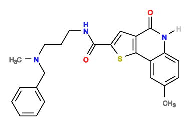

328F08

v>2.7 s=3.6

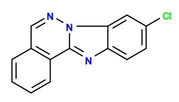

167G05

v>2.3 s=3.1

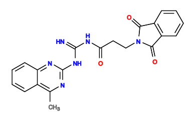

125F11

v>2.1 s=3.0

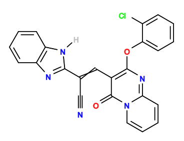

128G08

v>2.8 s=2.9

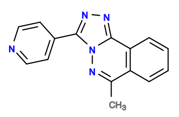

405G09

v>2.3 s=2.8

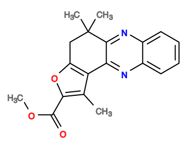

165D05

v>2.8

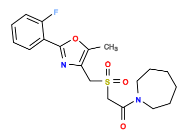

286G08

v>2.8

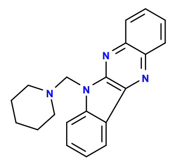

46D07

v>2.4 s=2.8

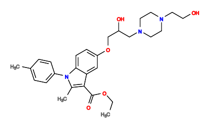

87H04

v>2.8

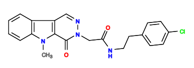

368H06

v>2.7

**473082538 GCTGTC ACTGGA TCGATGCT TCCGGT CTGACGA GTCC TTTCCCAGGCAGAAATTTGCTCGCGATCGT GGAC GAAACAGC**

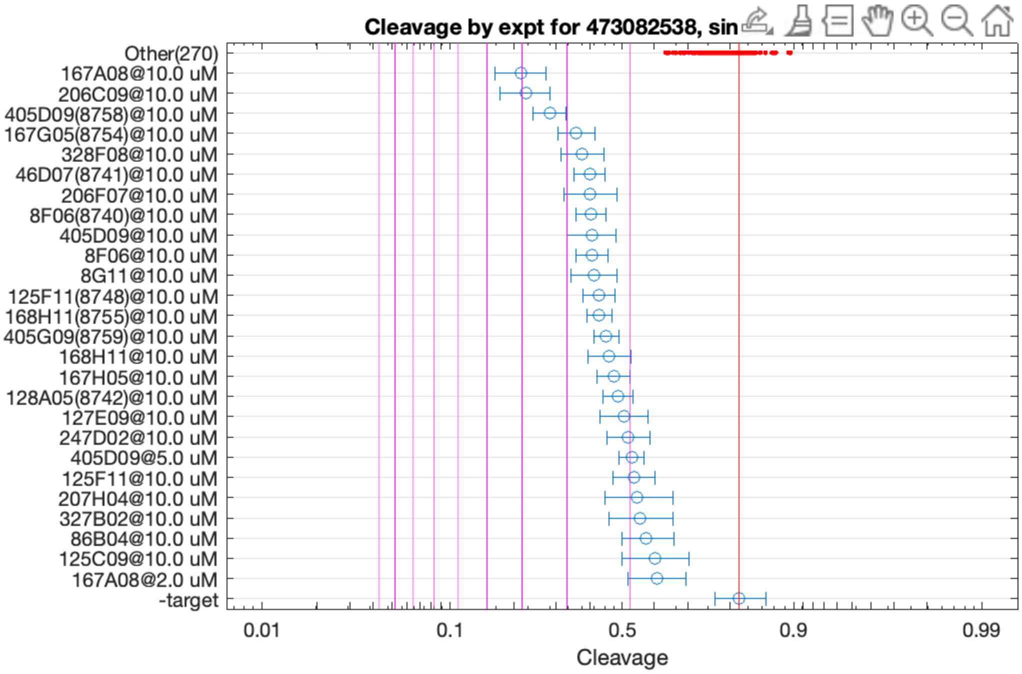


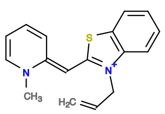
167A08
v>1.6 s=4.0

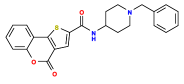
206C09
v>2.3 s=3.9

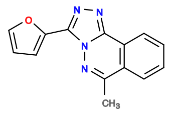
405D09
v>1.8 s=3.4

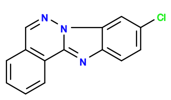
167G05
v>1.6 s=2.8

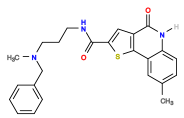
328F08
v>1.9 s=2.7

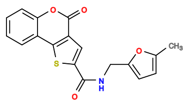
206F07
v>1.8 s=2.6

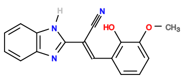
8F06
v>1.4 s=2.6

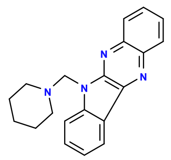
46D07
v>1.7 s=2.6

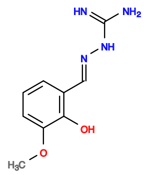
8G11
v>1.9 s=2.5

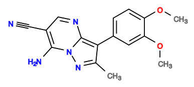
168H11
v>1.6 s=2.5

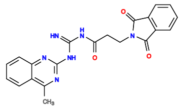
125F11
v>1.4 s=2.5

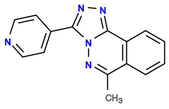
405G09
v>1.5 s=2.3

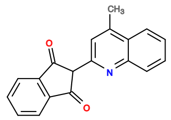
167H05
v>1.5 s=2.2

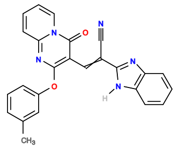
128A05
v>1.3 s=2.2

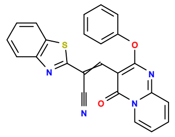
127E09
v>1.6 s=2.1

473086978 GCTGTC ACTGGA TGTGGTGC TCCGGT CTGACGA GTCC GATGTACGTTTGTCGACGCGCATCCCTTAA GGAC GAAACAGC

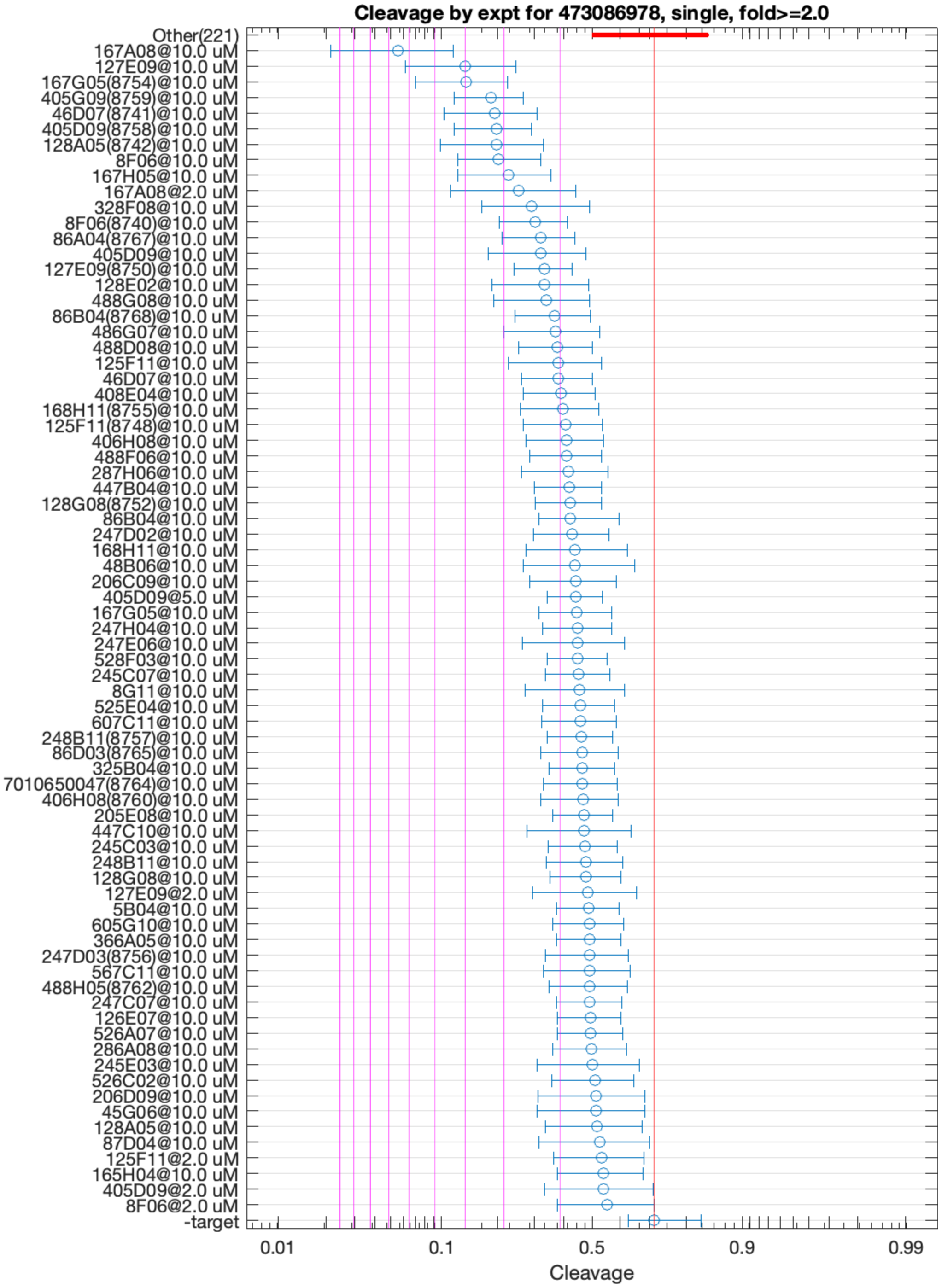


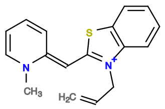

167A08

v>2.8 s=6.3

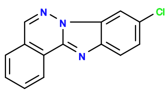

167G05

v>2.5 s=4.2

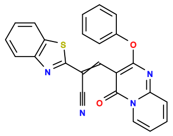

127E09

v>2.3 s=4.0

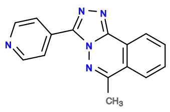

405G09

v>2.3 s=3.3

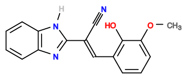

8F06

v>2.1 s=3.2

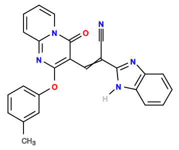

128A05

v>1.8 s=3.2

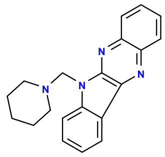

46D07

v>2.4 s=3.2

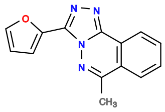

405D09

v>2.4 s=3.2

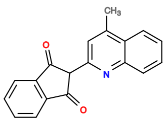

167H05

v>1.8 s=2.9

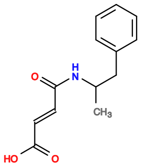

6E06

v>2.7

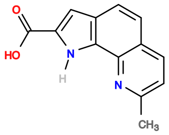

208E04

v>2.7

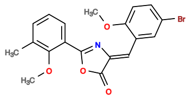

566G03

v>2.6

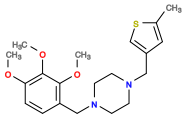

486G03

v>2.6

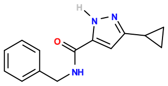

525B07

v>2.6

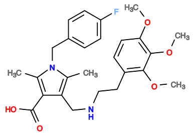

366H05

v>2.6

**473362869 GCTGTC ACTGGA TCGACGCT TCCGGT CTGACGA GTCC TTTCCCAGGCAGAAATTTGCTCGCGATCGC GGAC GAAACAGC**

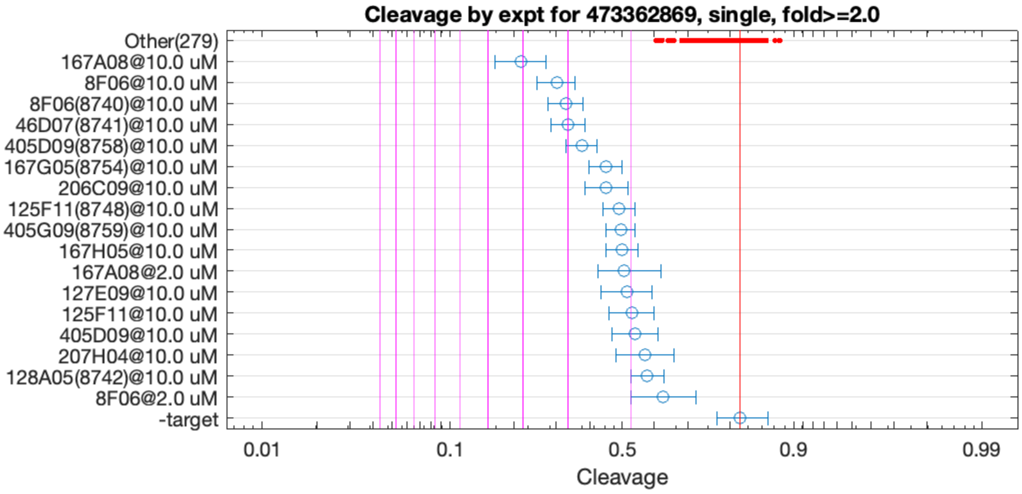


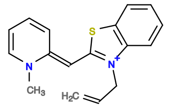

167A08

v>1.8 s=4.0

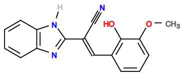

8F06

v>1.5 s=3.2

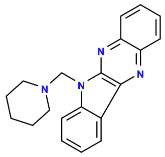

46D07

v>1.6 s=3.0

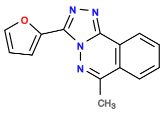

405D09

v>1.6 s=2.7

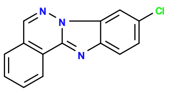

167G05

v>1.5 s=2.4

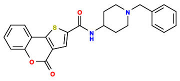

206C09

v>1.6 s=2.3

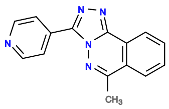

405G09

v>1.5 s=2.2

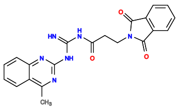

125F11

v>1.5 s=2.1

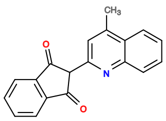

167H05

v>1.5 s=2.1

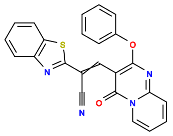

127E09

v>1.5 s=2.0

**476982823 GCTGTC AC TGGAAAACGTAAACT GT CTGATGA GTCC AAATCTGCTCGATTTCGTGTGGGTGCGTG GGAC GAAACAGC**

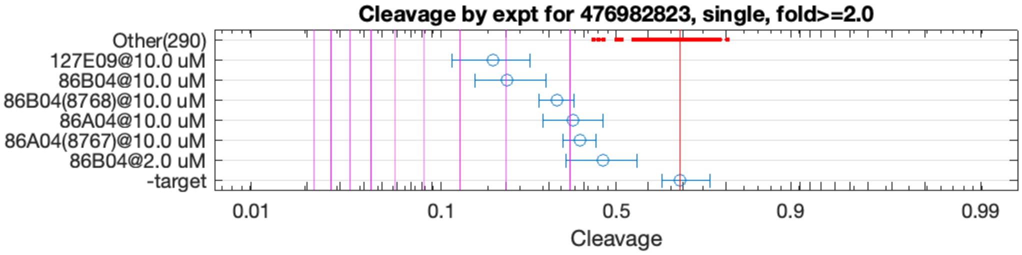


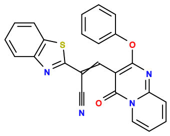

127E09

v>1.4 s=3.3

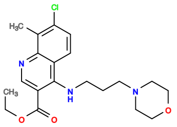

86B04

v>1.7 s=2.9

**476984717 GCTGTC ACTGGA AACGAAGG TCCGGT CTGAAGA GTCC GCCTAAGCGCTGCTCACGCAGTGGCTCGTG GGAC GAAACAGC**

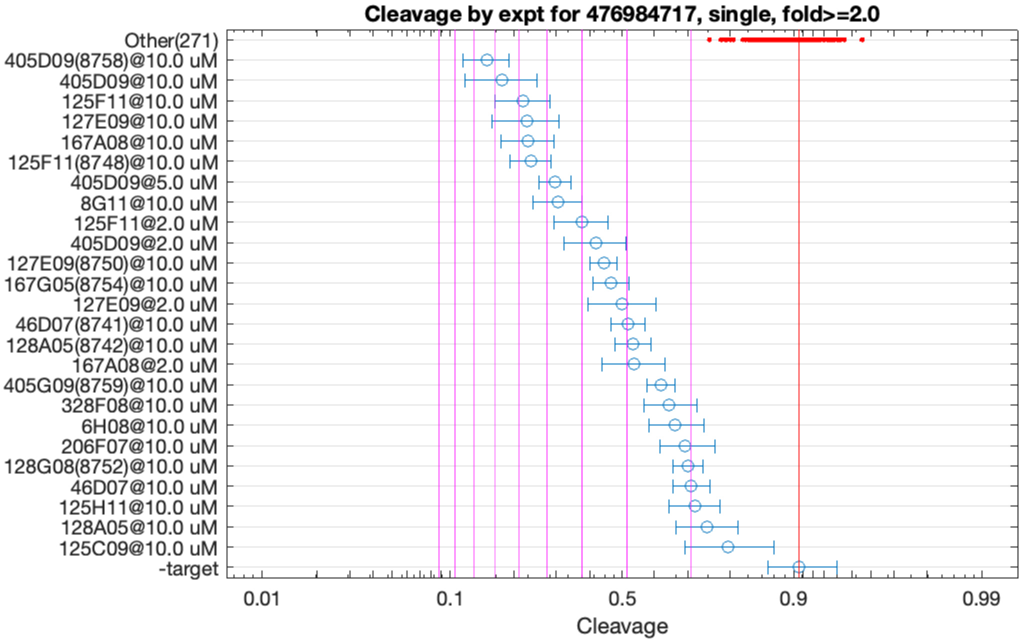


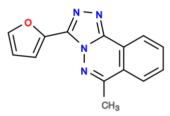

405D09

v>5.9 s=7.2

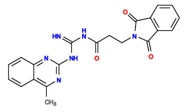

125F11

v>2.5 s=6.0

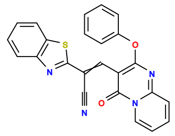

127E09

v>3.3 s=5.6

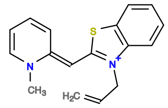

167A08

v>3.2 s=5.6

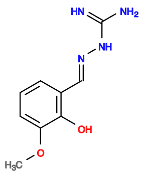

8G11

v>4.7 s=4.7

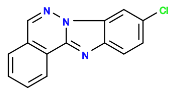

167G05

v>2.6 s=3.4

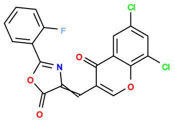

564E04

v>3.1

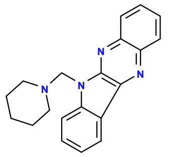

46D07

v>2.6 s=3.0

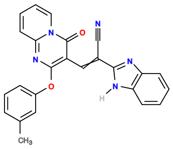

128A05

v>2.5 s=2.9

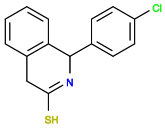

168A06

v>2.9

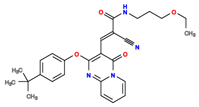

127C11

v>2.9

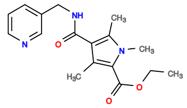

326A05

v>2.9

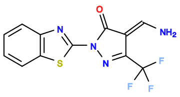

48A05

v>2.9

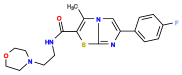

288G06

v>2.8

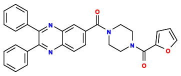

568D05

v>2.8

**477172569 GCTGTC ACTGGAT TGAACTTCTGCGTAGGTTCAGGCACCGTT ATCCGGT CCGATGA GTCC CTGGTGTA GGAC GAAACAGC**

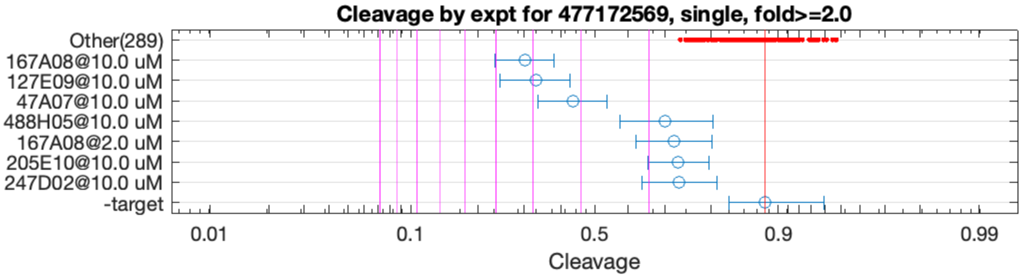


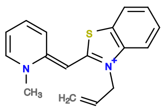

167A08

v>1.5 s=4.1

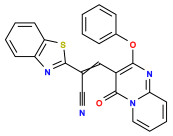

127E09

v>1.3 s=3.9

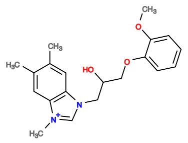

47A07

v>1.6 s=3.2

**477174303 GCTGTC AC TGGATTGGCGTAAATCC GT CTGATGA GTCC AGCGATGCCTGATACACATGCCTCTCCGCC GGAC GAAACAGC**

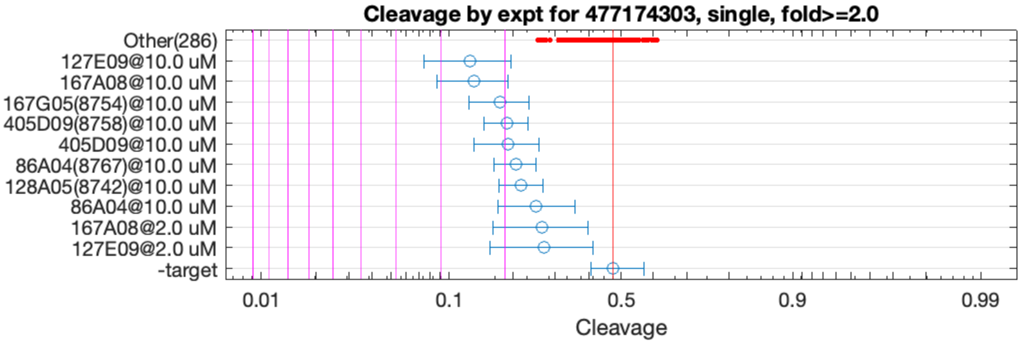


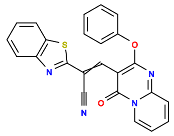

127E09

v>1.4 s=2.5

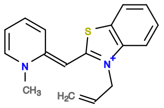

167A08

v>1.8 s=2.4

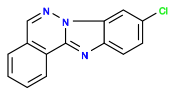

167G05

v>1.5 s=2.1

**477791857 GCTGTC ACAGGATCGGGAGA CTGTCTGA GGAGTCCTACGGCCGGAC GAAACAGC**

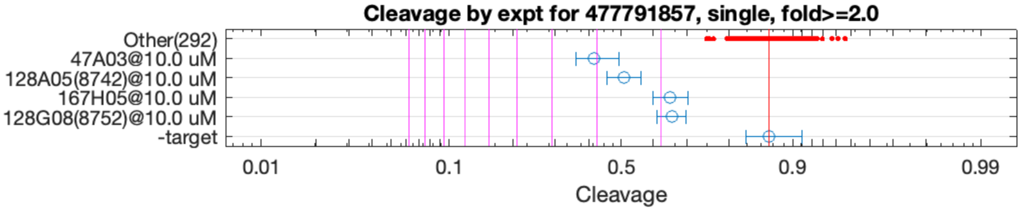


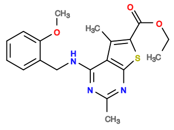

47A03

v>1.9 s=3.1

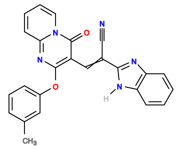

128A05

v>1.4 s=2.6

**478014511 GCTGTC ACTGGA ACGCGGA TCTGGT CTGATGA GTCC GTATACCTGCTCACCGGTGGGTACACGCGA GGAC GAAACAGC**

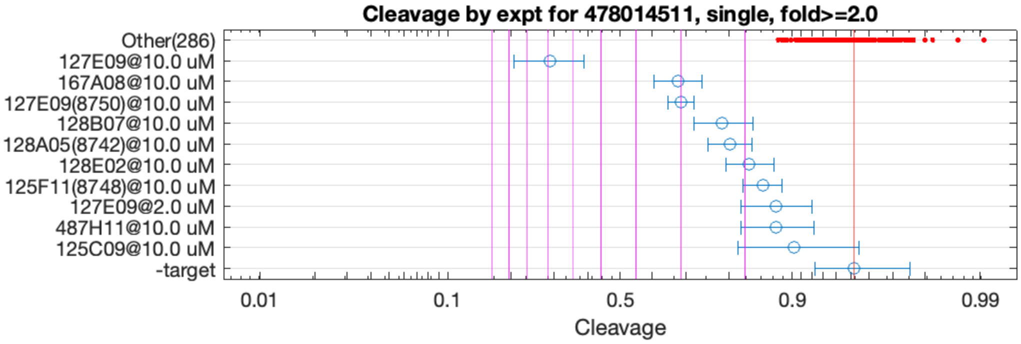


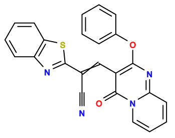

127E09

v>1.4 s=7.0

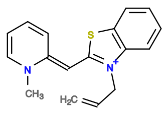

167A08

v>1.4 s=3.1

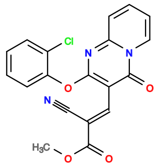

128B07

v>1.3 s=2.3

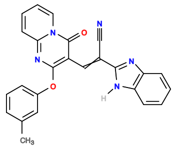

128A05

v>1.0 s=2.2

**482370107 GCTGTC ACTGGATGCGAT CTGATGA GTCC ACGTATGACGAAACAGCTCTGCCTGGCCCG GGAC GAAACAGC**

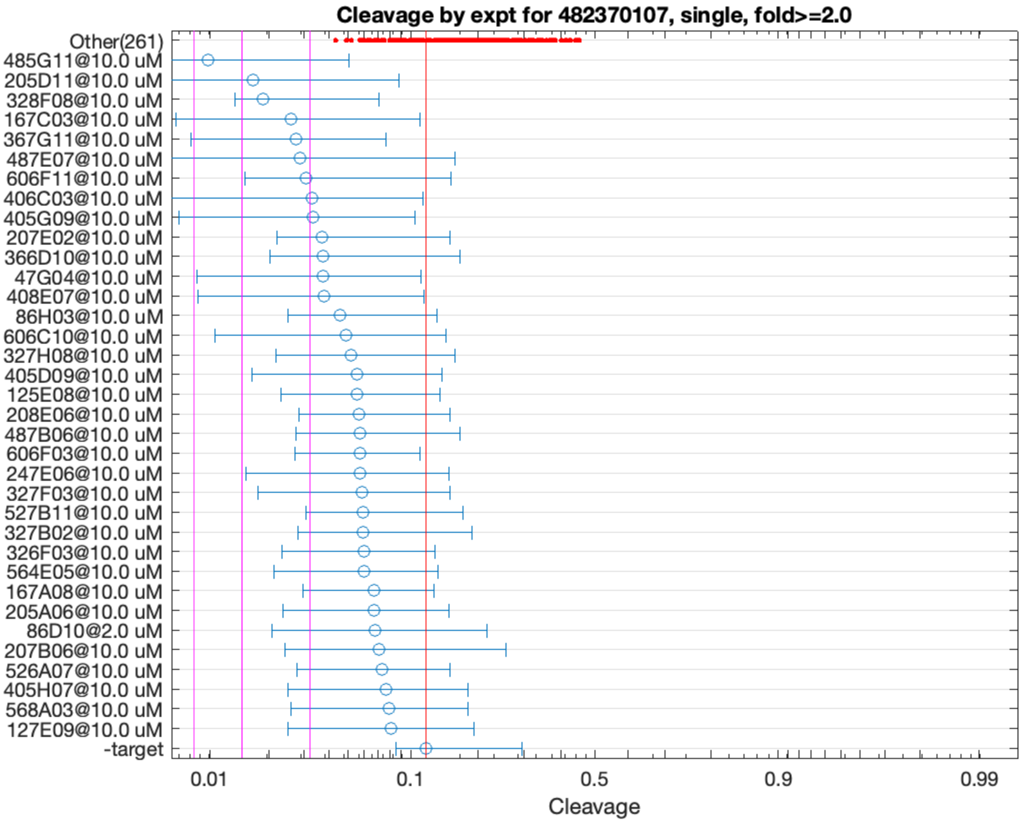


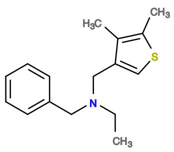

485G11

v>0.5 s=3.7

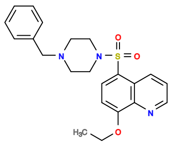

205D11

v>0.5 s=2.7

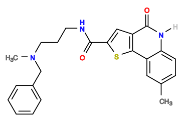

328F08

v>0.5 s=2.6

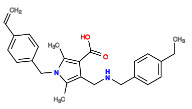

367G11

v>0.5 s=2.3

**483584637 GCTGTC ACTGGA GCGGAGA TCTGGT CTGATGA GTCC TGGATACTGAGTAACCACACCGTTACCGCGC GGAC GAAACAGC**

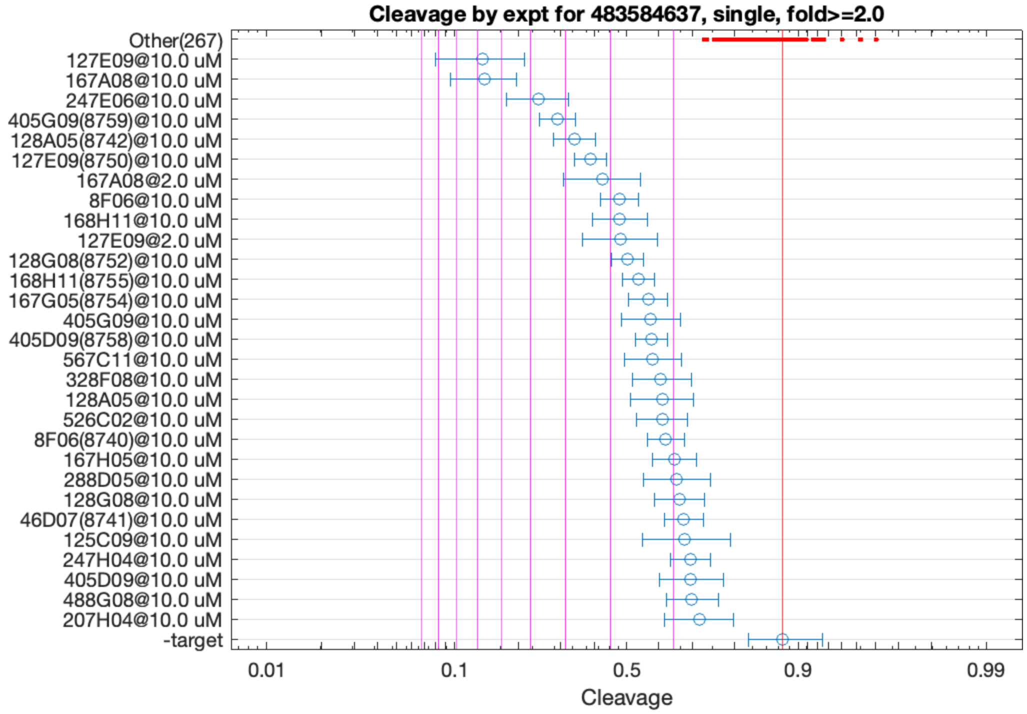


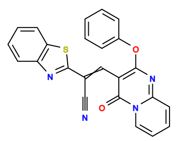

127E09

v>2.7 s=6.8

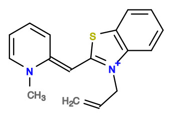

167A08

v>3.3 s=6.7

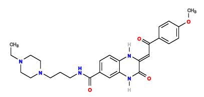

247E06

v>2.5 s=4.7

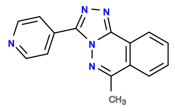

405G09

v>2.6 s=4.2

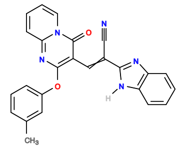

128A05

v>2.0 s=3.7

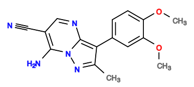

168H11

v>2.2 s=2.8

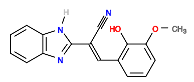

8F06

v>1.7 s=2.8

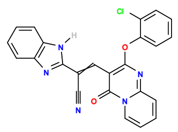

128G08

v>2.5 s=2.7

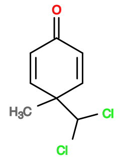

7A08

v>2.4

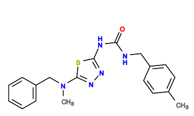

445E03

v>2.4

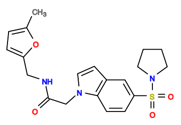

528B04

v>2.4

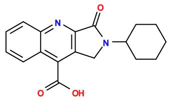

208A06

v>2.4

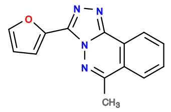

405D09

v>1.8 s=2.3

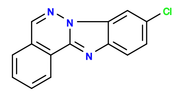

167G05

v>2.1 s=2.3

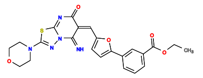

327D08

v>2.3

**485142034 GCTGTC ACTGGA GTTCGAGTTGTCGATTCGATTGAGGGACGG TCCGGT CTGACGA GTCCA CCATCC TGGAC GAAACAGC**

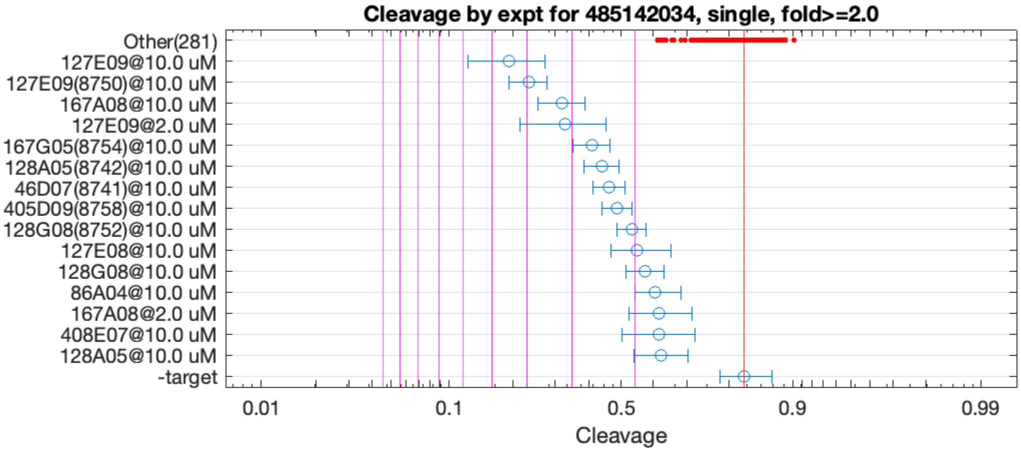


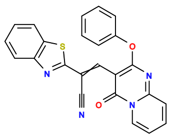

127E09

v>2.7 s=4.5

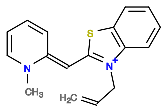

167A08

v>1.7 s=3.2

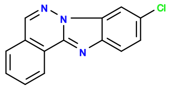

167G05

v>1.6 s=2.7

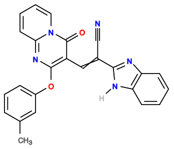

128A05

v>1.4 s=2.5

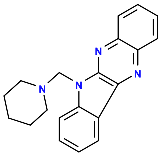

46D07

v>1.5 s=2.4

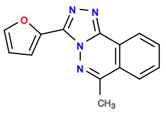

405D09

v>1.4 s=2.2

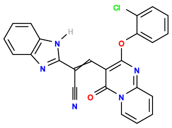

128G08

v>1.7 s=2.1

**485200498 GCTGTC ACTGGA GACGAAGG TCCAGT CTGAAGA GTCC GCCTAAGCGCTGCTCACACAGTGGCTCGTG GGAC GAAACAGC**

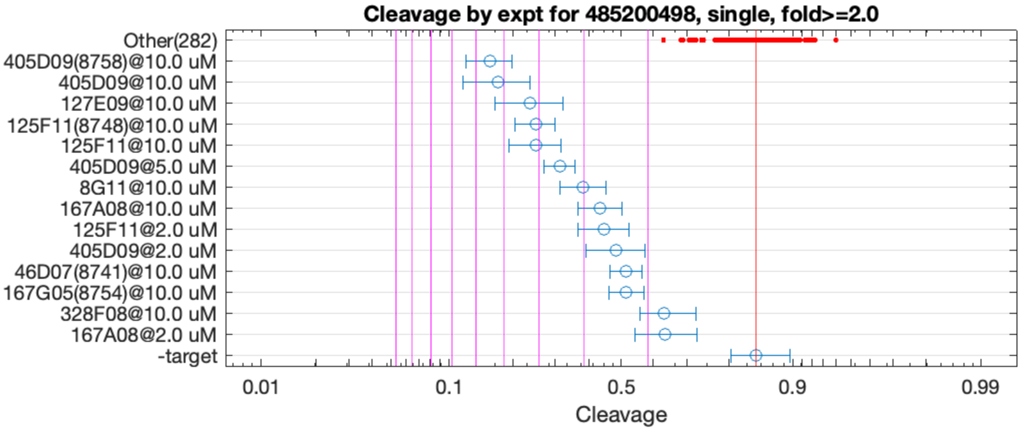


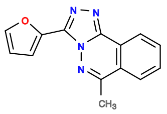

405D09

v>4.0 s=5.5

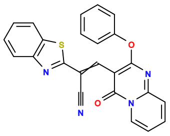

127E09

v>1.9 s=4.2

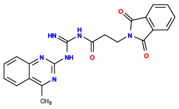

125F11

v>1.5 s=4.2

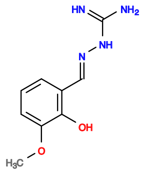

8G11

v>3.1 s=3.0

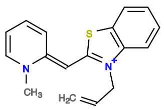

167A08

v>1.9 s=2.7

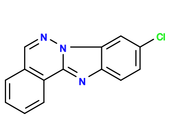

167G05

v>1.5 s=2.3

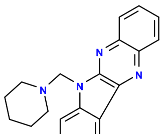

46D07

v>1.7 s=2.3

**487839220 GCTGTC ACTGGA TGCATGACCATGTTTGGTGTGCTGAGCGGC TCCGGT CTGATGA GTTC GTGGCG GGAC GAAACAGC**

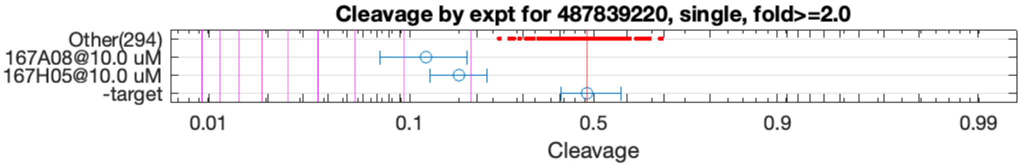


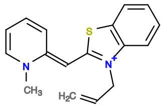

167A08

v>1.1 s=2.6

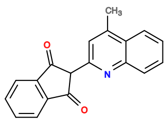

167H05

v>1.1 s=2.2

505772600 GCTGTC ACTGGA GCGGAGA TCTGGT CTGATGA GTCC TGGATTCTGAGTAACAACACCGTTACCGCGC GGAC GAAACAGC

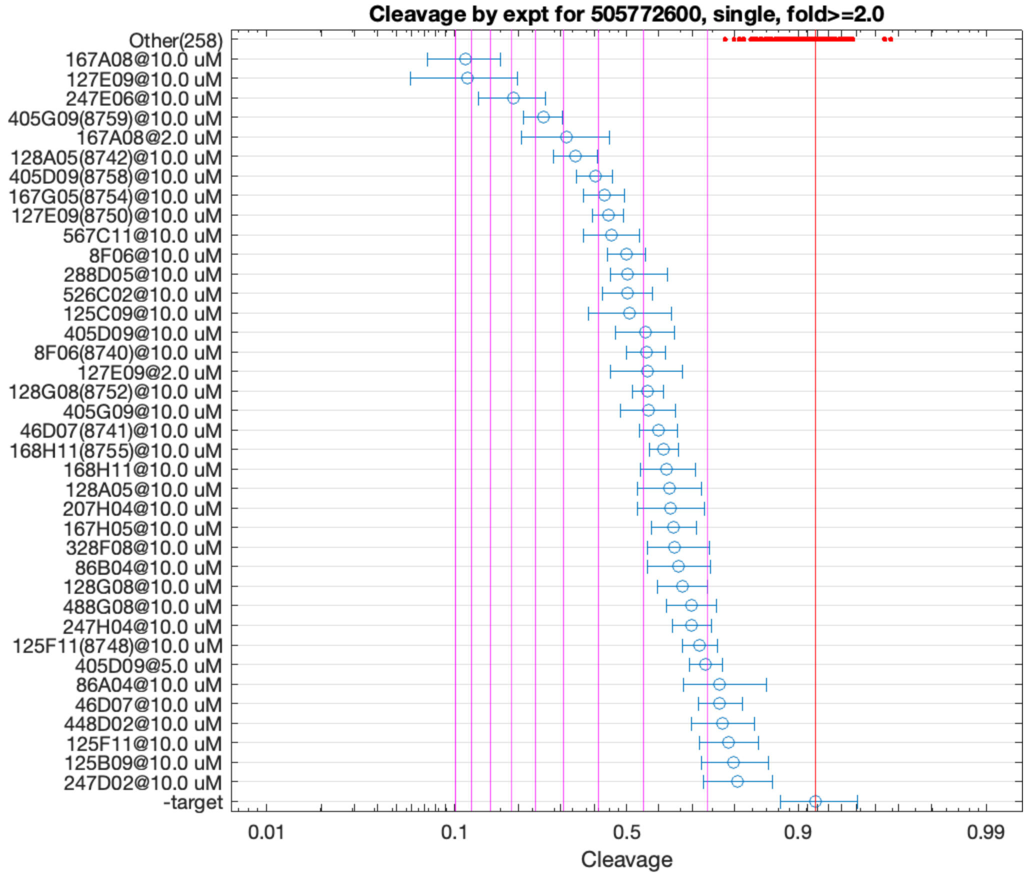


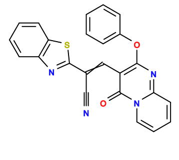

127E09

v>3.1 s=9.4

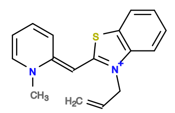

167A08

v>4.9 s=9.4

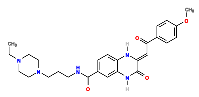

247E06

v>3.0 s=6.8

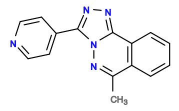

405G09

v>3.2 s=5.6

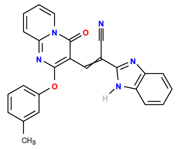

128A05

v>2.4 s=4.7

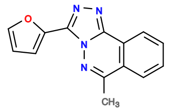

405D09

v>2.4 s=4.1

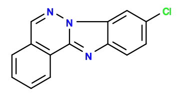

167G05

v>2.4 s=3.8

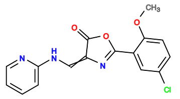

567C11

v>2.6 s=3.6

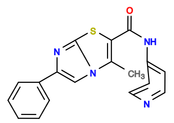

288D05

v>2.7 s=3.4

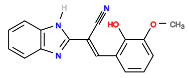

8F06

v>2.0 s=3.3

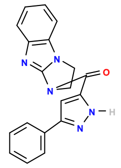

526C02

v>2.8 s=3.3

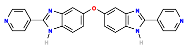

125C09

v>2.7 s=3.3

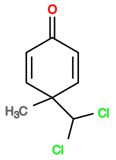

7A08

v>3.0

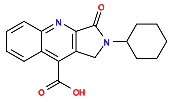

208A06

v>3.0

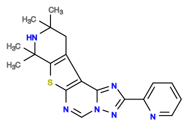

125D07

v>2.9

**510203249 GCTGTC ACTGGA AGGCACGTCTTTGAGTAGCTAAAACTGGAA TCCGGT CTGACGA GTCC GATACGTTGGCGAGTAGTTGCATGTGCCCG GGAC GAAACAGC**

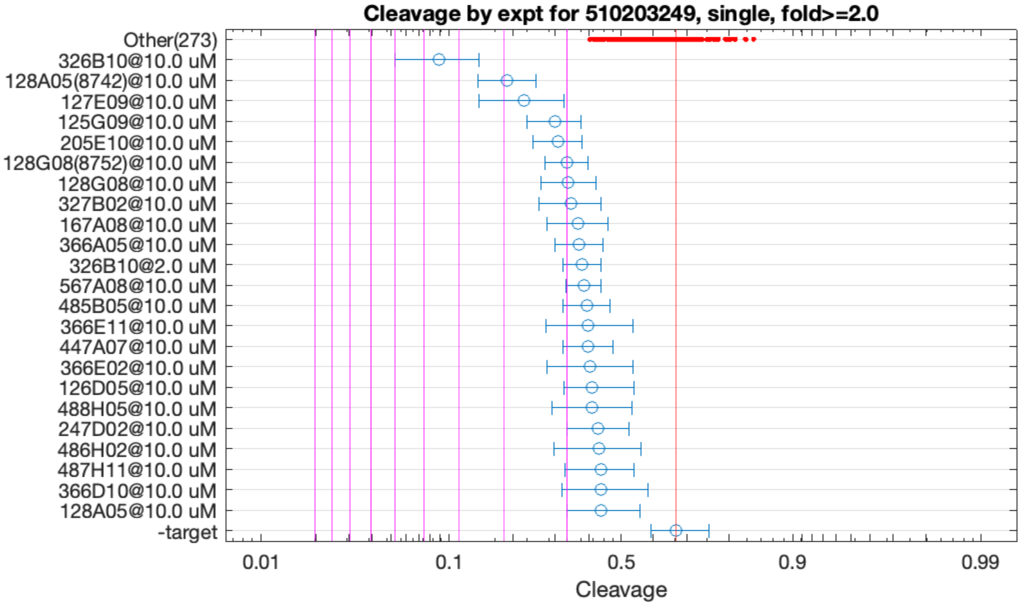


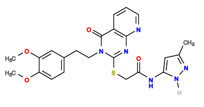

326B10

v>1.6 s=4.5

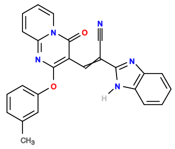

128A05

v>1.2 s=3.0

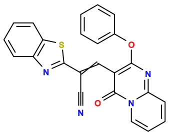

127E09

v>1.1 s=2.6

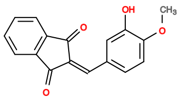

125G09

v>1.2 s=2.1

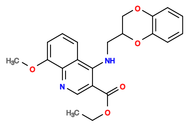

205E10

v>1.1 s=2.1

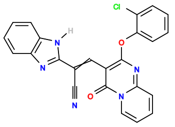

128G08

v>1.4 s=2.0

**510203337 GCTGTC ACTGGA CCTGGTGGAGAGATTACTTGGGTGTTGCGA TCCGGT CTGATGA GTCC AATACGTGTTCAAGCTTCTCCTCTTACGGC GGAC GAAACAGC**

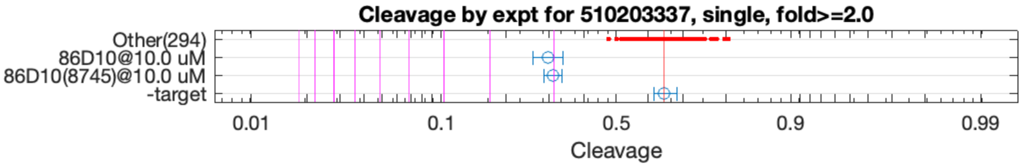


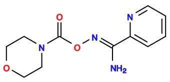

86D10

v>1.6 s=2.1

**510218739 GCTGTC ACTGGA CTTTCTCGGAGGTTTATTCGGGAAGGCGGA TCCGGT CTGATGA ATCCGTGGCGGGAC GAAACAGC**

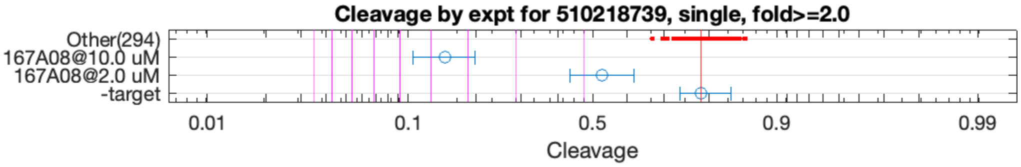


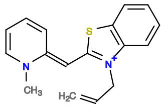

167A08

v>1.3 s=4.6

**510245864 GCTGTC ACTGGA AAACCGAACGATTGATGGACGGACACCTC TCCGGT CTGACGA GTCC GATGTGTCTGCGTGTTGGGTG GGAC GAAACAGC**

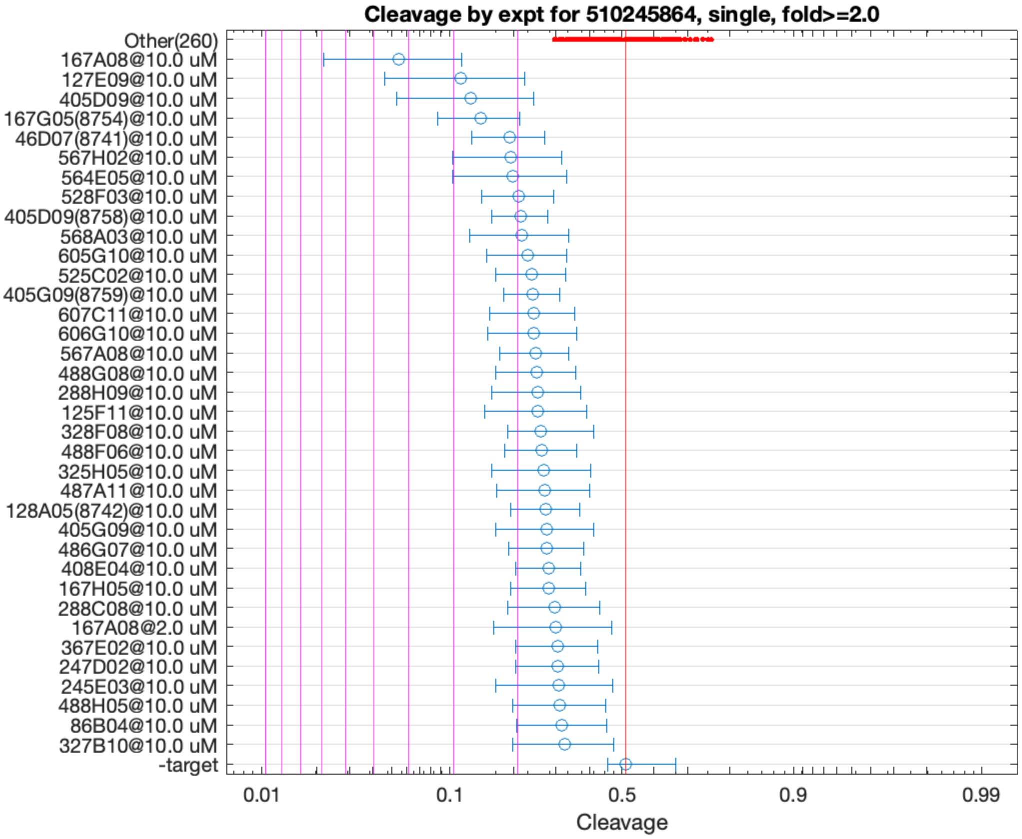


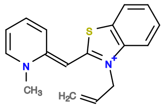

167A08

v>1.8 s=4.3

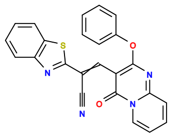

127E09

v>1.5 s=2.8

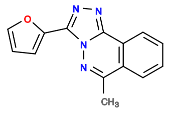

405D09

v>1.5 s=2.8

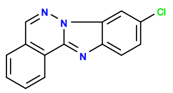

167G05

v>1.6 s=2.5

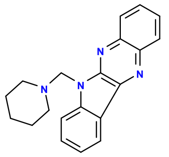

46D07

v>1.4 s=2.1

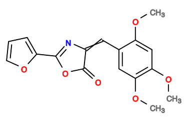

564E05

v>1.6 s=2.1

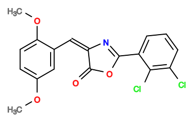

567H02

v>1.2 s=2.0

**510255226 GCTGTC ACTGGA CTAGA TTCGGT CTGAGGA ACTGTGCGACGCAGC GAAACAGC ATTAGTGCGGGACGAAACAGC**

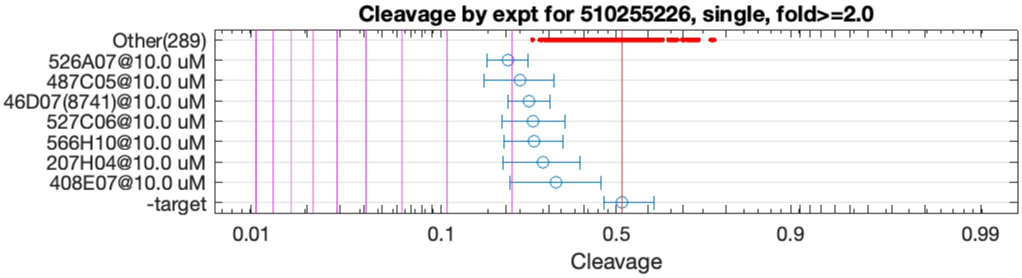


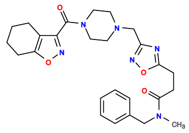

526A07

v>1.1 s=2.0

**510318329 GCTGTC ACTGG ATGCATGACCATATTTGGTGTGCCGAGCGGCA CCAGT CTGATGA GTTC GTGGCG GGAC GAAACAGC**

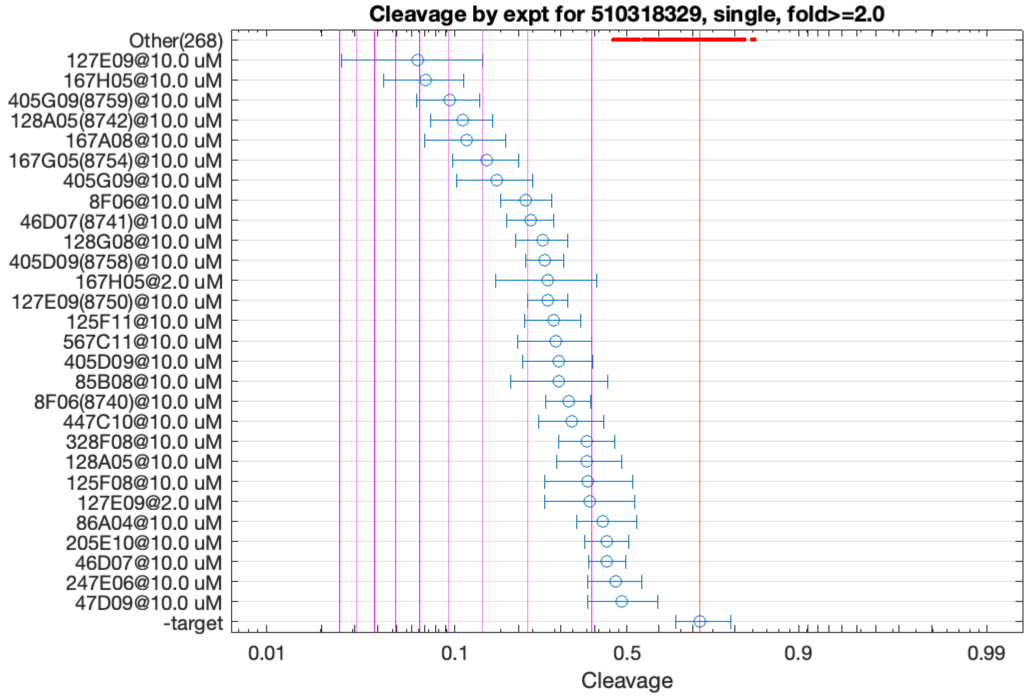


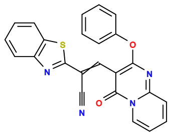

127E09

v>2.6 s=6.2

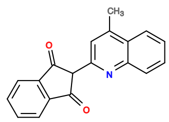

167H05

v>1.4 s=5.8

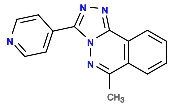

405G09

v>3.0 s=4.9

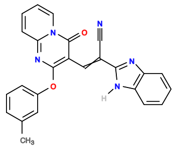

128A05

v>2.2 s=4.5

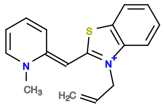

167A08

v>2.6 s=4.5

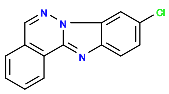

167G05

v>2.6 s=3.9

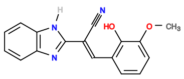

8F06

v>2.2 s=3.0

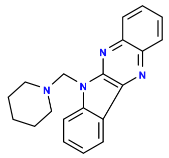

46D07

v>2.2 s=2.9

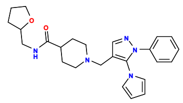

446C03

v>2.9

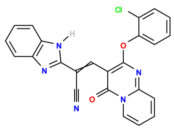

128G08

v>2.9 s=2.8

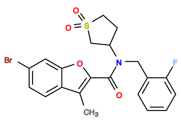

408A05

v>2.8

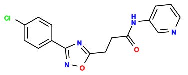

405A06

v>2.7

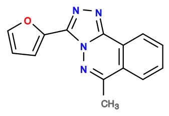

405D09

v>2.4 s=2.7

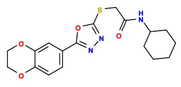

246A07

v>2.7

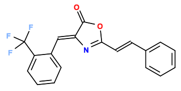

566D10

v>2.7

**510337926 GCTGTC ACTGG ATTTTCGTG TCAGT CTGATGA GTTCT TGTTGACGAAACAGCAGAGTGCCC GGGAC GAAACAGC**

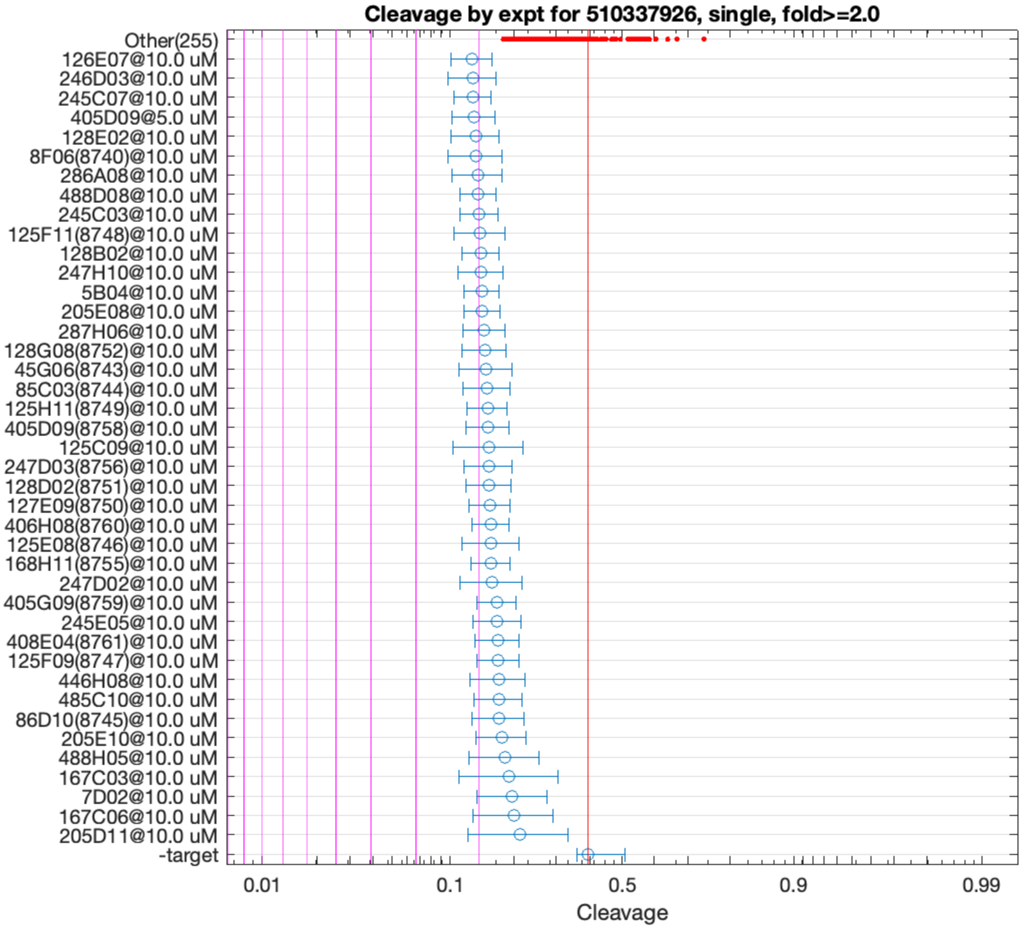


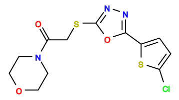

246D03

v>0.9 s=2.1

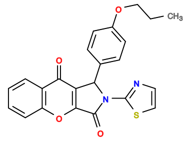

126E07

v>1.0 s=2.1

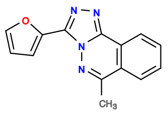

405D09

v>1.0 s=2.1

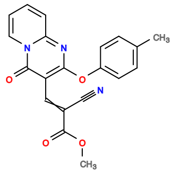

128E02

v>0.9 s=2.1

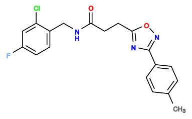

245C07

v>0.9 s=2.1

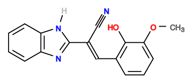

8F06

v>1.0 s=2.1

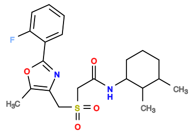

286A08

v>0.7 s=2.1

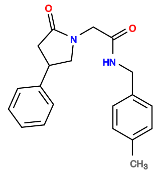

247H10

v>0.9 s=2.0

**510463768 GCTGTC ACTGGA TTTCCAAGATGATGGACAGACACCTC TCCGGT CTGATGA GTCC GATGTGTCTGCGAATGTCTGCGTGTTGGTG GGAC GAAACAGC**

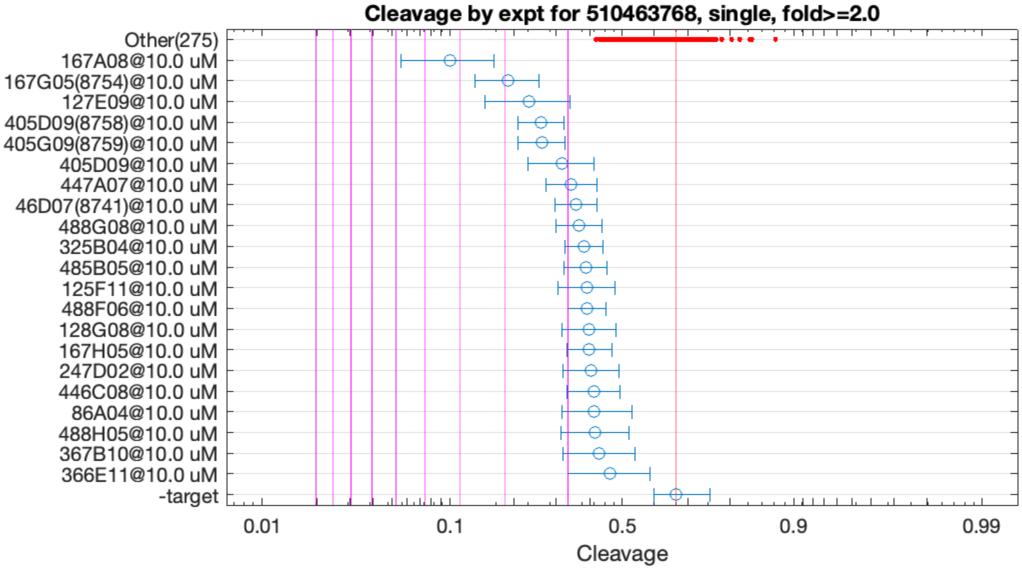


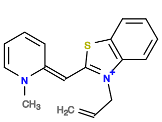

167A08

v>1.6 s=4.3

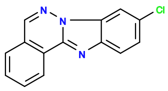

167G05

v>1.3 s=2.9

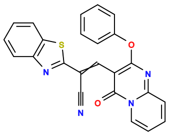

127E09

v>1.3 s=2.5

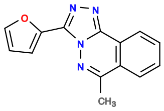

405D09

v>1.5 s=2.4

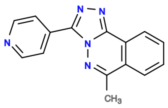

405G09

v>1.4 s=2.4

**511084246 GCTGTC ACTGGA AACGAAGG TCTAGT CTGAAGA GTCC GCCTAAGCGCTGCTCGCACAGTGGCTCGTG GGAC GAAACAGC**

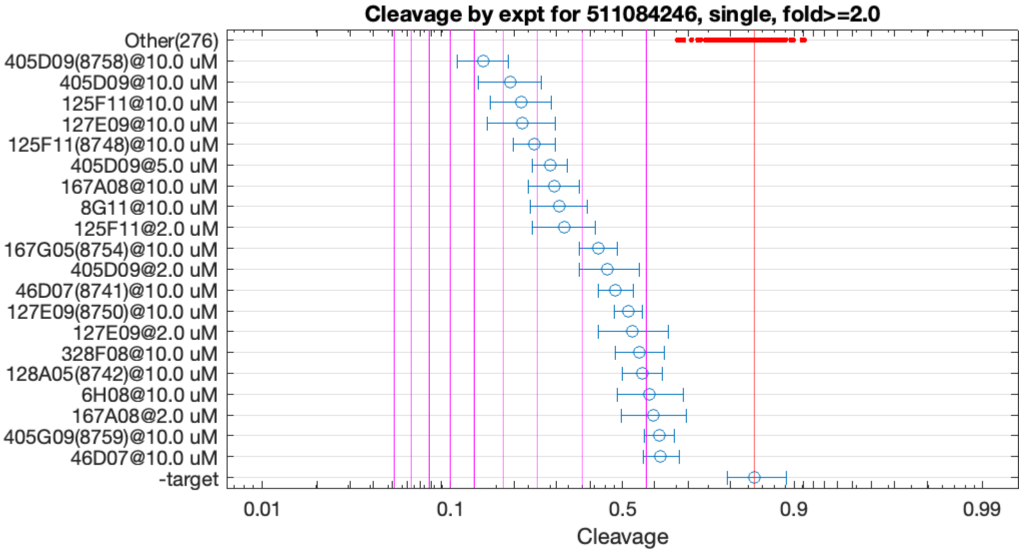


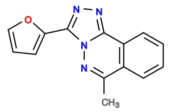

405D09

v>4.3 s=5.6

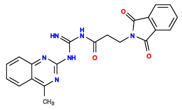

125F11

v>2.0 s=4.4

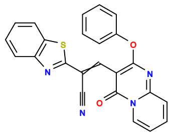

127E09

v>2.4 s=4.3

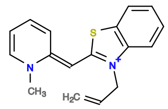

167A08

v>2.4 s=3.6

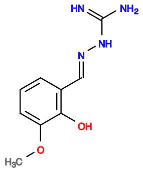

8G11

v>3.5 s=3.5

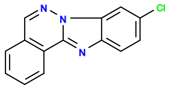

167G05

v>2.0 s=2.7

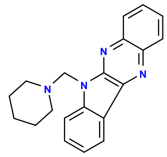

46D07

v>2.2 s=2.4

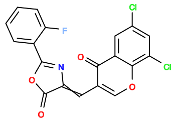

564E04

v>2.4

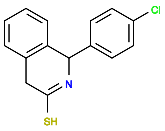

168A06

v>2.3

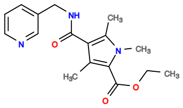

326A05

v>2.3

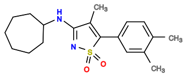

446G09

v>2.3

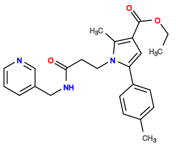

325C10

v>2.3

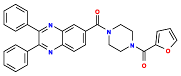

568D05

v>2.2

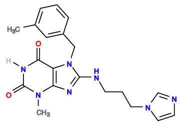

88D02

v>2.2

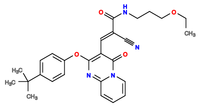

127C11

v>2.2

**511447013 GCTGTC AC TGGAGATGAGTCC GT CTGATGA GCGC TTAGCC GCGT GAAACAGC CTACCTGCCCGGACGAAACAGC**

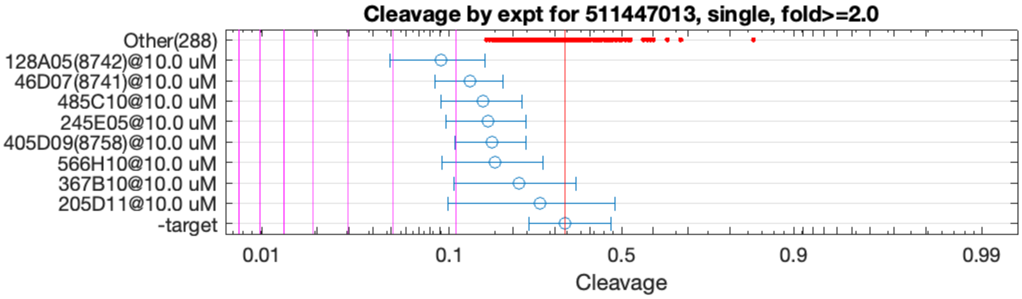


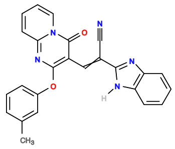

128A05

v>0.6 s=2.3

512101813 GCTGTC ACTGG ATGCATGACCATATTTGGTGTGCCGAGCGGCA CCAGT CTGACGA GTCC GTGGCG GGAC GAAACAGC

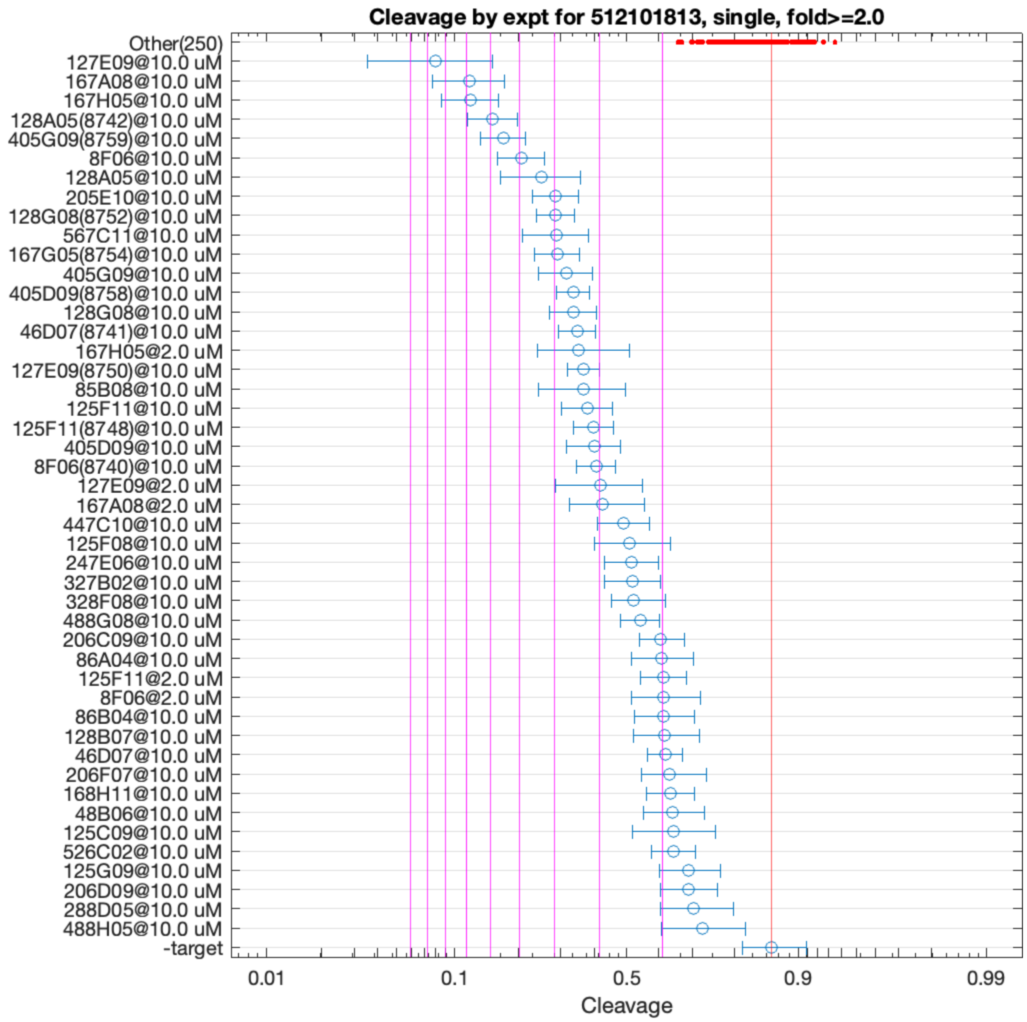


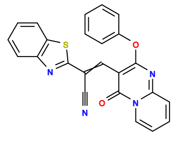

127E09

v>3.5 s=8.7

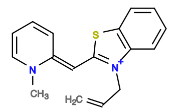

167A08

v>3.0 s=7.0

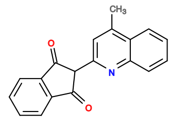

167H05

v>3.5 s=6.9

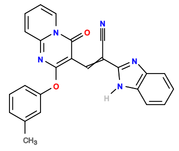

128A05

v>3.1 s=5.9

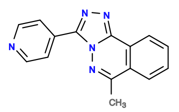

405G09

v>3.9 s=5.6

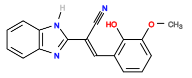

8F06

v>2.5 s=4.9

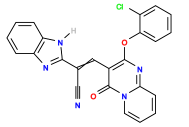

128G08

v>3.3 s=3.9

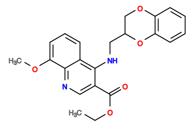

205E10

v>2.8 s=3.9

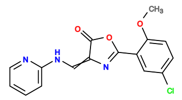

567C11

v>2.7 s=3.9

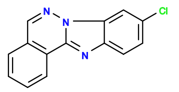

167G05

v>3.0 s=3.9

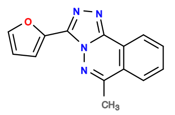

405D09

v>2.6 s=3.5

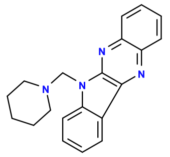

46D07

v>2.6 s=3.5

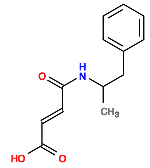

6E06

v>3.4

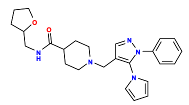

446C03

v>3.4

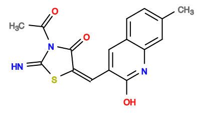

85B08

v>2.9 s=3.3

**512112258 GCTGTC ACTGGA AACGAAGG TCTAGT CTGAAGA GTCC GCCTAAGCGCTGCTCACACAGTGTCTCGTG GGAC GAAACAGC**

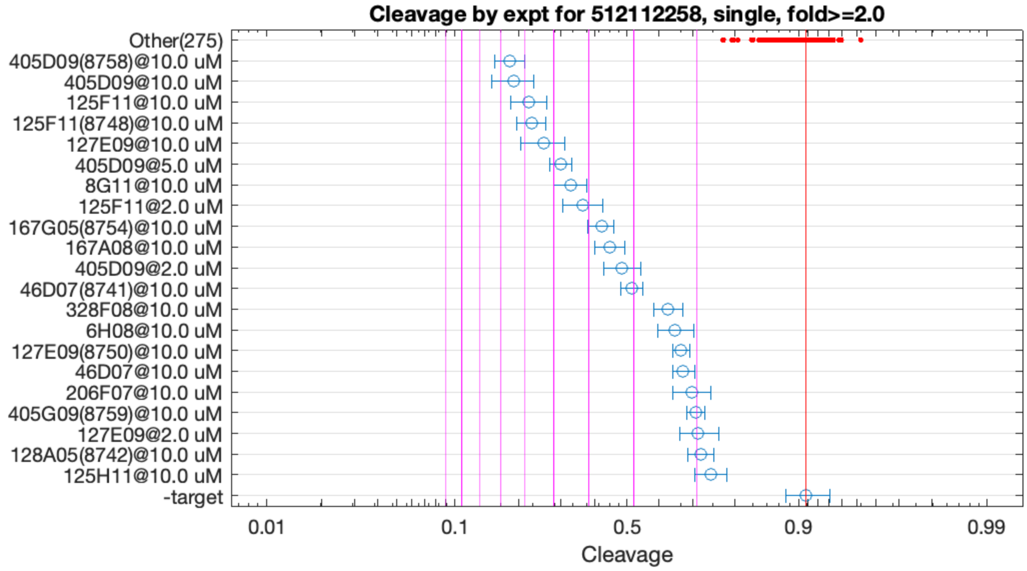


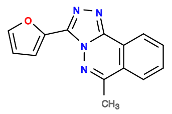

405D09

v>6.4 s=6.7

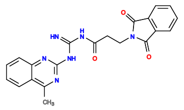

125F11

v>2.7 s=5.8

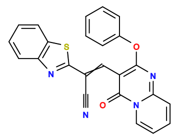

127E09

v>3.0 s=5.3

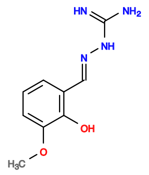

8G11

v>5.1 s=4.5

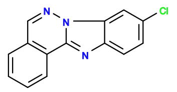

167G05

v>2.8 s=3.6

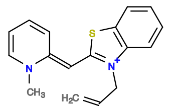

167A08

v>2.9 s=3.5

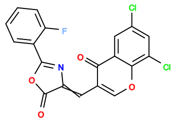

564E04

v>3.2

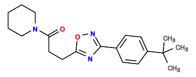

405A02

v>3.1

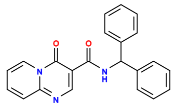

325F04

v>3.1

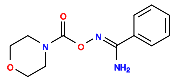

86H09

v>3.0

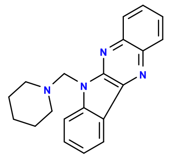

46D07

v>3.1 s=3.0

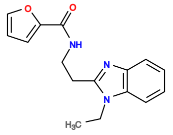

408C11

v>3.0

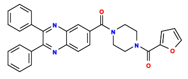

568D05

v>2.9

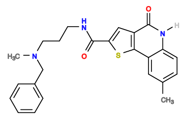

328F08

v>2.5 s=2.4

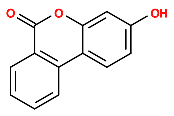

6H08

v>3.8 s=2.3

512117004 GCTGTC ACTGGA GGGGCAAG TCCGGT CTGACGA GTCCT GATAACGAGCCTATTGGCGCACCCGGG GGGAC GAAACAGC

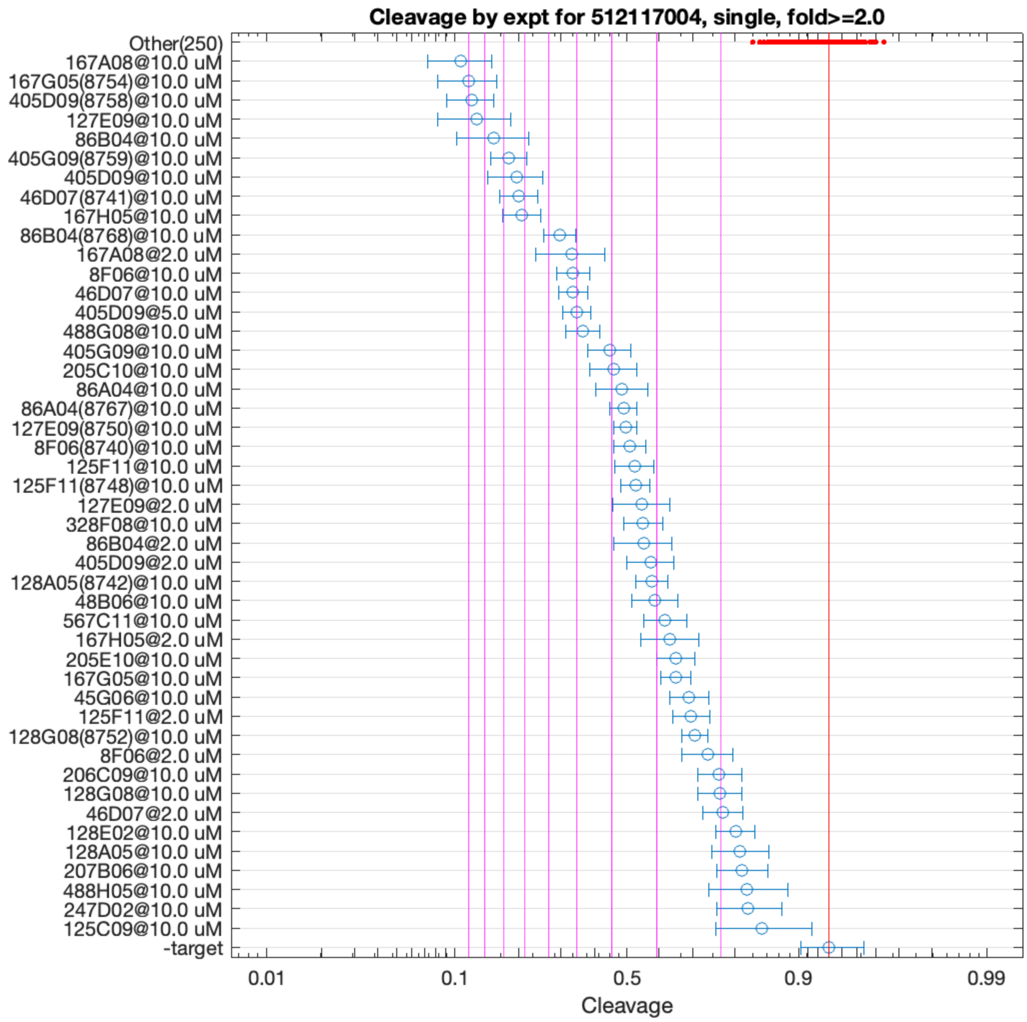


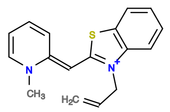

167A08

v>8.0 s=10.2

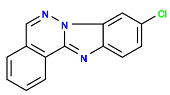

167G05

v>7.5 s=10.1

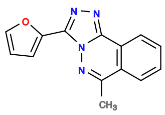

405D09

v>7.4 s=9.6

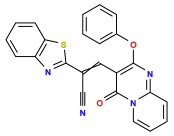

127E09

v>5.1 s=9.6

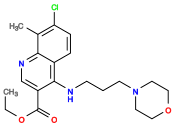

86B04

v>6.3 s=8.5

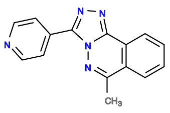

405G09

v>6.4 s=7.9

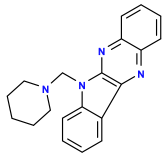

46D07

v>6.6 s=7.3

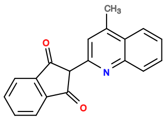

167H05

v>3.8 s=7.1

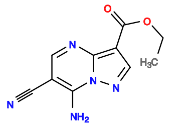

205E05

v>6.1

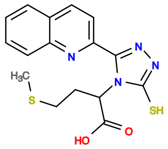

168F07

v>6.0

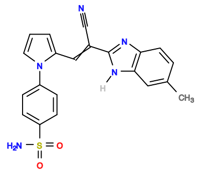

46B07

v>5.8

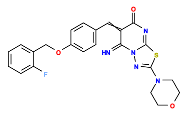

327H06

v>5.8

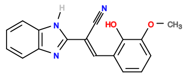

8F06

v>3.9 s=5.2

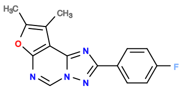

488G08

v>4.2 s=4.8

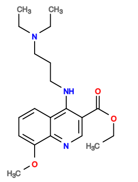

205C10

v>6.4 s=3.9

512119028 GCTGTC ACTGG ATGCATGACCATATTTGGTGTGCCGAGCGGCA CCAGT CTGACGA GTCT GTGGCG GGAC GAAACAGC

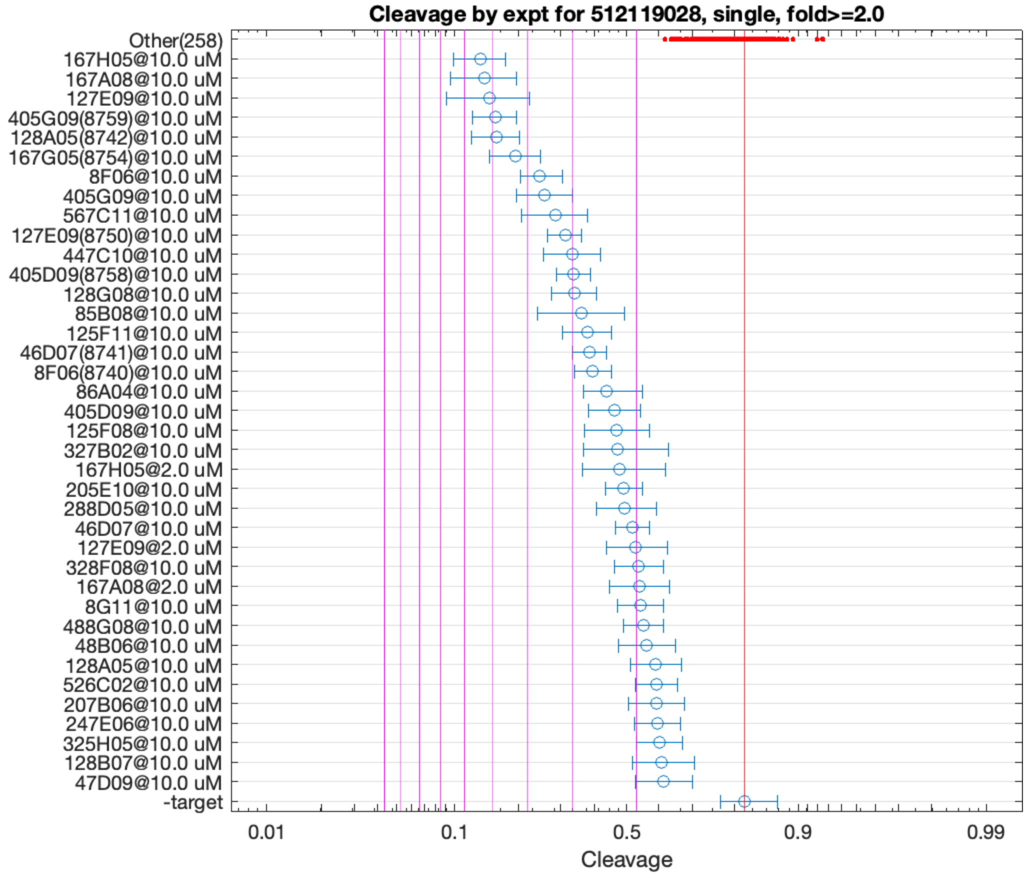


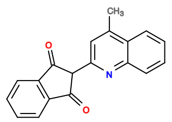

167H05

v>1.7 s=5.5

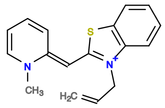

167A08

v>2.9 s=5.3

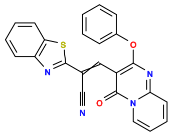

127E09

v>3.0 s=5.1

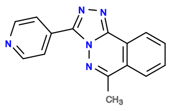

405G09

v>3.6 s=4.9

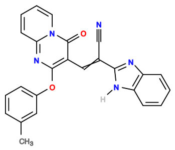

128A05

v>2.4 s=4.9

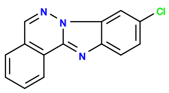

167G05

v>3.0 s=4.3

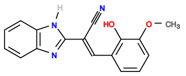

8F06

v>2.3 s=3.7

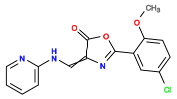

567C11

v>2.5 s=3.4

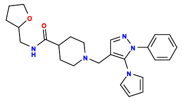

446C03

v>3.1

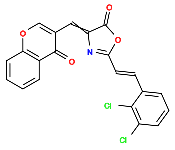

567D05

v>3.1

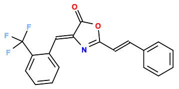

566D10

v>3.0

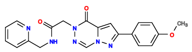

447C10

v>2.9 s=3.0

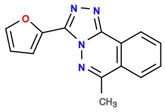

405D09

v>2.4 s=3.0

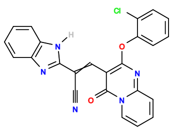

128G08

v>3.3 s=3.0

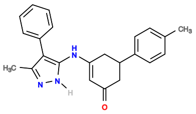

168E02

v>2.9

512119032 GCTGTC ACTGGA ATTATGCCTTTCGAAGGGTTGCCGATCTAA TCTGGT CTGAAGA GTCT GTGGCG GGAC GAAACAGC

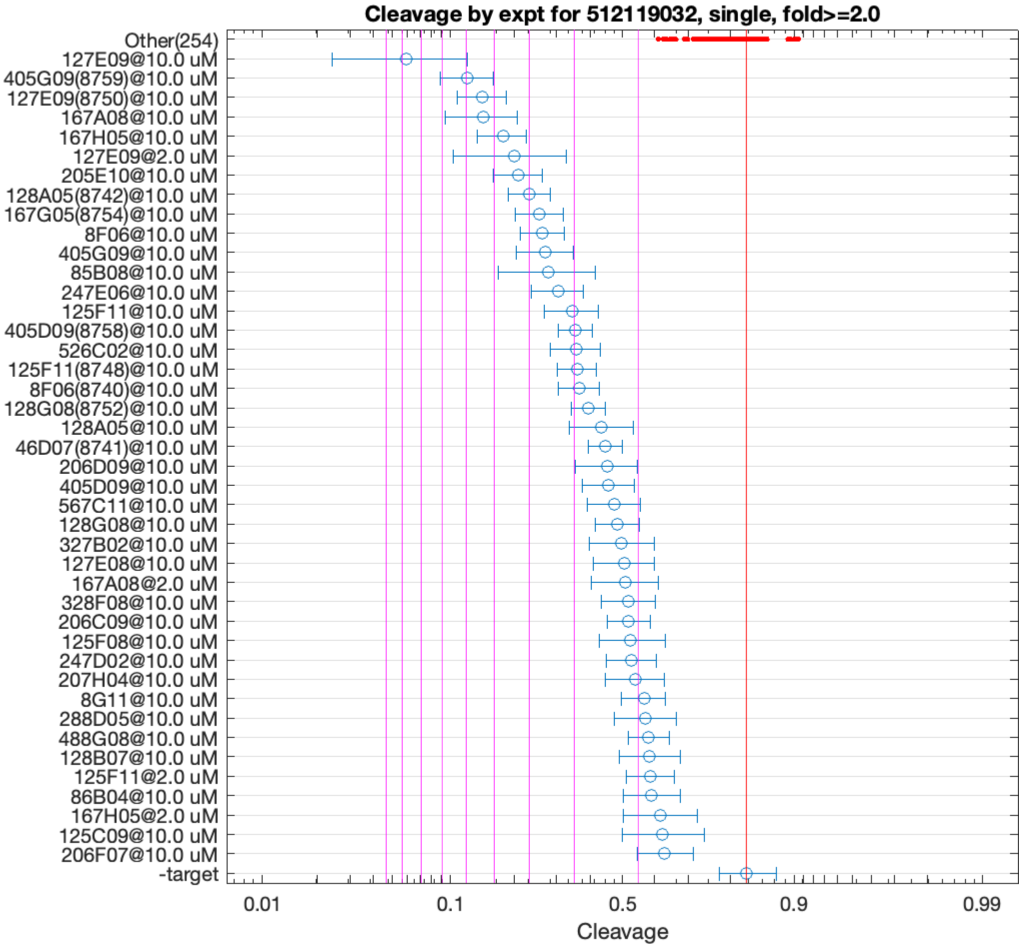


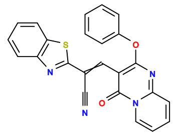

127E09

v>3.5 s=8.7

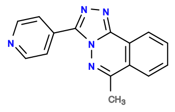

405G09

v>3.6 s=6.0

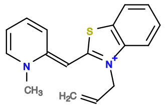

167A08

v>2.7 s=5.3

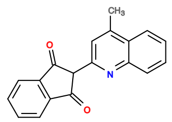

167H05

v>2.7 s=4.7

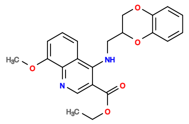

205E10

v>2.8 s=4.3

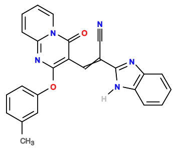

128A05

v>2.4 s=4.0

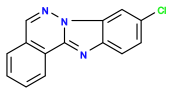

167G05

v>2.9 s=3.8

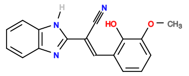

8F06

v>2.3 s=3.7

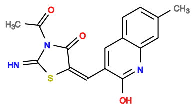

85B08

v>2.7 s=3.5

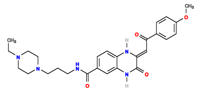

247E06

v>2.6 s=3.3

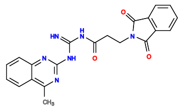

125F11

v>2.2 s=3.1

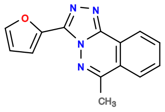

405D09

v>2.5 s=3.0

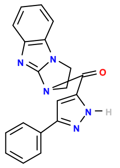

526C02

v>2.9 s=3.0

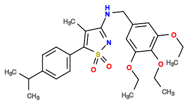

447C04

v>2.9

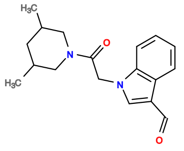

606E08

v>2.9

512119326 GCTGTC ACTGG ATGCATGACCATATTTGGTGTGCCGAGCGGCA CCAGT CTGACGA GTTC GTGGCG GGAC GAAACAGC

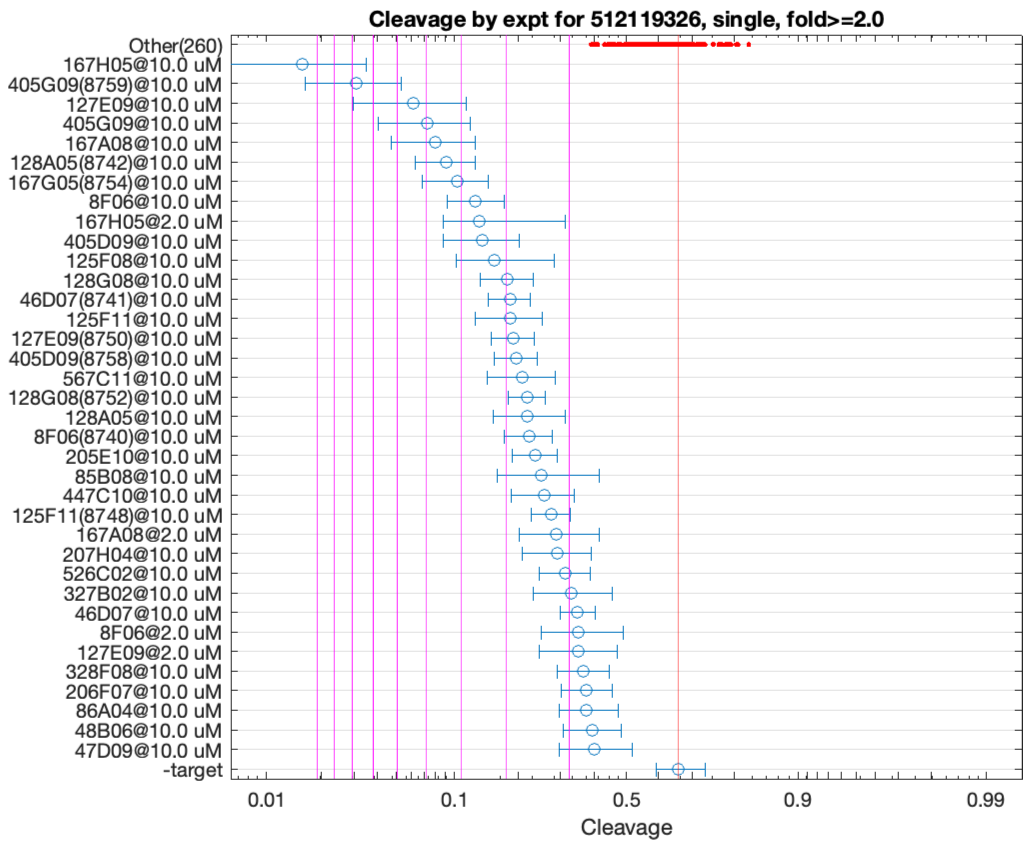


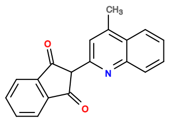

167H05

v>2.3 s=11.3

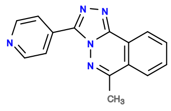

405G09

v>3.2 s=7.7

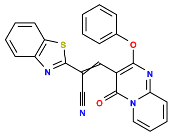

127E09

v>3.0 s=5.4

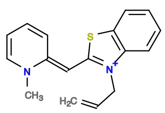

167A08

v>2.9 s=4.8

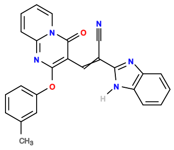

128A05

v>2.5 s=4.4

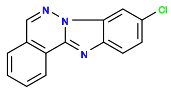

167G05

v>3.1 s=4.2

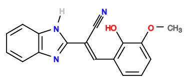

8F06

v>2.3 s=3.6

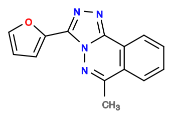

405D09

v>2.4 s=3.5

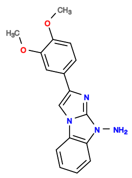

125F08

v>2.4 s=3.2

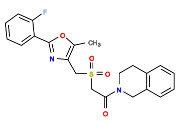

286H08

v>3.0

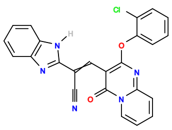

128G08

v>3.1 s=3.0

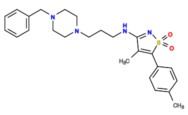

446E08

v>2.9

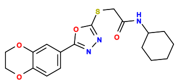

246A07

v>2.9

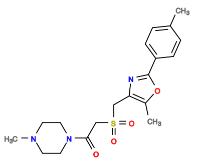

285A10

v>2.9

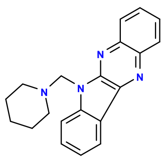

46D07

v>2.6 s=2.9

**512158904 GCTGTC ACTGGAG AAACTGCACATCGTGTAGTGGAGGCAAC TTCCGGT CTGAAGA GTCC TAGAGCCT GGAC GAAACAGC**

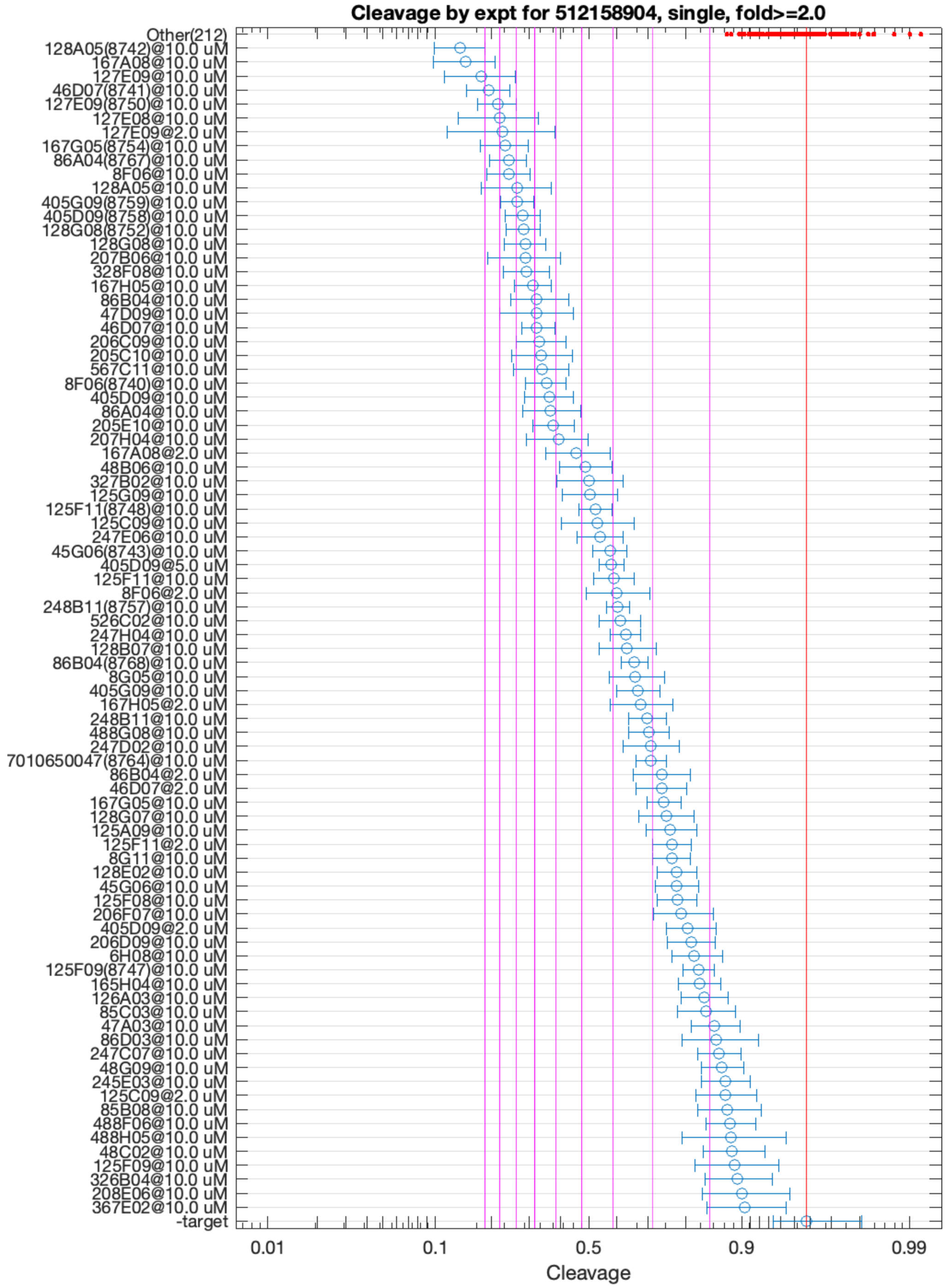


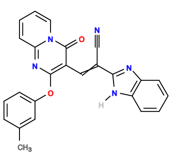

128A05

v>6.9 s=11.8

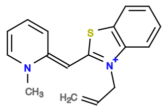

167A08

v>7.4 s=11.5

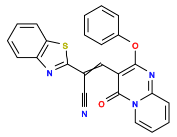

127E09

v>8.6 s=10.4

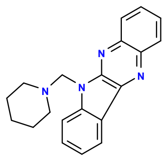

46D07

v>7.1 s=9.8

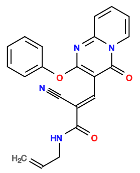

127E08

v>8.8 s=8.9

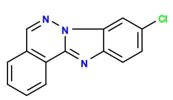

167G05

v>7.4 s=8.8

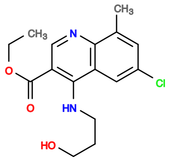

86A04

v>6.0 s=8.7

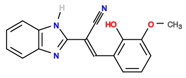

8F06

v>6.1 s=8.5

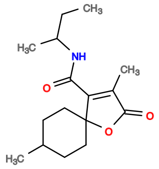

286D11

v>8.3

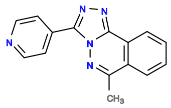

405G09

v>6.3 s=8.1

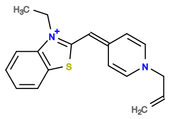

167E07

v>8.1

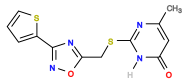

168G03

v>8.0

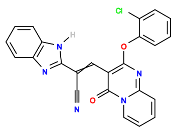

128G08

v>7.7 s=7.8

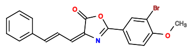

567A09

v>7.7

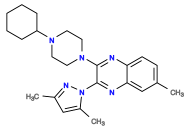

248E02

v>7.7

**512213078 GCTGTC ACTGGA TGCATGACCTATTTGGTGTGCTGAGCTGC TCTGGT CTGACGA GTCC GTGGCG GGAC GAAACAGC**

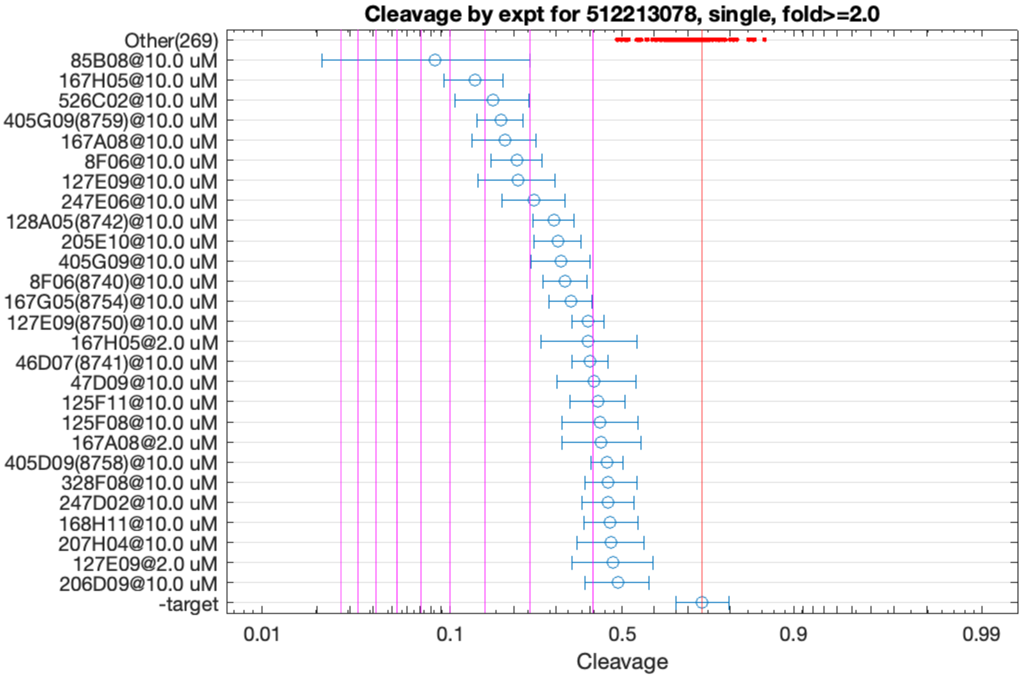


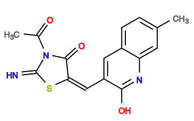

85B08

v>2.7 s=5.6

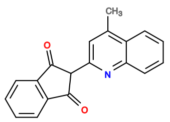

167H05

v>2.1 s=4.3

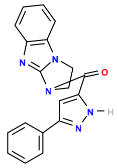

526C02

v>2.7 s=3.8

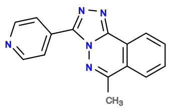

405G09

v>2.5 s=3.6

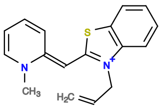

167A08

v>2.2 s=3.5

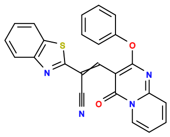

127E09

v>2.2 s=3.3

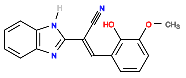

8F06

v>1.9 s=3.2

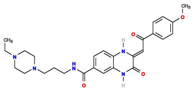

247E06

v>2.2 s=2.9

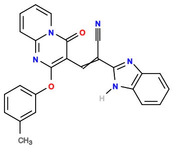

128A05

v>1.9 s=2.6

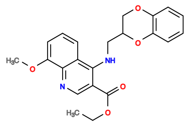

205E10

v>2.0 s=2.5

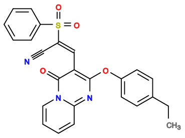

128G06

v>2.5

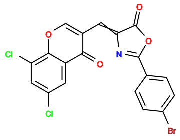

564H11

v>2.5

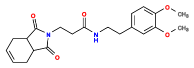

205G07

v>2.5

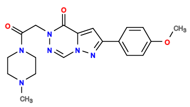

447E09

v>2.4

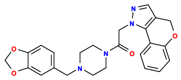

328G11

v>2.4

**512213338 GCTGTC ACTGGA TGGAGGCT TCTAGT CTGACGA GTCCTG CAACAACCCAGTTGCACGCAAGCC CGGGAC GAAACAGC**

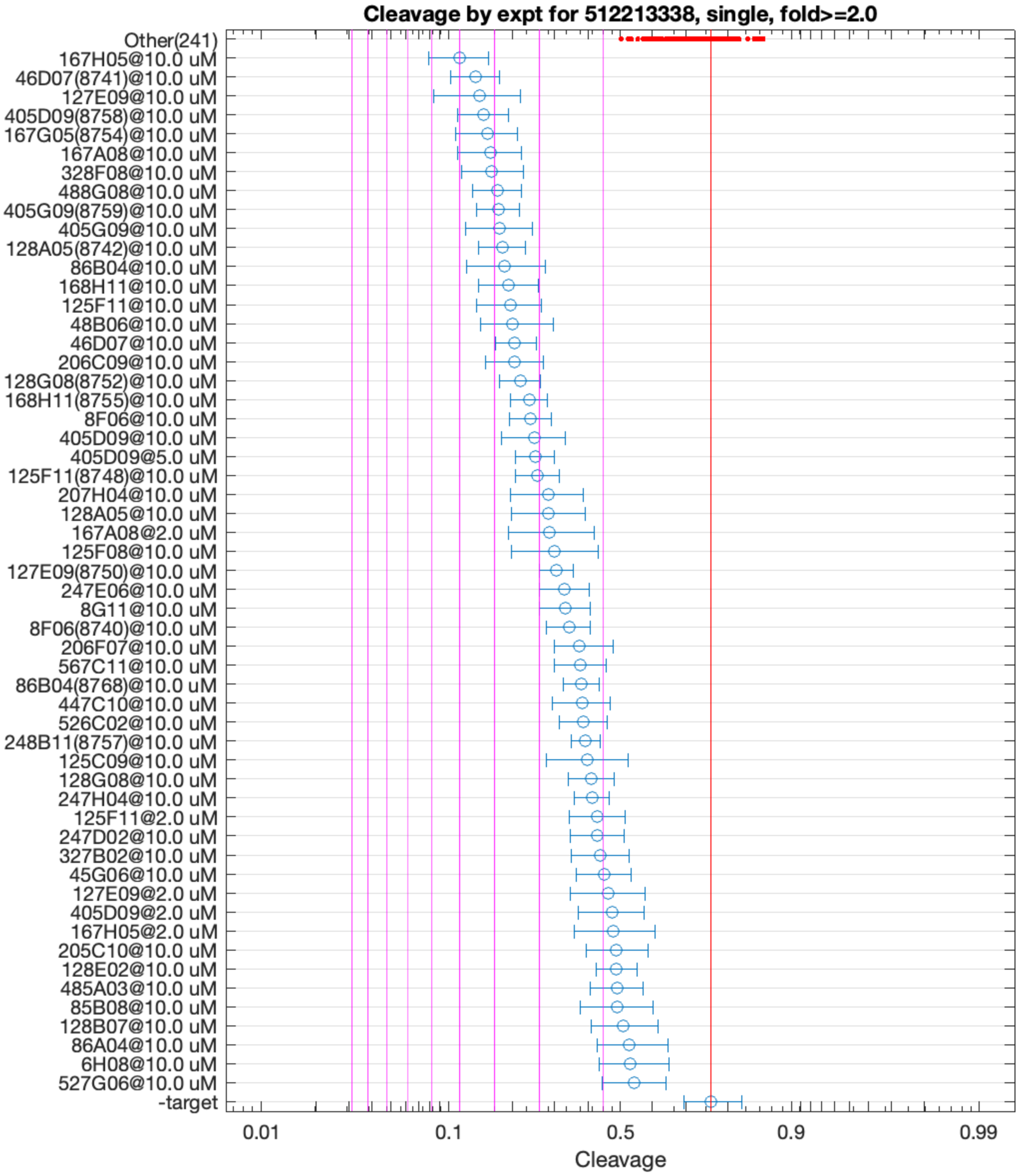


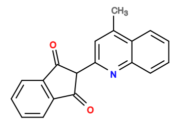

167H05

v>3.1 s=5.1

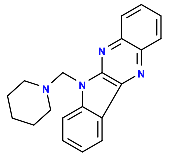

46D07

v>3.1 s=4.5

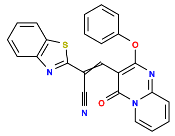

127E09

v>3.0 s=4.4

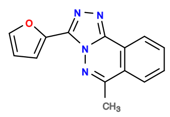

405D09

v>3.4 s=4.4

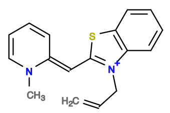

167A08

v>3.1 s=4.1

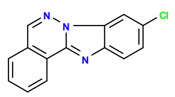

167G05

v>3.3 s=4.1

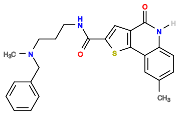

328F08

v>3.4 s=4.0

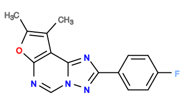

488G08

v>3.0 s=4.0

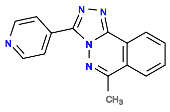

405G09

v>3.0 s=4.0

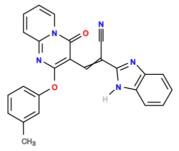

128A05

v>2.8 s=3.8

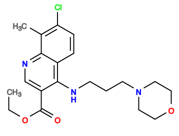

86B04

v>3.2 s=3.8

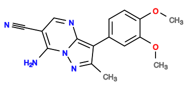

168H11

v>3.2 s=3.7

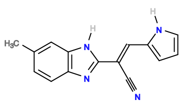

48B06

v>3.3 s=3.6

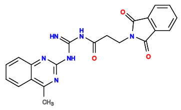

125F11

v>2.7 s=3.6

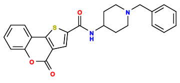

206C09

v>2.9 s=3.5

**512228786 GCTGTC ACTGGA AACGAAGG TCTAGT CTGAAGA GTCC GCCTAAGCGCTGCTCACACAGTGACTCGTG GGAC GAAACAGC**

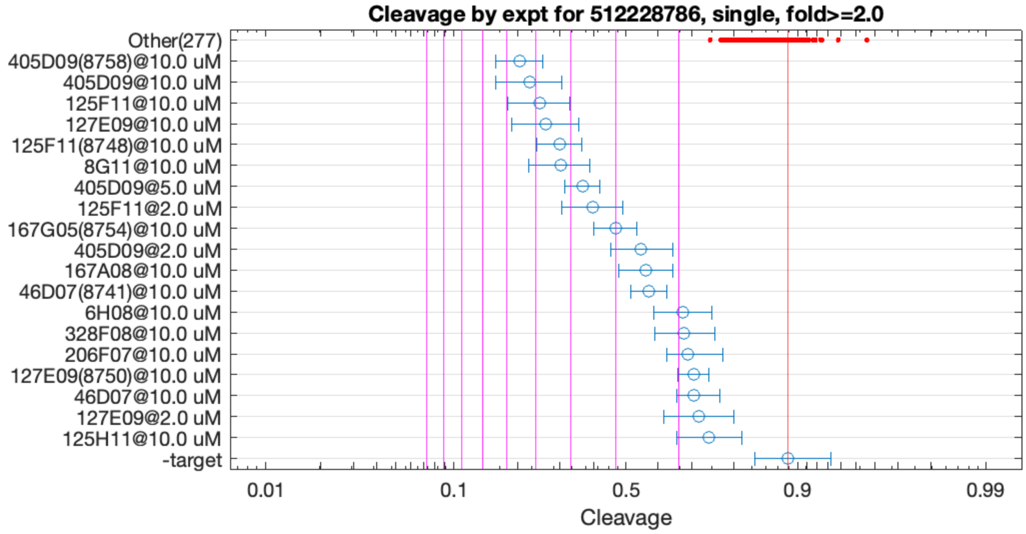


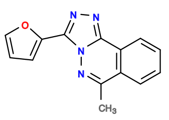

405D09

v>4.7 s=5.5

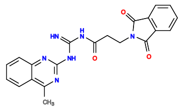

125F11

v>2.0 s=4.8

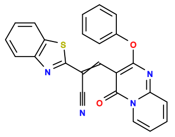

127E09

v>2.2 s=4.6

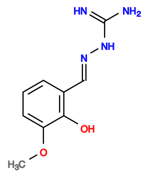

8G11

v>3.7 s=4.2

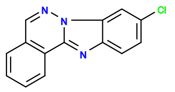

167G05

v>2.1 s=3.0

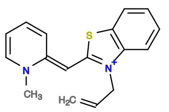

167A08

v>2.0 s=2.5

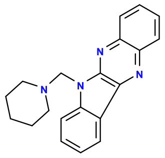

46D07

v>2.3 s=2.4

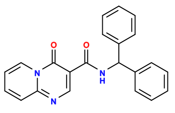

325F04

v>2.2

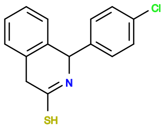

168A06

v>2.2

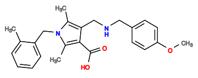

365C08

v>2.2

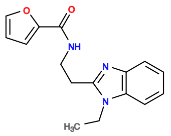

408C11

v>2.2

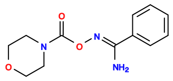

86H09

v>2.1

**512257380 GCTGTC ACTG GATGGGAAA CAGT CTGATGA GTCC TGCGATGCCTGATACACATGCCTCTCCGC GGAC GAAACAGC**

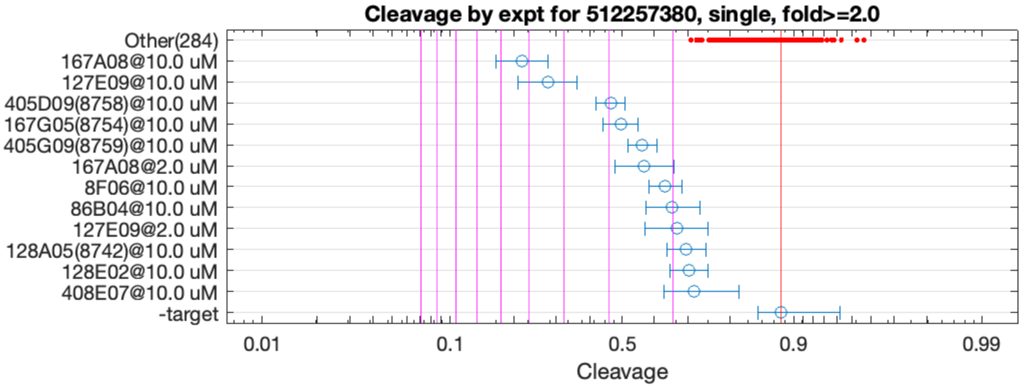


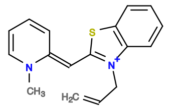

167A08

v>2.9 s=5.3

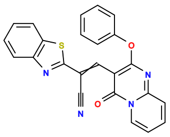

127E09

v>2.4 s=4.4

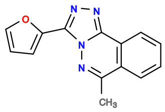

405D09

v>2.4 s=3.0

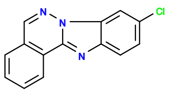

167G05

v>2.4 s=2.8

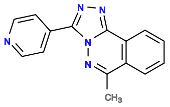

405G09

v>2.3 s=2.4

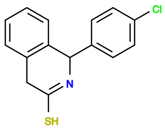

168A06

v>2.3

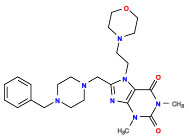

166C04

v>2.2

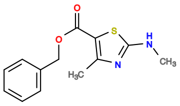

86F10

v>2.2

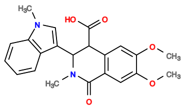

207A03

v>2.2

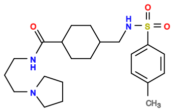

606A04

v>2.2

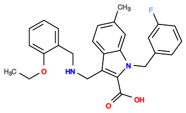

527D04

v>2.2

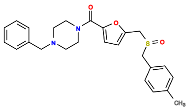

487A06

v>2.1

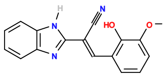

8F06

v>1.9 s=2.1

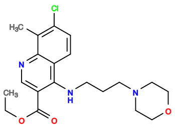

86B04

v>2.3 s=2.0

**512293240 GCTGTC ACTGGA GAACACG TTCAGT CTGATGA GTCC ACTGCCTCTGCCCTATCTGCAAGCGTACCG GGAC GAAACAGC**

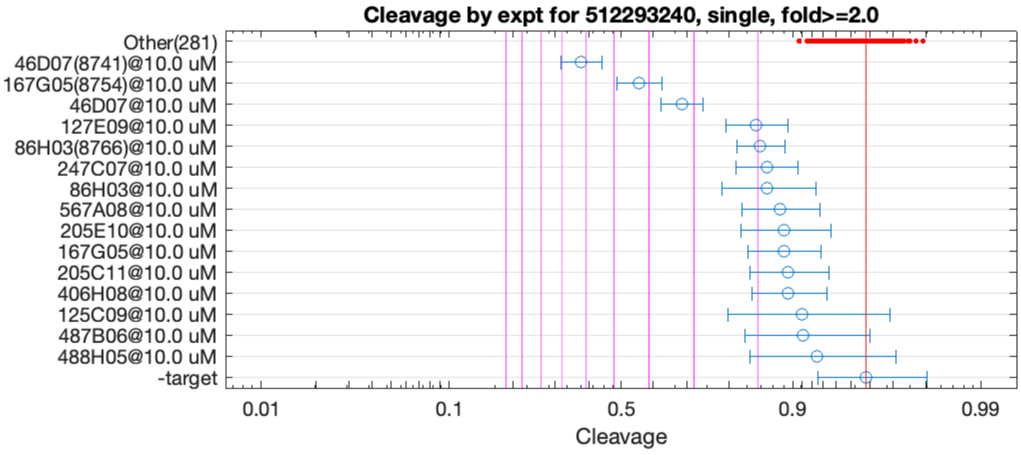


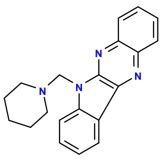

46D07

v>2.1 s=6.2

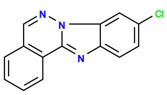

167G05

v>1.2 s=4.2

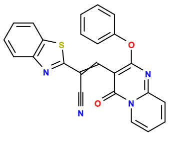

127E09

v>1.0 s=2.0

**512295690 GCTGTC ACTGGA GGCAGGA TCTGGT CTGATGA GTCC TCGATGTGGTCGCAATTCCGCATACGTGCC GGAC GAAACAGC**

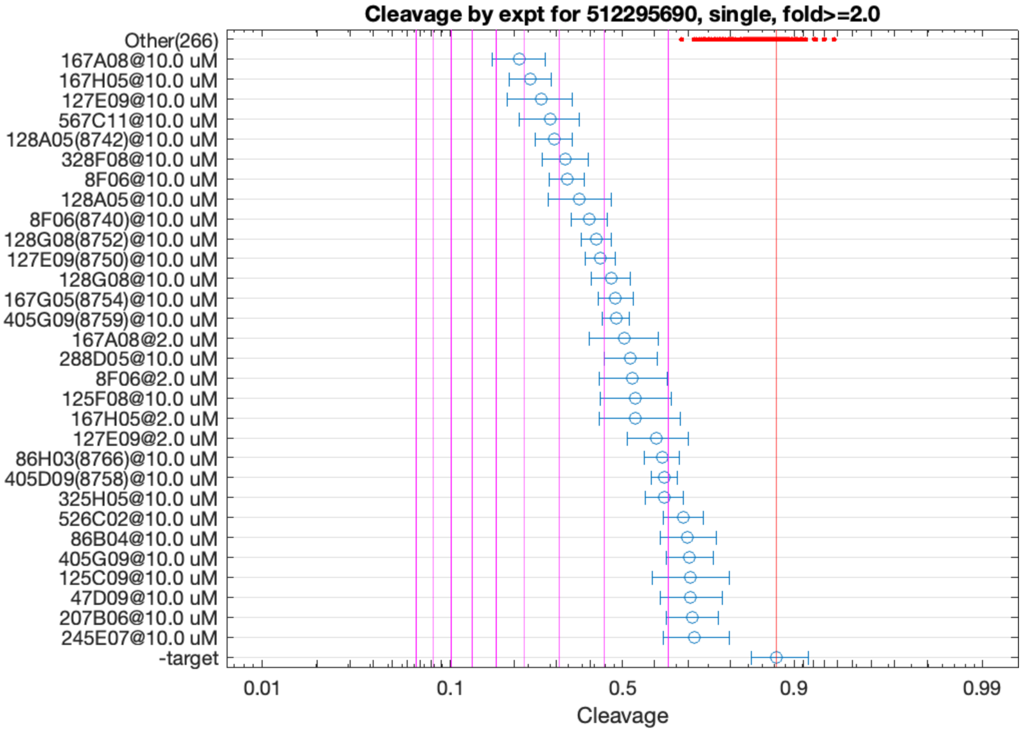


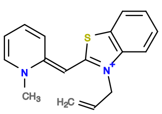

167A08

v>3.4 s=5.3

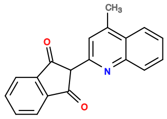

167H05

v>3.3 s=4.8

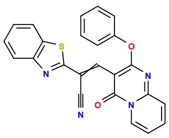

127E09

v>2.9 s=4.5

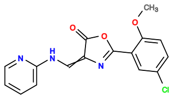

567C11

v>3.4 s=4.3

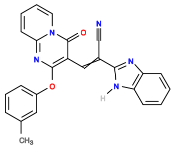

128A05

v>2.8 s=4.1

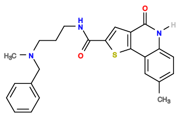

328F08

v>3.4 s=3.8

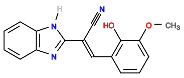

8F06

v>3.0 s=3.8

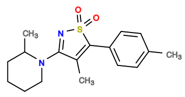

446H05

v>3.7

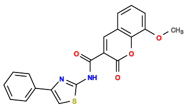

568H09

v>3.7

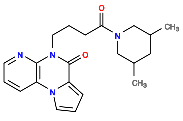

485H06

v>3.6

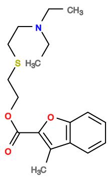

48G05

v>3.5

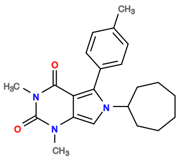

408D06

v>3.5

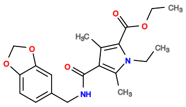

326G06

v>3.5

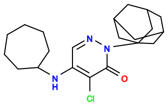

247D10

v>3.5

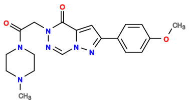

447E09

v>3.5

512298529 GCTGTC ACTGGAT CGGACA ATCTAGT CTGAAGA GTCC TGCGATGCCTGATACACATGCCTCTCCGC GGAC GAAACAGC

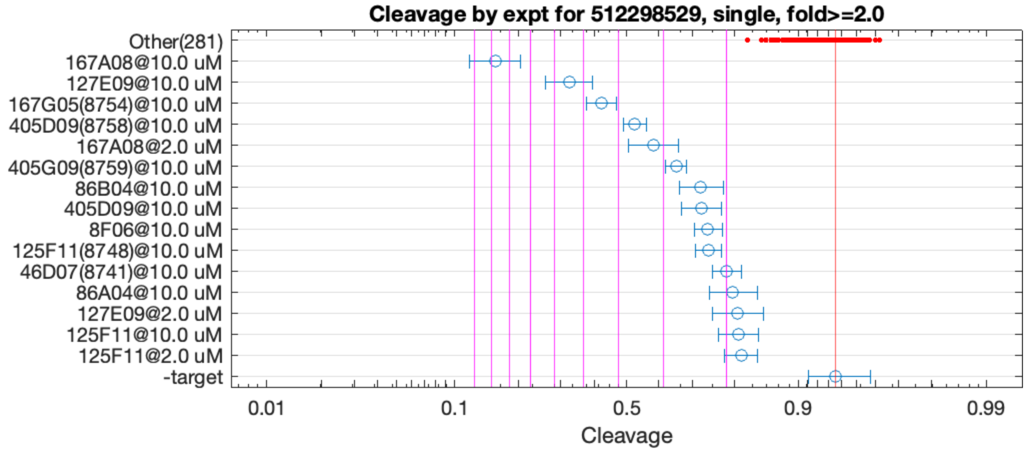


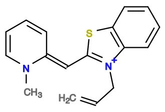

167A08

v>3.0 s=8.7

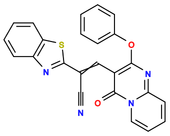

127E09

v>2.0 s=5.4

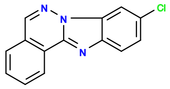

167G05

v>2.2 s=4.5

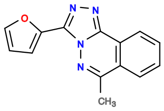

405D09

v>2.2 s=3.5

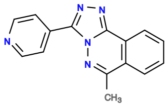

405G09

v>2.1 s=2.8

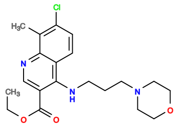

86B04

v>1.7 s=2.4

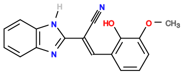

8F06

v>1.5 s=2.2

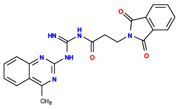

125F11

v>1.5 s=2.2

**512299935 GCTGTC ACTGGA TGCATGACCATATTTGGTGTGCTGAGCTGC TCCGGT CTGATGA GTCTT TGGC AGGAC GAAACAGC**

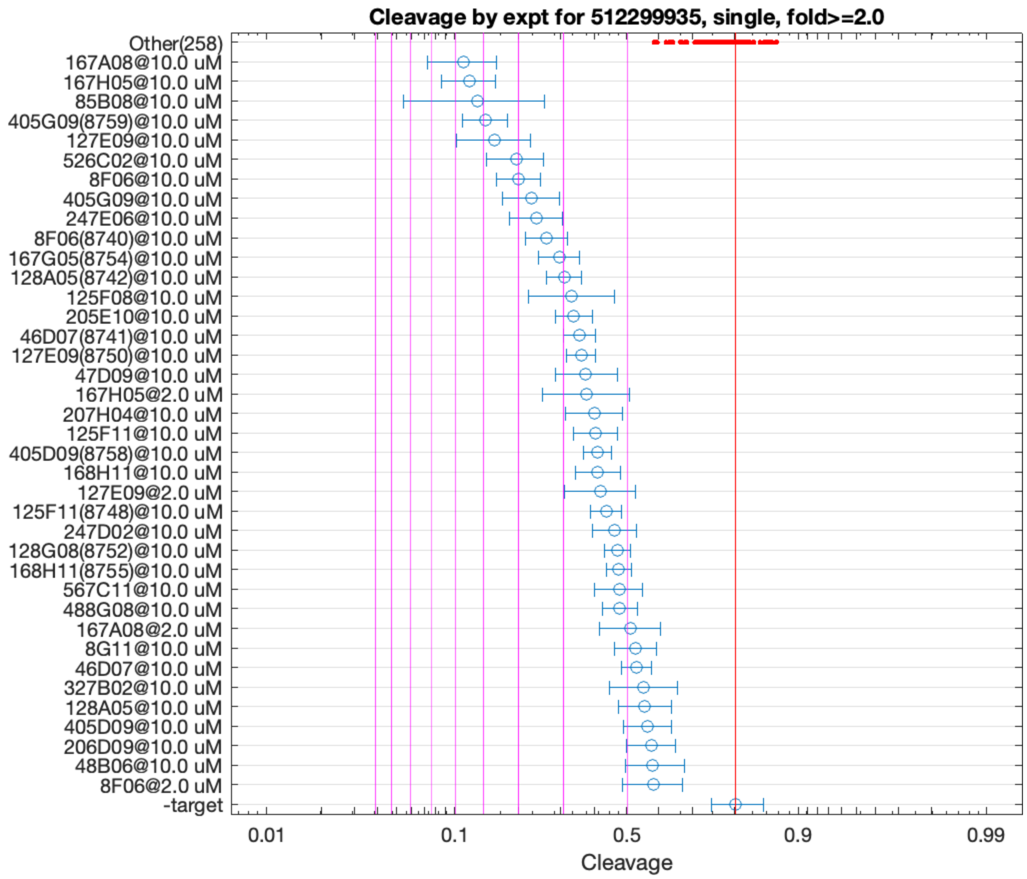


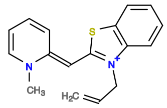

167A08

v>3.1 s=5.8

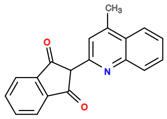

167H05

v>2.8 s=5.5

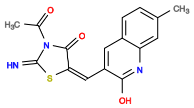

85B08

v>3.7 s=5.2

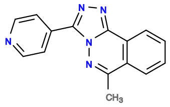

405G09

v>3.5 s=5.0

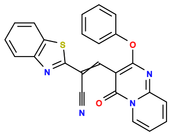

127E09

v>2.8 s=4.6

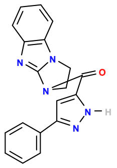

526C02

v>3.7 s=4.1

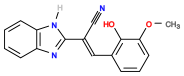

8F06

v>2.5 s=4.0

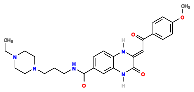

247E06

v>2.8 s=3.5

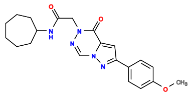

447E10

v>3.3

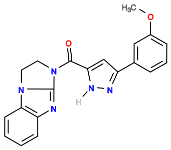

525D11

v>3.3

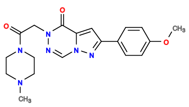

447E09

v>3.3

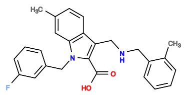

527H03

v>3.3

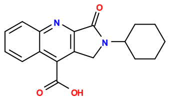

208A06

v>3.3

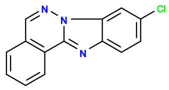

167G05

v>2.9 s=3.1

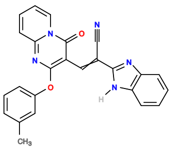

128A05

v>2.4 s=3.0

512299937 GCTGTC ACTGG ATGCATGACCATATTTTGGTGTGCCGAGCGGCA CCAGT CTGATGA GTTC GTGGCG GGAC GAAACAGC

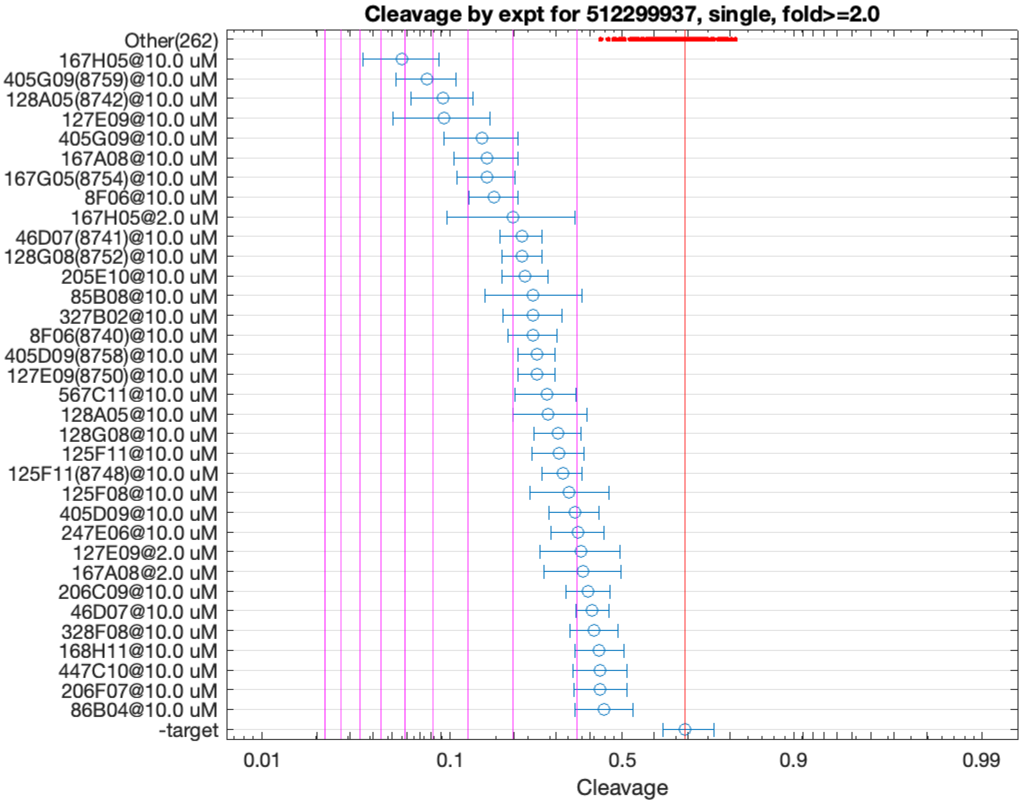


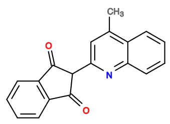

167H05

v>3.1 s=6.0

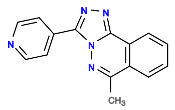

405G09

v>3.1 s=5.2

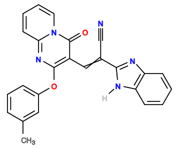

128A05

v>2.6 s=4.7

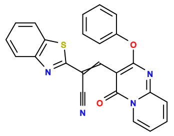

127E09

v>2.9 s=4.7

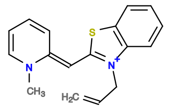

167A08

v>2.3 s=3.5

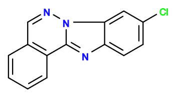

167G05

v>2.6 s=3.5

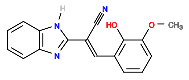

8F06

v>2.0 s=3.4

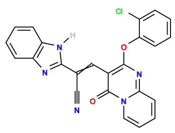

128G08

v>2.8 s=2.9

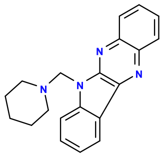

46D07

v>2.1 s=2.8

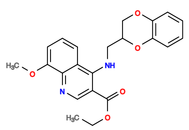

205E10

v>2.4 s=2.8

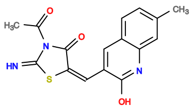

85B08

v>2.3 s=2.7

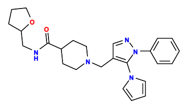

446C03

v>2.7

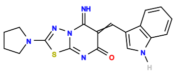

327B02

v>2.7 s=2.7

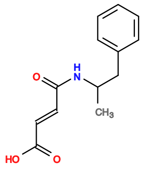

6E06

v>2.6

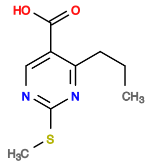

488B10

v>2.6

**512329455 GCTGTC ACTGGA GTTAAGAGTGGTTGTTGCTACTGTGCCGATCTAA TCTGGT CTGACGA GTCC GTGGCG GGAC GAAACAGC**

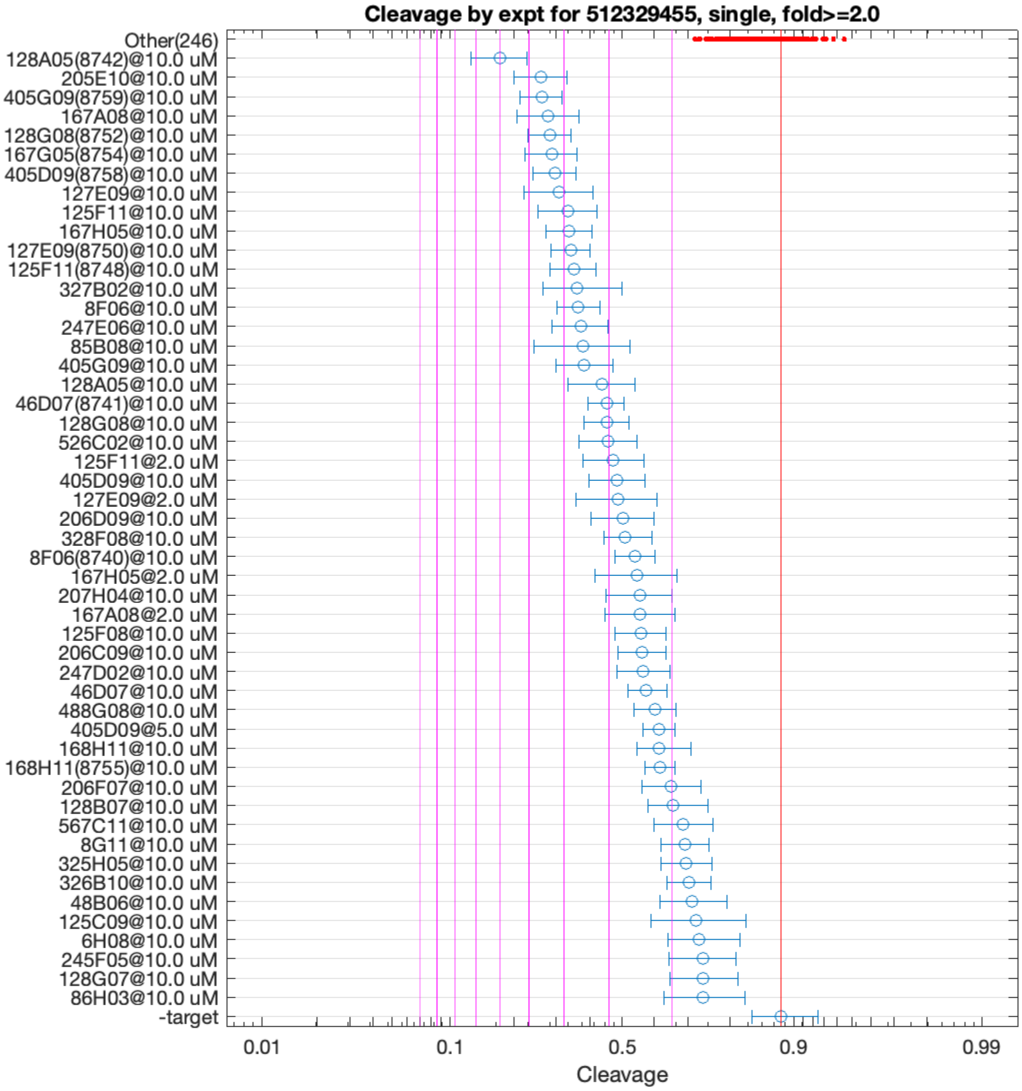


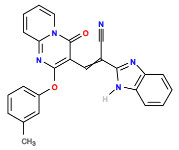

128A05

v>2.9 s=6.0

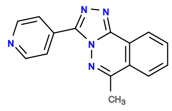

405G09

v>3.1 s=4.7

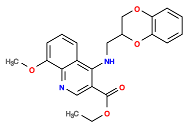

205E10

v>3.5 s=4.5

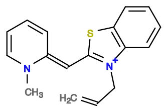

167A08

v>2.8 s=4.4

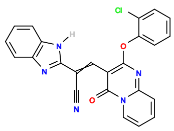

128G08

v>3.4 s=4.4

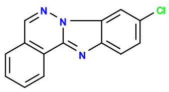

167G05

v>3.0 s=4.2

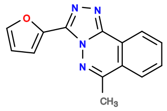

405D09

v>2.7 s=4.2

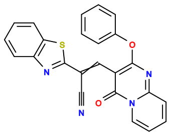

127E09

v>2.9 s=4.1

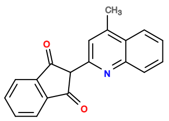

167H05

v>2.9 s=3.9

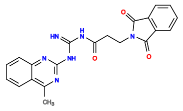

125F11

v>2.5 s=3.9

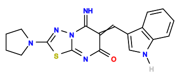

327B02

v>3.2 s=3.7

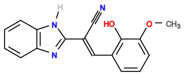

8F06

v>2.5 s=3.7

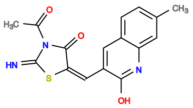

85B08

v>2.8 s=3.6

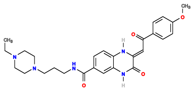

247E06

v>2.5 s=3.6

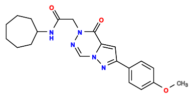

447E10

v>3.5

**513125001 GCTGTC ACTGGA CATTTCTGGGCATTTGGGTTGAATTGGCGAA TCCGGT CTGACGA GTCCTGTA GTCGGTTACCGTGAAGCTCGGG TGTGGGAC GAAACAGC**

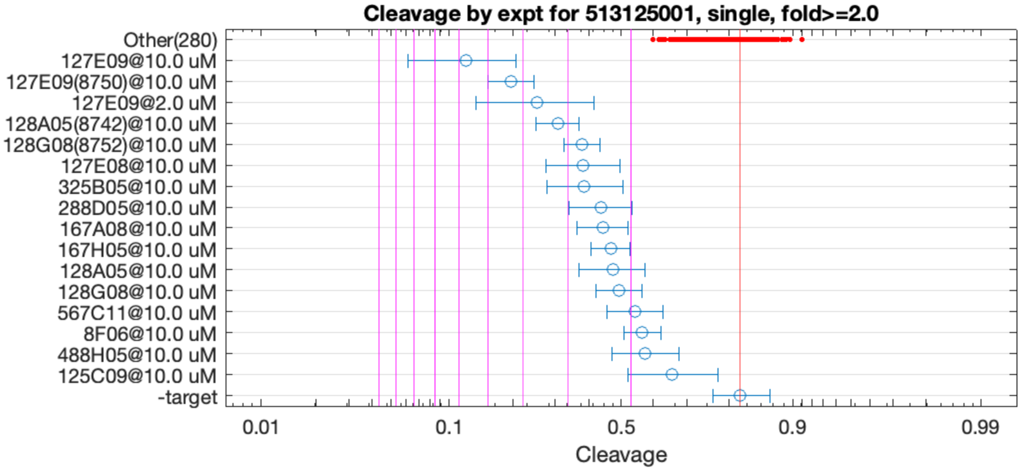


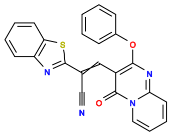

127E09

v>3.3 s=5.8

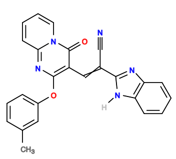

128A05

v>1.7 s=3.2

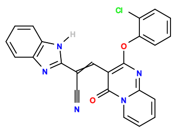

128G08

v>1.9 s=2.8

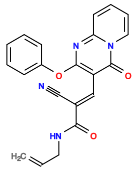

127E08

v>1.7 s=2.7

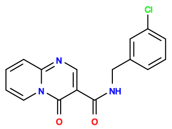

325B05

v>1.4 s=2.7

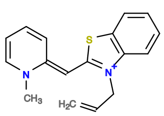

167A08

v>1.4 s=2.5

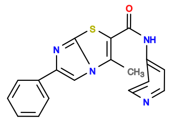

288D05

v>1.8 s=2.4

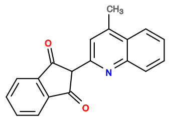

167H05

v>1.3 s=2.2

**513135887 GCTGTC ACTGGA AACCTGCGGTTGGCAAAAGAGAGACACCTC TCCGGT CTGACGA GTCC GATGTGTCTGCGAATGTCTGCGTGTTGGTG GGAC GAAACAGC**

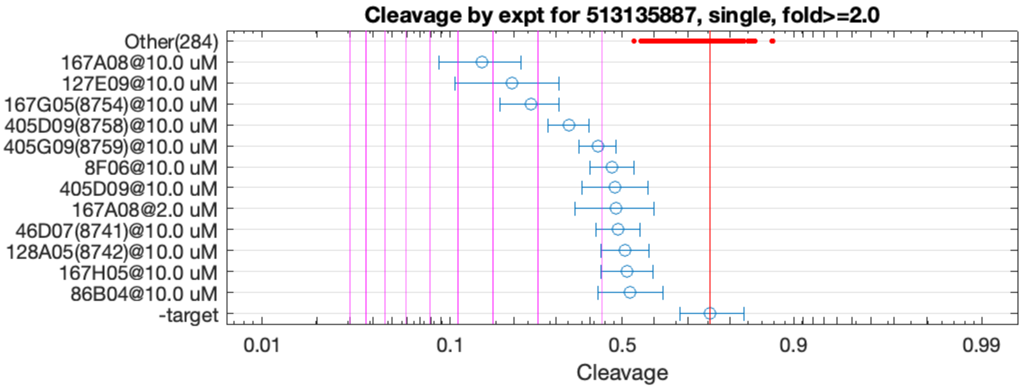


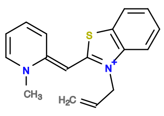

167A08

v>2.1 s=4.2

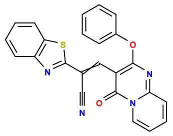

127E09

v>1.8 s=3.5

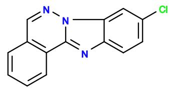

167G05

v>1.8 s=3.2

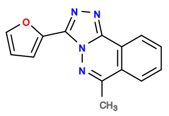

405D09

v>1.8 s=2.5

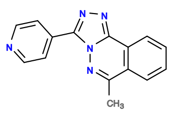

405G09

v>1.7 s=2.0

**513140430 GCTGTC ACTGGAG CCTAGCGGCACTACAACCAACGAACACCT CTCCGGT CTGACGA GTCC GATGTGTCTGCGTGTGGTG GGAC GAAACAGC**

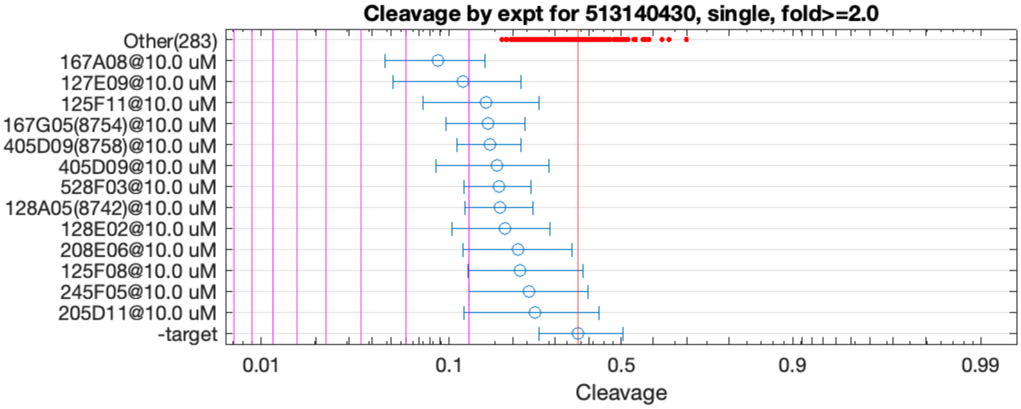


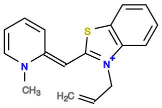

167A08

v>1.4 s=2.4

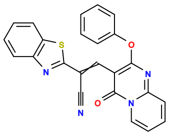

127E09

v>1.2 s=2.0

**513339614 GCTGTC ACTGGA TTTGGGAAACAACGAAACGCCTCAATACAT TCCGGT CTGACGA GTC GTTT GAC GAAACAGC GCGGGACGAAACAGC**

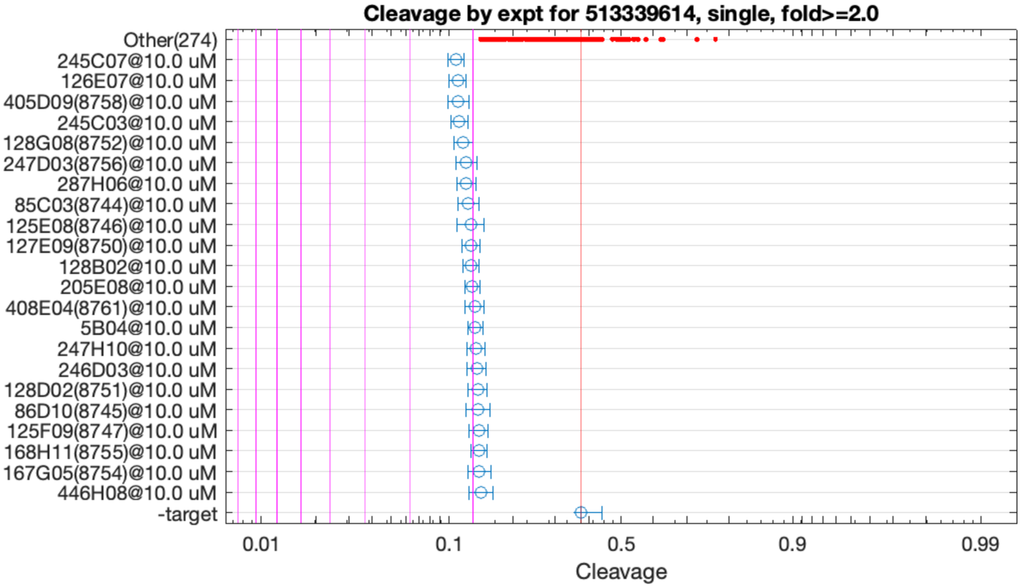


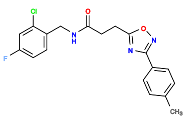

245C07

v>1.0 s=2.2

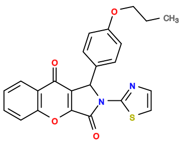

126E07

v>1.1 s=2.2

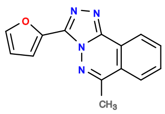

405D09

v>1.0 s=2.2

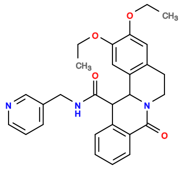

245C03

v>1.1 s=2.2

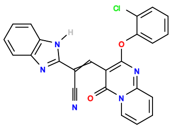

128G08

v>1.1 s=2.1

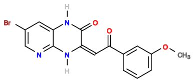

247D03

v>1.0 s=2.1

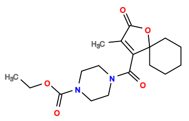

287H06

v>1.0 s=2.1

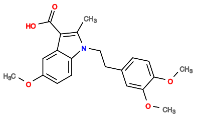

85C03

v>0.9 s=2.0

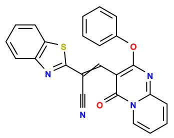

127E09

v>1.0 s=2.0

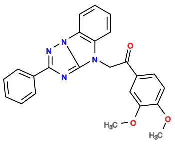

125E08

v>1.1 s=2.0

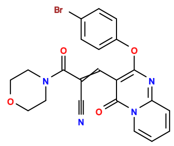

128B02

v>1.0 s=2.0

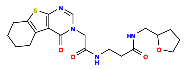

205E08

v>1.0 s=2.0

**513400071 GCTGTC ACTGGA TGGAGGCT TTCAGT CTGACGA GTCCTG CAACAACCCAATTGCACGCAAGCC CGGGAC GAAACAGC**

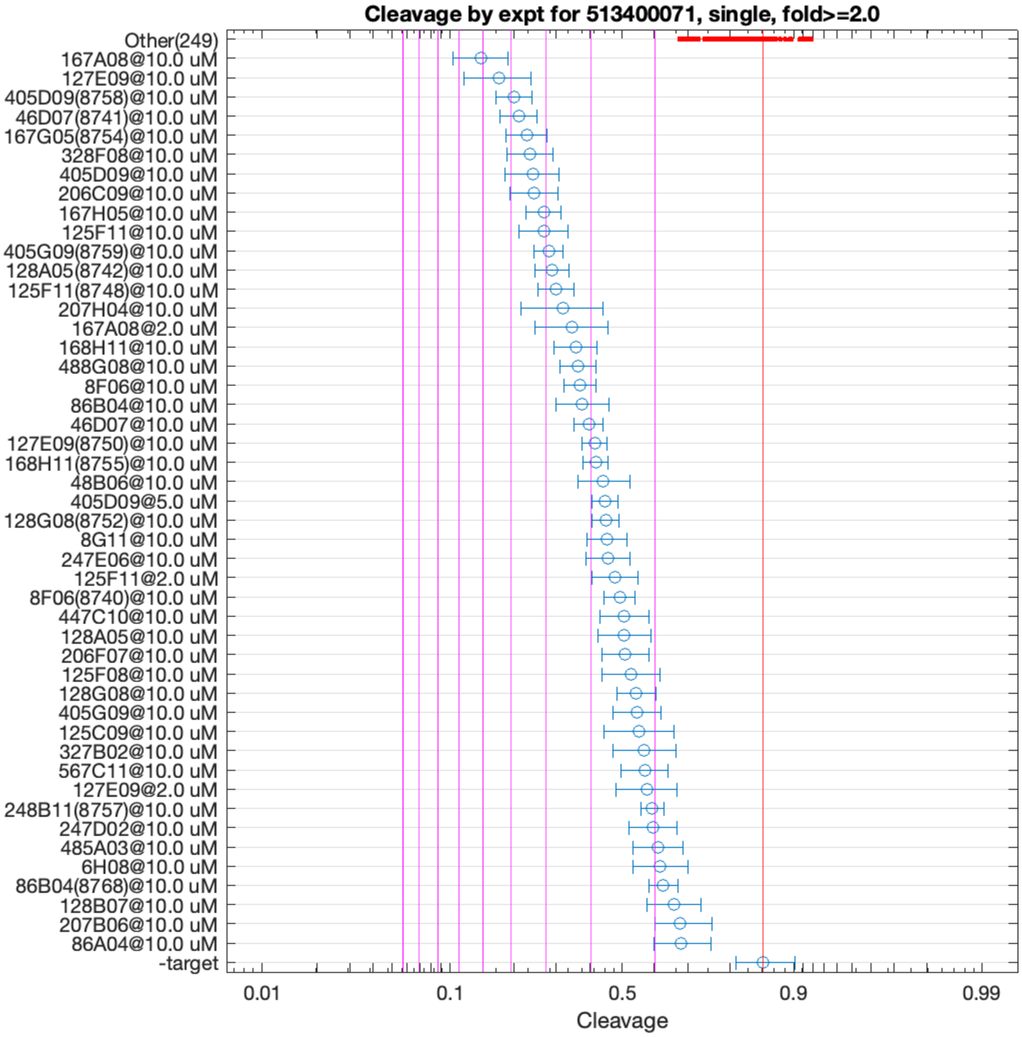


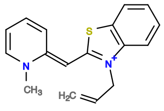

167A08

v>3.9 s=6.2

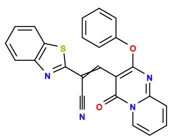

127E09

v>3.3 s=5.6

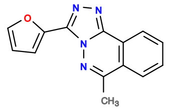

405D09

v>3.6 s=4.9

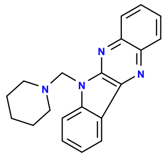

46D07

v>3.4 s=4.8

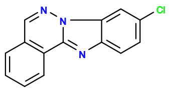

167G05

v>3.5 s=4.5

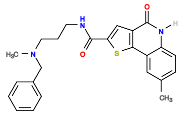

328F08

v>3.5 s=4.5

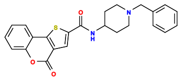

206C09

v>3.6 s=4.4

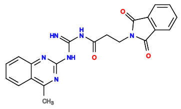

125F11

v>2.8 s=4.1

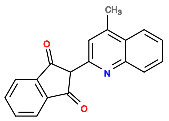

167H05

v>3.1 s=4.1

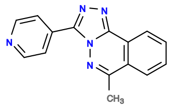

405G09

v>3.4 s=3.9

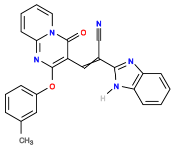

128A05

v>3.0 s=3.9

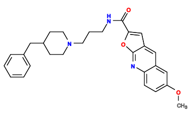

207H04

v>3.2 s=3.7

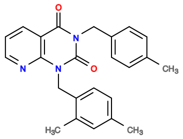

328B03

v>3.6

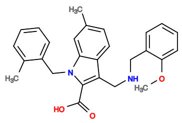

526C11

v>3.6

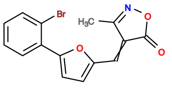

126B06

v>3.6

**514037045 GCTGTC ACTGGA AGCATGACCATATTTGGTGTGCCGAGCGGC TCCGGT CTGATGA GTCTT TGGC AGGAC GAAACAGC**

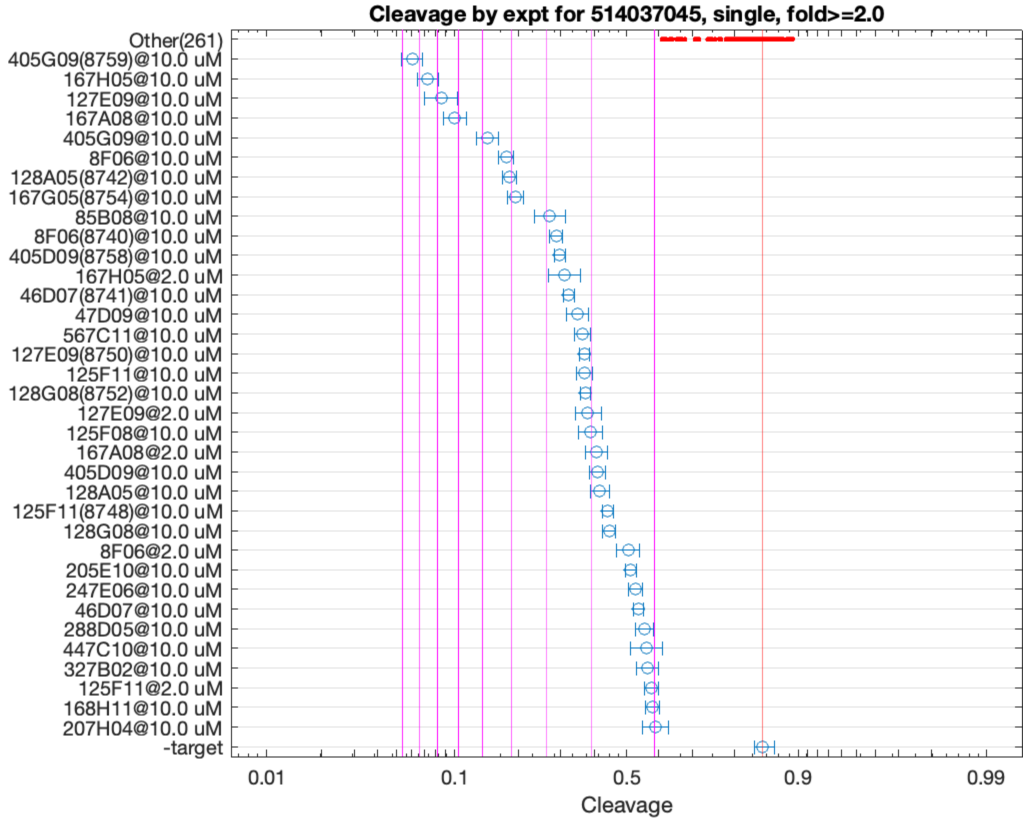


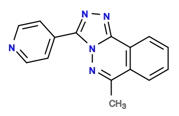

405G09

v>5.8 s=9.4

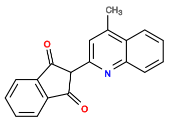

167H05

v>3.2 s=8.5

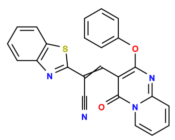

127E09

v>3.8 s=7.8

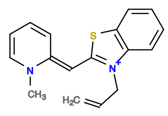

167A08

v>3.5 s=7.2

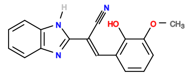

8F06

v>2.9 s=5.2

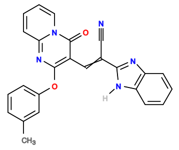

128A05

v>3.4 s=5.1

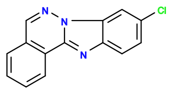

167G05

v>3.6 s=4.8

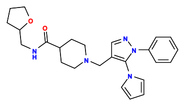

446C03

v>3.9

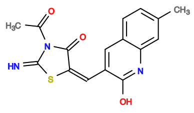

85B08

v>3.7 s=3.9

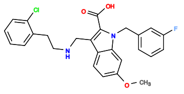

526H09

v>3.8

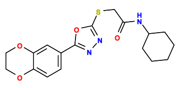

246A07

v>3.8

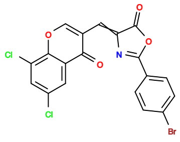

564H11

v>3.8

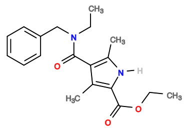

325F11

v>3.7

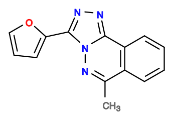

405D09

v>3.2 s=3.7

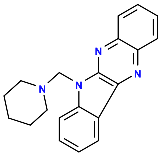

46D07

v>3.5 s=3.5

**514294722** GCTGTC ACTGGA AAGTTGCGTGAGTAAGGTGCGGTAAGATCA TCCGGT CTGATGA GTCC GTTGACCCGTGCTTGAGTAGCGTCGACTGGG GGAC GAAACAGC

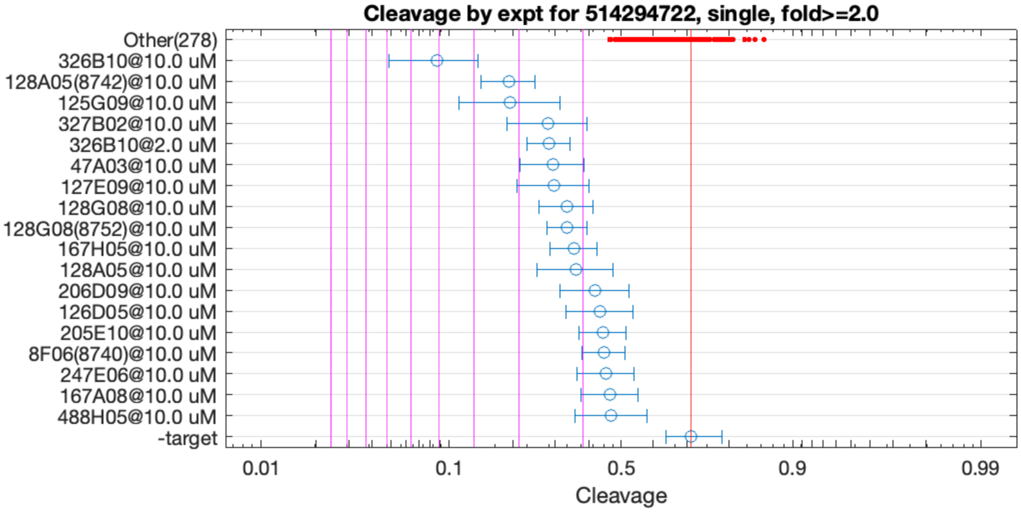


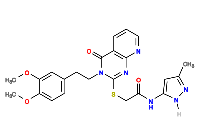

326B10

v>2.2 s=5.2

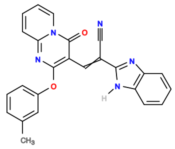

128A05

v>1.6 s=3.2

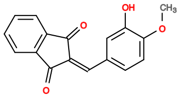

125G09

v>1.5 s=3.2

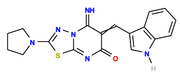

327B02

v>1.8 s=2.5

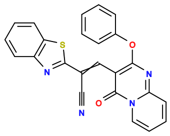

127E09

v>1.3 s=2.5

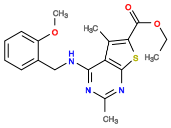

47A03

v>1.4 s=2.4

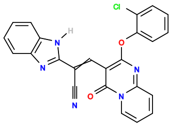

128G08

v>1.9 s=2.2

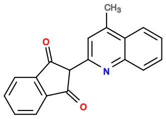

167H05

v>1.6 s=2.1

**514868644 GCTGTC ACTGGA TGCATGATCATATTTGGTGTGCTGAGCTGC TCTAGT CTGACGA GTCC GTGGCG GGAC GAAACAGC**

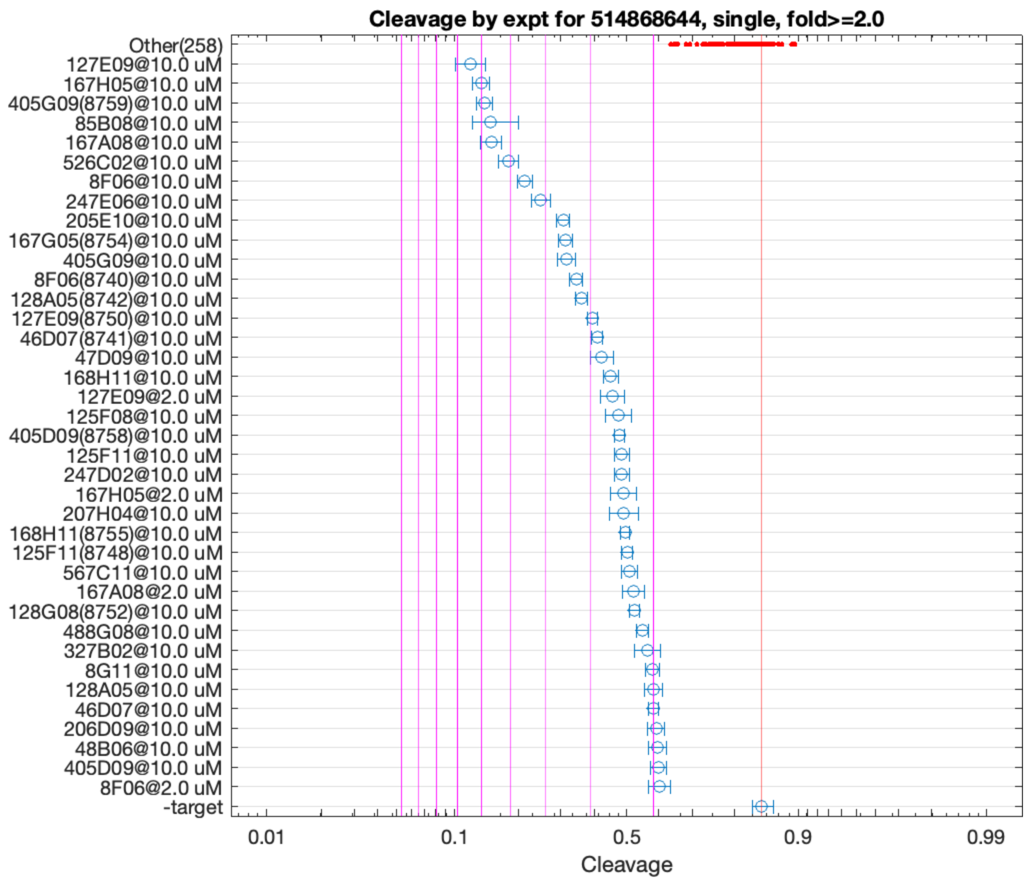


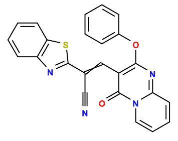

127E09

v>3.4 s=6.5

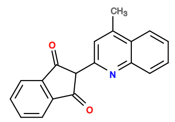

167H05

v>3.5 s=6.0

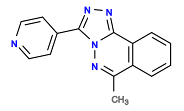

405G09

v>4.3 s=5.9

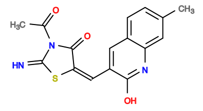

85B08

v>4.6 s=5.7

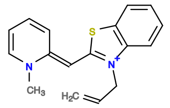

167A08

v>3.4 s=5.6

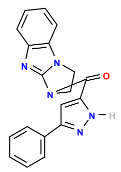

526C02

v>4.8 s=5.0

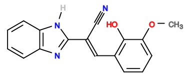

8F06

v>2.8 s=4.6

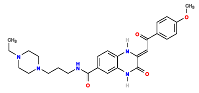

247E06

v>3.3 s=4.1

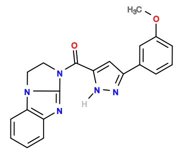

525D11

v>4.0

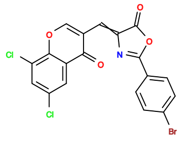

564H11

v>4.0

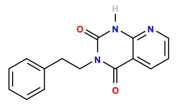

328H04

v>3.9

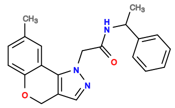

207F08

v>3.9

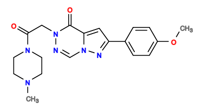

447E09

v>3.9

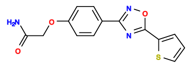

125B06

v>3.8

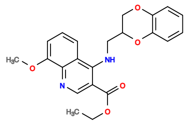

205E10

v>3.5 s=3.6

**524192145 GCTGTC ACTGGA TTTCCAACTTGCGTAATTTGATGACGCCTC TCCGGT CTGATGA GTCC GATGTGTCTGCGTGTTGGTG GGAC GAAACAGC**

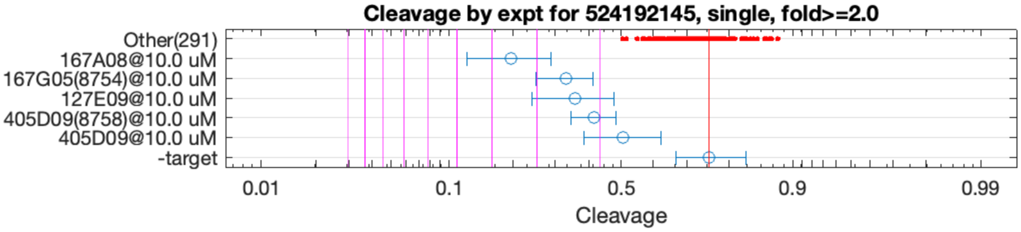


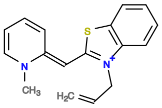

167A08

v>1.4 s=3.6

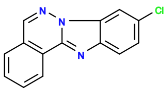

167G05

v>1.3 s=2.5

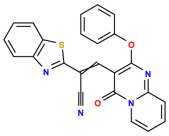

127E09

v>1.3 s=2.3

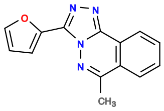

405D09

v>1.3 s=2.1

**533994655 GCTGTC ACTGGA TGGAGGCT TTCAGT CTGACGA GTCCTG CAACAACCAATTGCACGCAAGCC CGGGAC GAAACAGC**

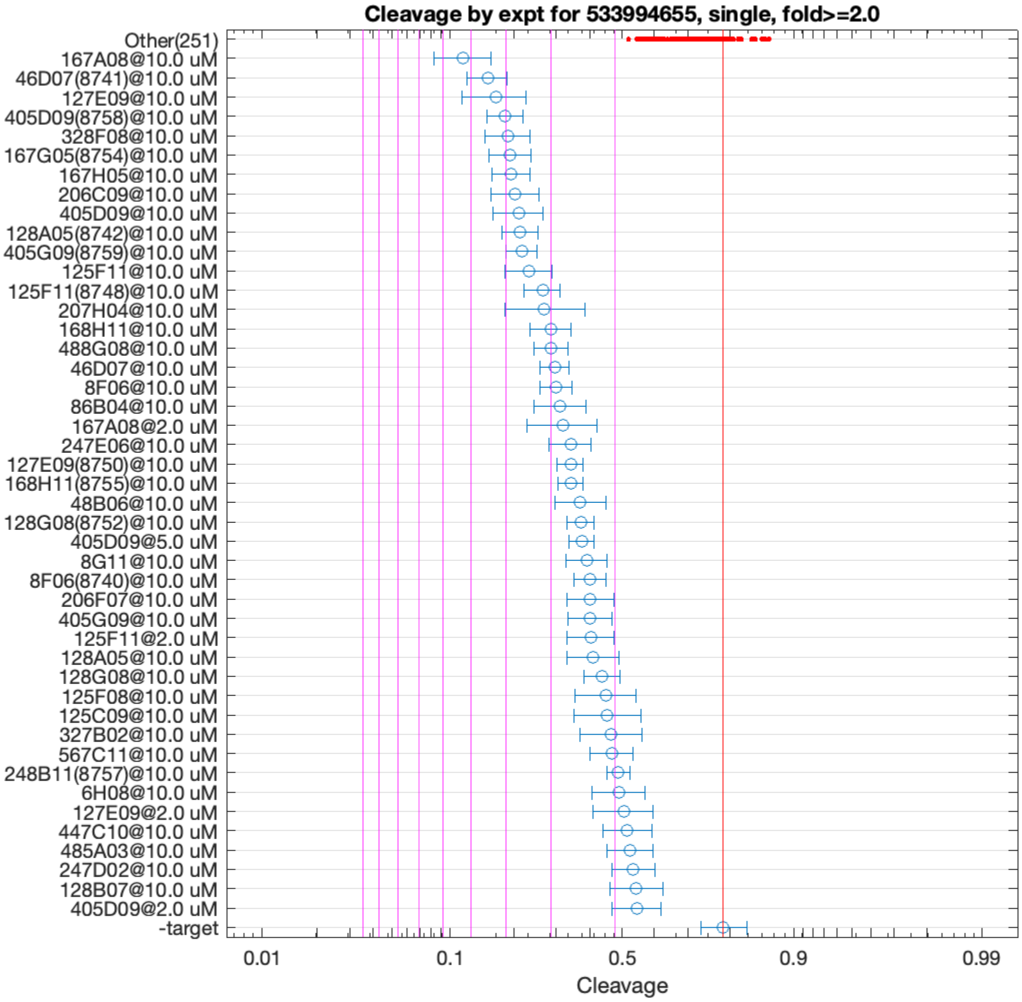


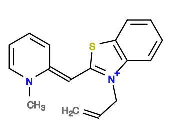

167A08

v>3.4 s=5.3

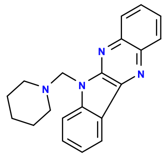

46D07

v>3.2 s=4.5

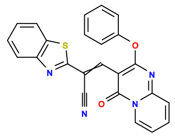

127E09

v>3.0 s=4.3

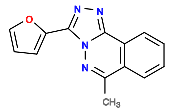

405D09

v>3.3 s=4.1

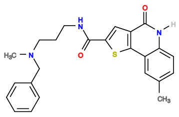

328F08

v>3.2 s=3.9

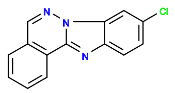

167G05

v>3.1 s=3.9

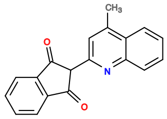

167H05

v>2.8 s=3.8

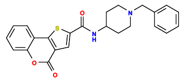

206C09

v>3.0 s=3.8

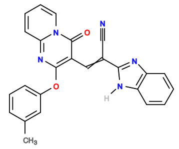

128A05

v>2.6 s=3.6

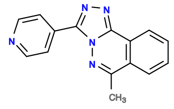

405G09

v>3.0 s=3.6

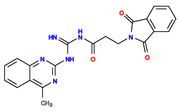

125F11

v>2.6 s=3.5

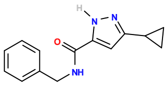

525B07

v>3.3

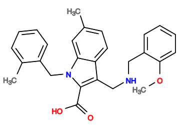

526C11

v>3.3

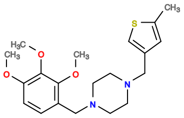

486G03

v>3.3

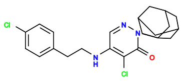

247D09

v>3.2

**535993570 GCTGTC AC TGGATTGGCGTAAATCC GT CTGATGA GTCC TGCGATGCCTAATGCACGTGCCTCTCCGCC GGAC GAAACAGC**

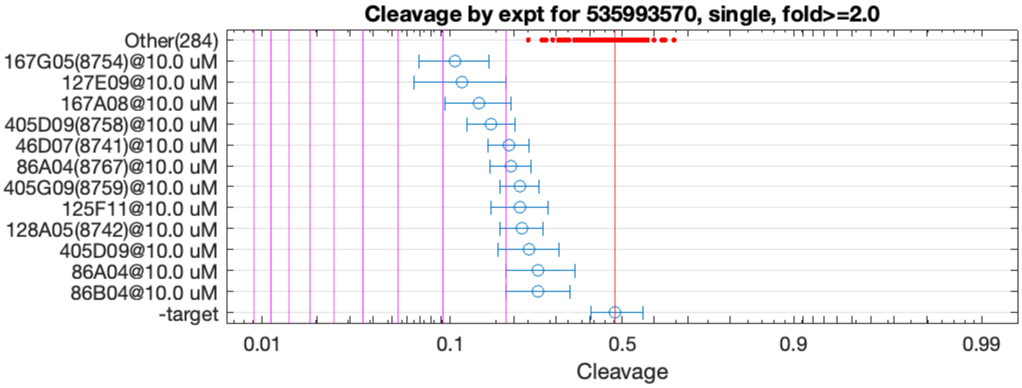


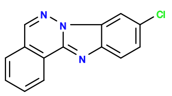

167G05

v>1.7 s=2.8

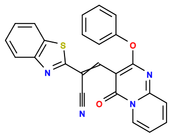

127E09

v>1.6 s=2.6

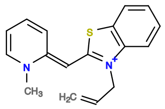

167A08

v>1.7 s=2.4

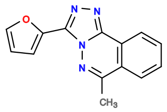

405D09

v>1.6 s=2.2

**536070526 GCTGTC ACTGGA GTAACCGATCTTTCGGCGAACTTACCGATA TCCGGT CTGACGA GTCT GTGGTG GGAC GAAACAGC**

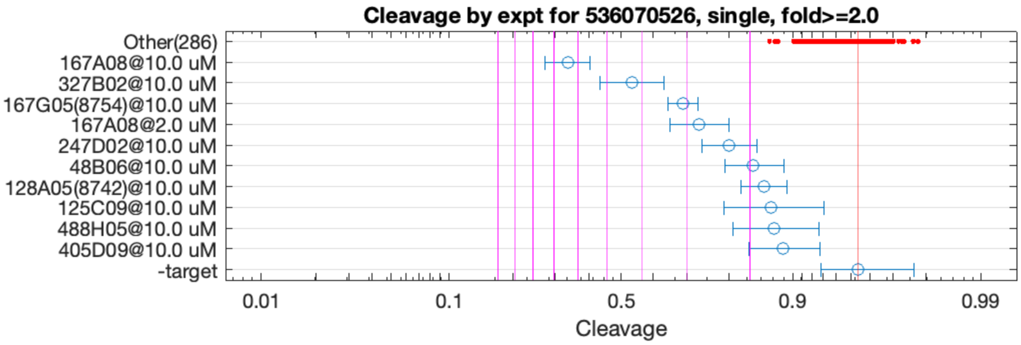


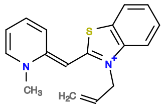

167A08

v>1.5 s=6.4

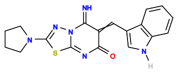

327B02

v>1.3 s=4.2

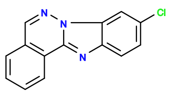

167G05

v>1.1 s=3.1

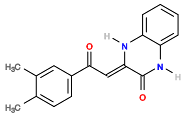

247D02

v>1.1 s=2.3

**536384056 GCTGTC AC TGGAATGCGCGAACT GT CTGATGA GTCC AAATCTGCTCGATTTCGTGTGGGTGCGCG GGAC GAAACAGC**

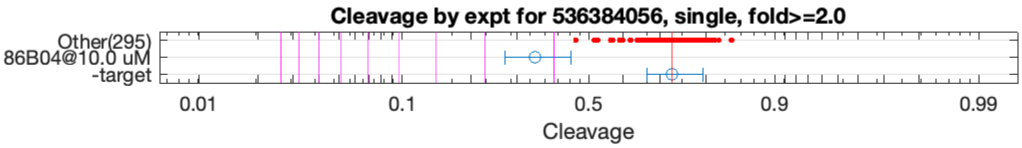


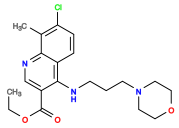

86B04

v>1.1 s=2.3

**565352371 GCTGTC ACTG TATTTTCCATCACTGGATTTGAT TGGT CTGATGA GTGCGTGTATGCC GAAACAGC TCGTGATGCCCCGGGACGAAACAGC**

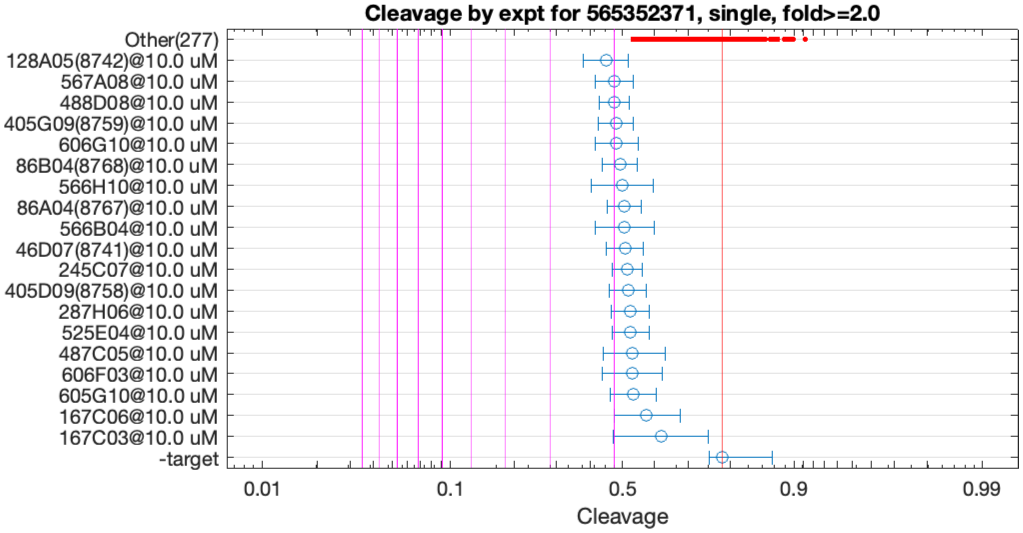


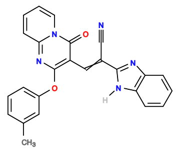

128A05

v>0.8 s=2.1

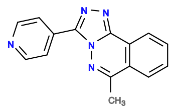

405G09

v>0.8 s=2.0

**565352652 GCTGTC ACTGGA AAGGTGGTAGGAAGAGTGCTCGTCGTAACA TCCGGT CTGATGA GTCC GATAAAACGTCATGTAGAAGCAGTACCACG GGAC GAAACAGC**

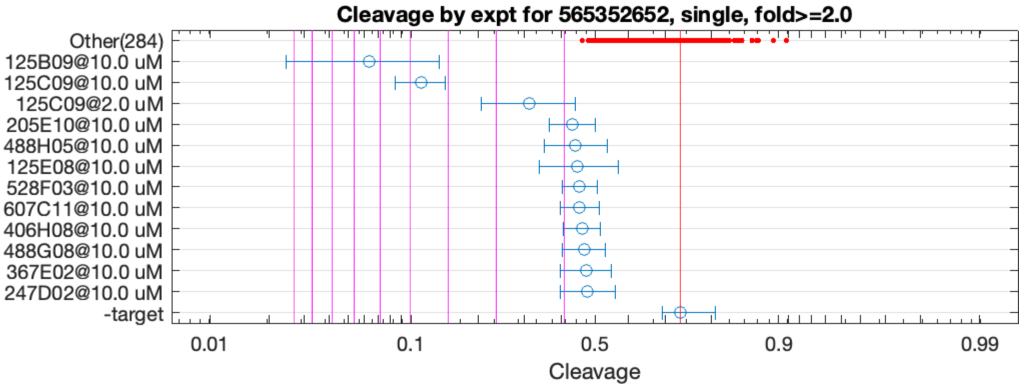


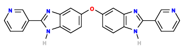

125B09

v>4.0 s=6.4

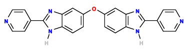

125C09

v>4.4 s=4.7

**565352773 GCTGTC ACTGGAA CGGGACA TTCTGGT CTGATGA GTCT ACTGCCTCTGCACTATCTGCAAGCGTACCG GGAC GAAACAGC**

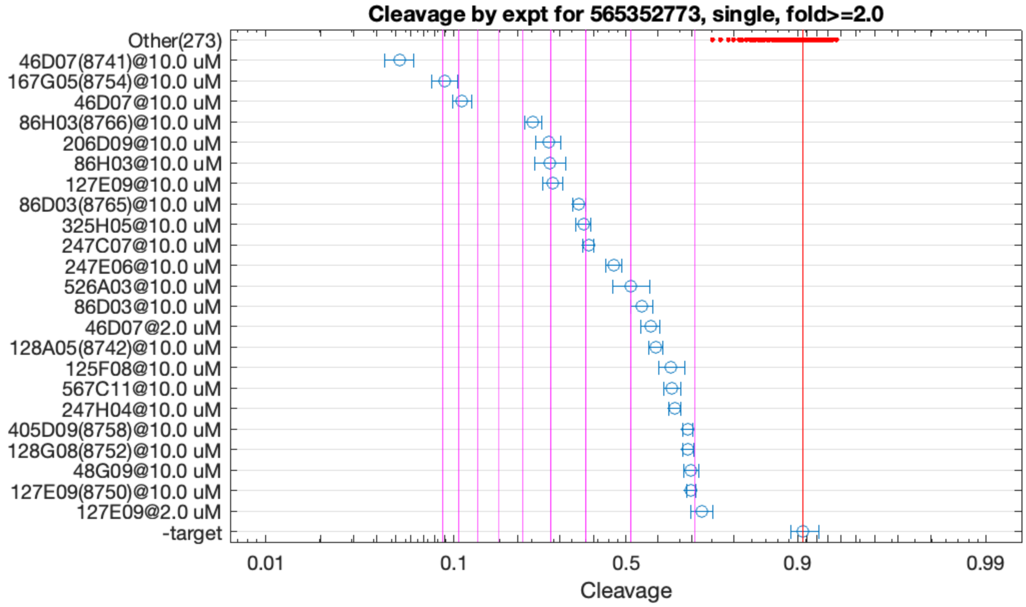


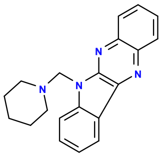

46D07

v>7.7 s=13.2

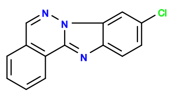

167G05

v>4.6 s=9.8

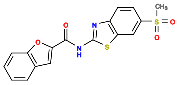

86H03

v>3.4 s=5.6

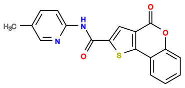

206D09

v>4.6 s=5.1

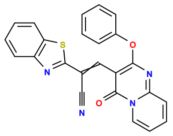

127E09

v>2.3 s=5.0

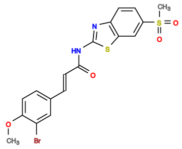

86D03

v>3.9 s=4.2

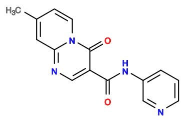

325H05

v>4.0 s=4.1

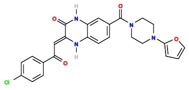

247C07

v>3.2 s=3.9

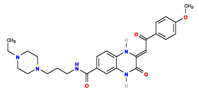

247E06

v>4.2 s=3.4

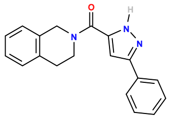

526A03

v>2.2 s=3.0

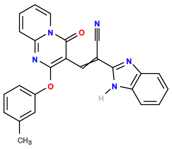

128A05

v>2.0 s=2.6

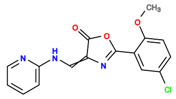

567C11

v>2.5 s=2.3

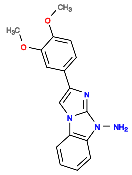

125F08

v>2.5 s=2.3

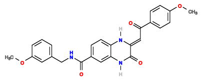

247H04

v>2.3 s=2.3

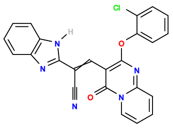

128G08

v>2.2 s=2.1

**565352780 GCTGTC ACTGGAA CTAGAT TTTCGGT CTGAAGA GTCC ACGCATCTCTGTGTGTGTTGGTGGGACGAAACAGGTTGTGTTGCC GGAC GAAACAGC**

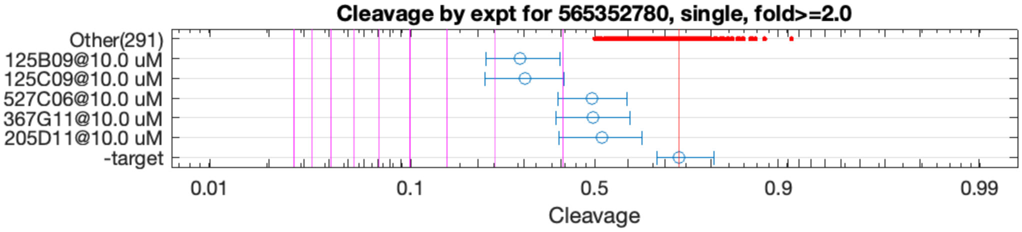


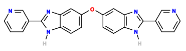

125B09

v>2.0 s=2.6

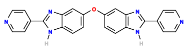

125C09

v>2.1 s=2.6

**565352815 GCTGTC ACTGGA AGCGTCTACGACAGGTAGGGAGTGACACAA TCCGGT CTGATGA GCACAAC GAAACAGC ACTAACGCCGGACGAAACAGC**

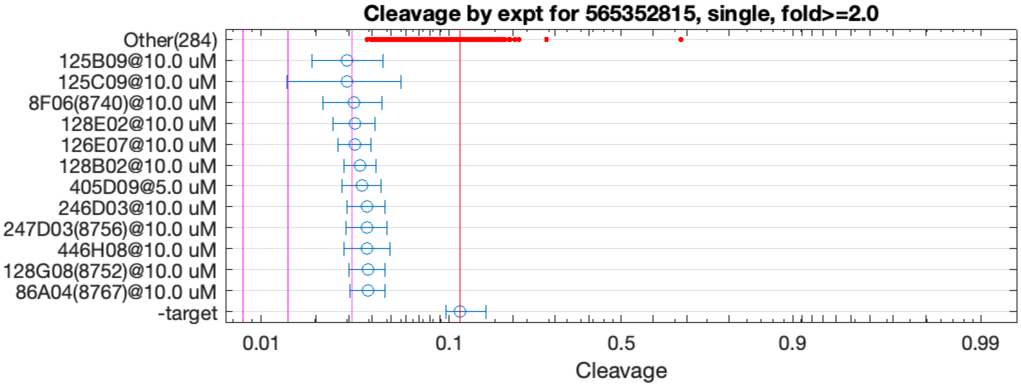


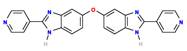

125C09

v>1.7 s=2.1

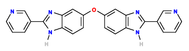

125B09

v>1.5 s=2.1

**565352879 GCTGTC ACTGGA ATGCGAAAAGTGGGGAATTGAAGAGCCTCA TCCGGT CTGATGA GTG ATGCCA CAC GAAACAGC CTCTTGTGCCGGACGAAACAGC**

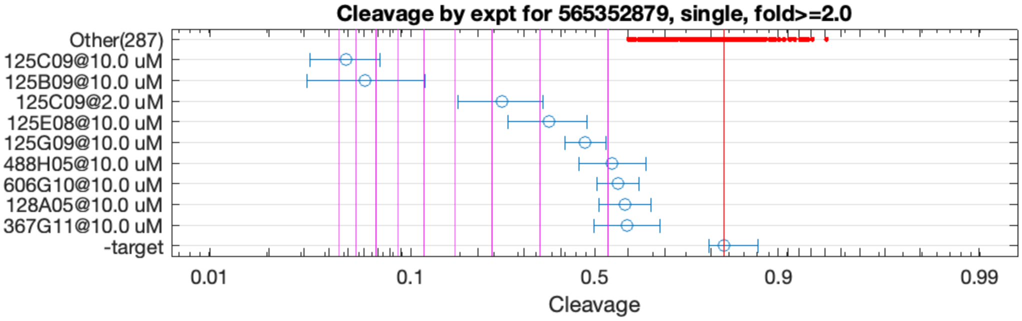


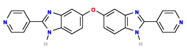

125C09

v>4.8 s=9.6

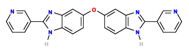

125B09

v>4.9 s=8.5

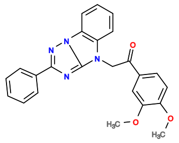

125E08

v>1.1 s=2.9

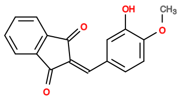

125G09

v>1.5 s=2.3

**565353287 GCTGTC ACTGGA GACCTACGCGGTCAGCGCAAACCTCGCATA TCCGGT CTGACGA GTCTC ATAGACGAAACAGCGACGCCCC GGGAC GAAACAGC**

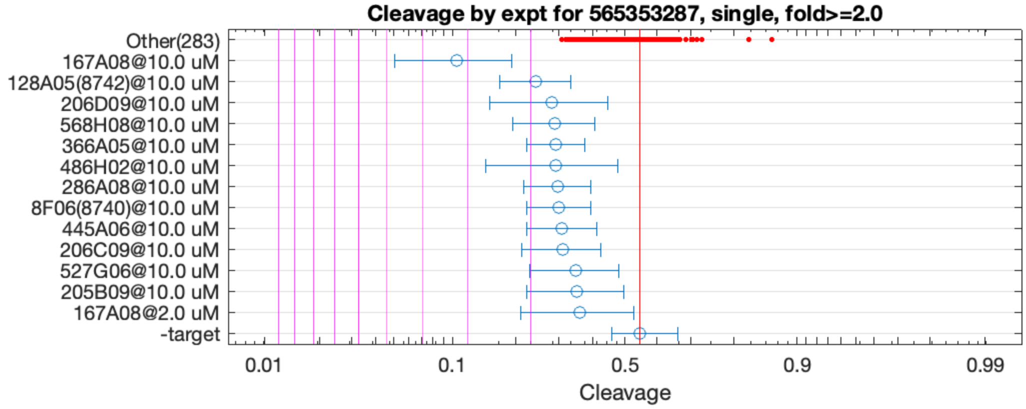


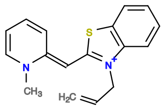

167A08

v>1.2 s=3.2

**565353501 GCTGTC ACTGGA TAAGAAGAGCTACTAGGTAGGGAAGAGGCT TCCGGT CTGATGA GACCTTGAATGGC GAAACAGC ACTACGTTGCCCGGGACGAAACAGC**

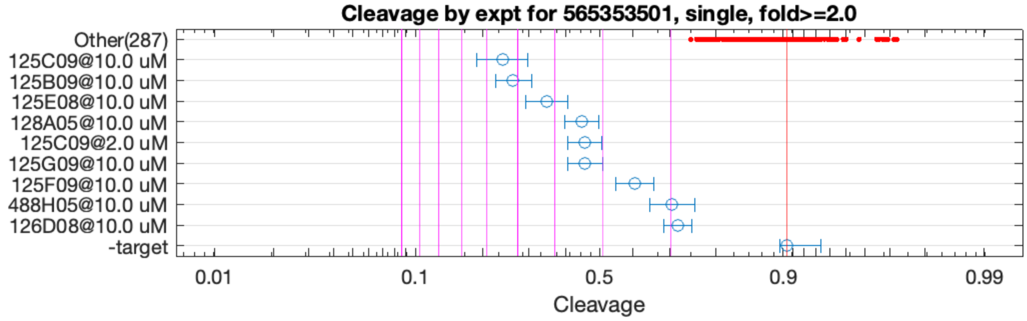


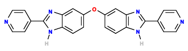

125C09

v>4.3 s=5.4

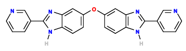

125B09

v>4.1 s=5.1

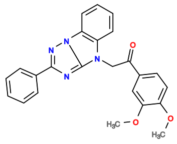

125E08

v>1.8 s=4.2

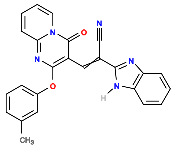

128A05

v>1.2 s=3.4

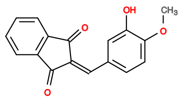

125G09

v>2.5 s=3.3

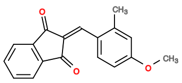

125F09

v>1.9 s=2.5

**565354562 GCTGTC ACTG GATTTTCAGTTCGACGAACACTGGATACCGTAAA CAGT CTGAAGA GTCC AAATCTGCTCGATCGCGTGTGGGTGCGGG GGAC GAAACAGC**

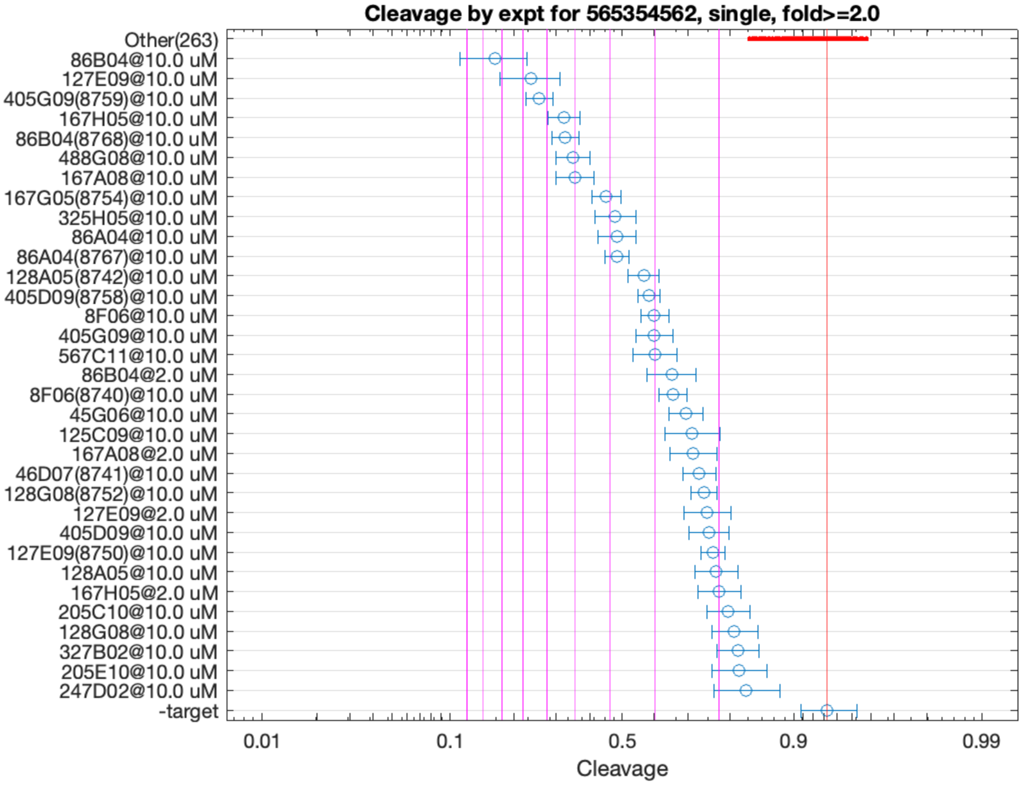


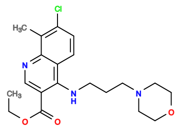

86B04

v>3.5 s=8.4

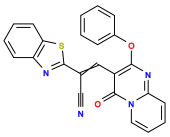

127E09

v>2.6 s=6.5

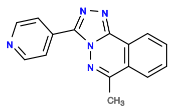

405G09

v>3.4 s=6.3

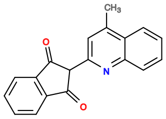

167H05

v>2.6 s=5.4

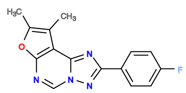

488G08

v>2.7 s=5.1

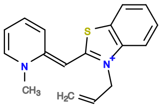

167A08

v>2.3 s=5.0

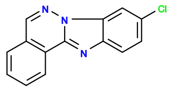

167G05

v>2.8 s=4.1

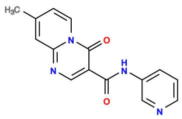

325H05

v>3.6 s=3.9

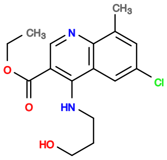

86A04

v>2.8 s=3.8

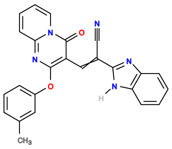

128A05

v>2.1 s=3.2

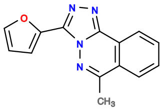

405D09

v>2.4 s=3.1

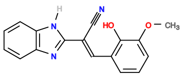

8F06

v>2.1 s=3.0

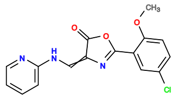

567C11

v>2.5 s=2.9

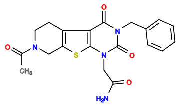

327A11

v>2.9

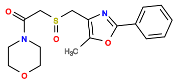

285E02

v>2.8

**565354742 GCTGTC ACTGG ATTTTCCAGGATCACTGGATGCATGACCATATCTTGGTGTGCTGAGCGGCA CCAGT CTGATGA GTTC GTGGCG GGAC GAAACAGC**

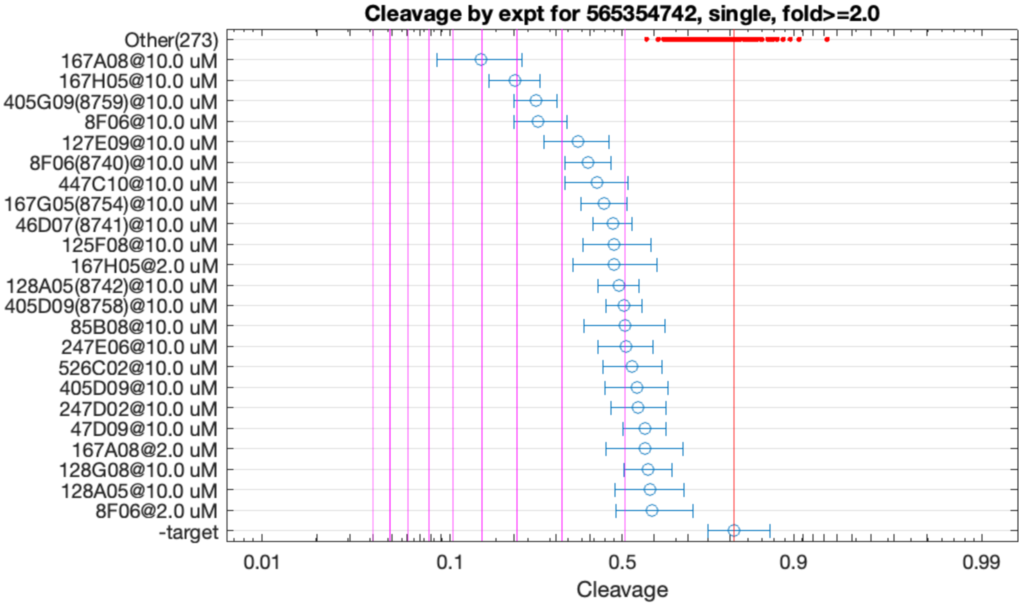


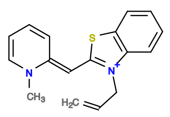

167A08

v>2.5 s=5.0

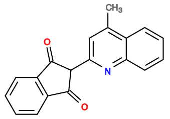

167H05

v>2.5 s=4.0

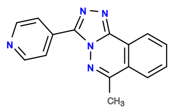

405G09

v>2.4 s=3.5

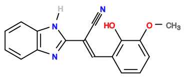

8F06

v>1.9 s=3.5

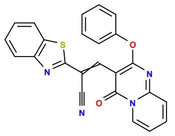

127E09

v>1.9 s=2.7

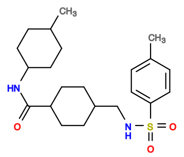

605E10

v>2.4

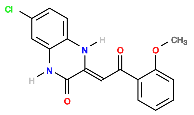

247C02

v>2.3

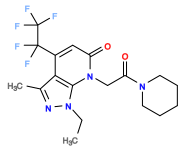

488F10

v>2.3

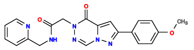

447C10

v>2.2 s=2.3

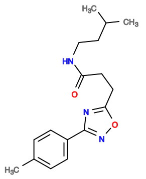

245A06

v>2.3

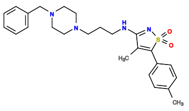

446E08

v>2.3

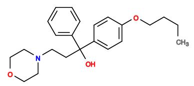

125E11

v>2.3

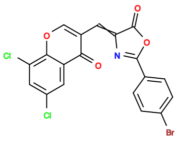

564H11

v>2.3

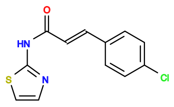

6D10

v>2.3

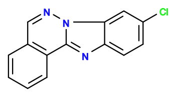

167G05

v>2.2 s=2.3

**565355200 GCTGTC ACTGG CCTTCCTTTTTCACTGGAACTGGATGCGAT CTAGT CTGACGA CTCCTGCGATGGAC GAAACAGC TTGGCCCGGGACGAAACAGC**

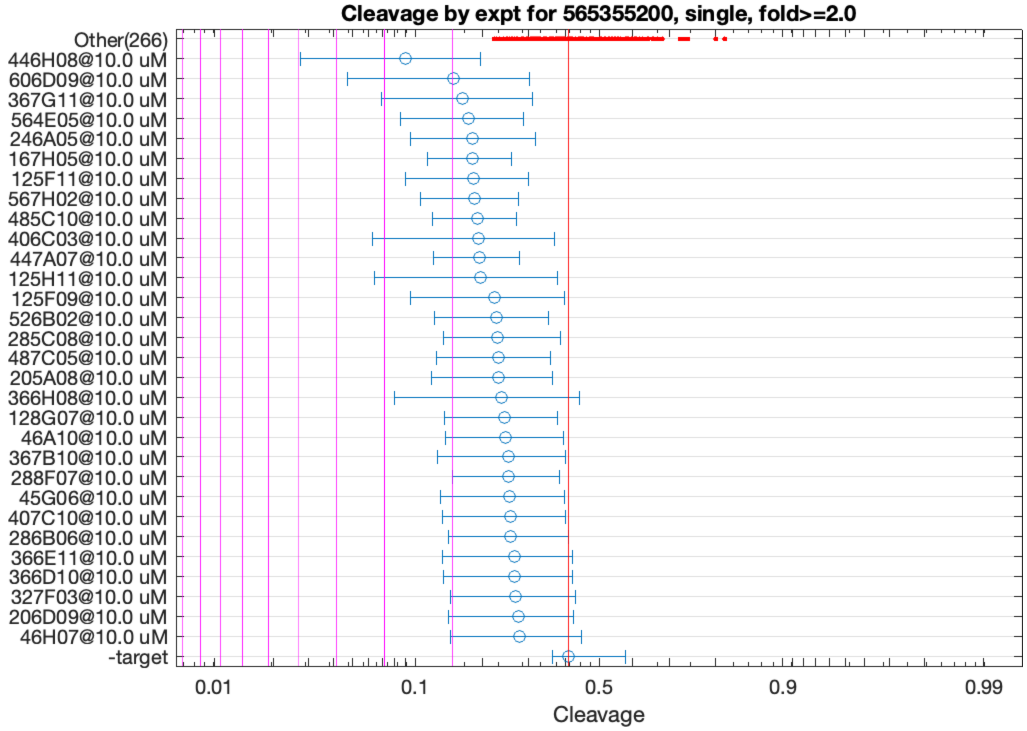


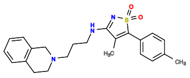

446H08

v>0.7 s=2.6

**565358693 GCTGTC ACTGGA AACTCTCTGAATGAAGCGGGTATCGCAATG TCCGGT CTGATGA GTCC GATGCGGTACAGGTCAGGGTTGAATGCG GGAC GAAACAGC**

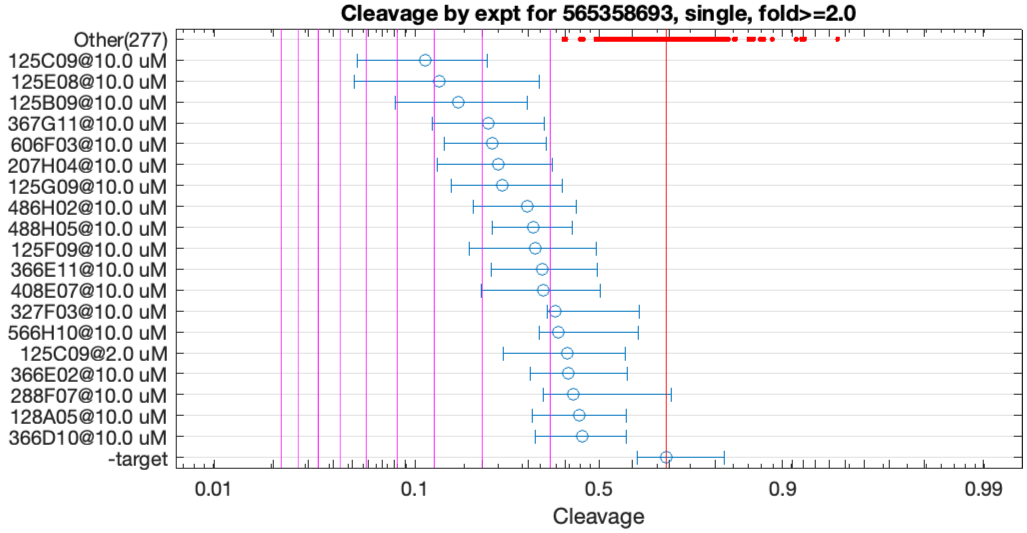


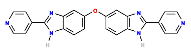

125C09

v>2.6 s=4.1

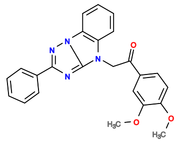

125E08

v>1.1 s=4.1

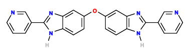

125B09

v>2.5 s=3.5

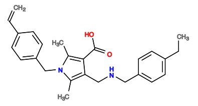

367G11

v>0.9 s=2.9

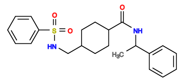

606F03

v>0.6 s=2.8

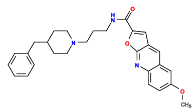

207H04

v>0.8 s=2.7

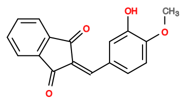

125G09

v>1.2 s=2.7

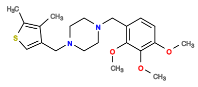

486H02

v>0.8 s=2.3

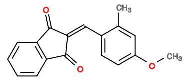

125F09

v>1.1 s=2.2

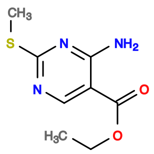

488H05

v>0.8 s=2.2

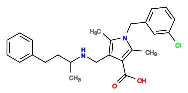

366E11

v>0.7 s=2.1

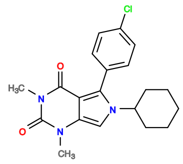

408E07

v>0.8 s=2.1

**565359918 GCTGTC ACTGGA ACTGGATGCATGACCATATTTGGTGTGCCGAGCTGCC TCTAGT CTGACGA GTCC GTGGCG GGAC GAAACAGC**

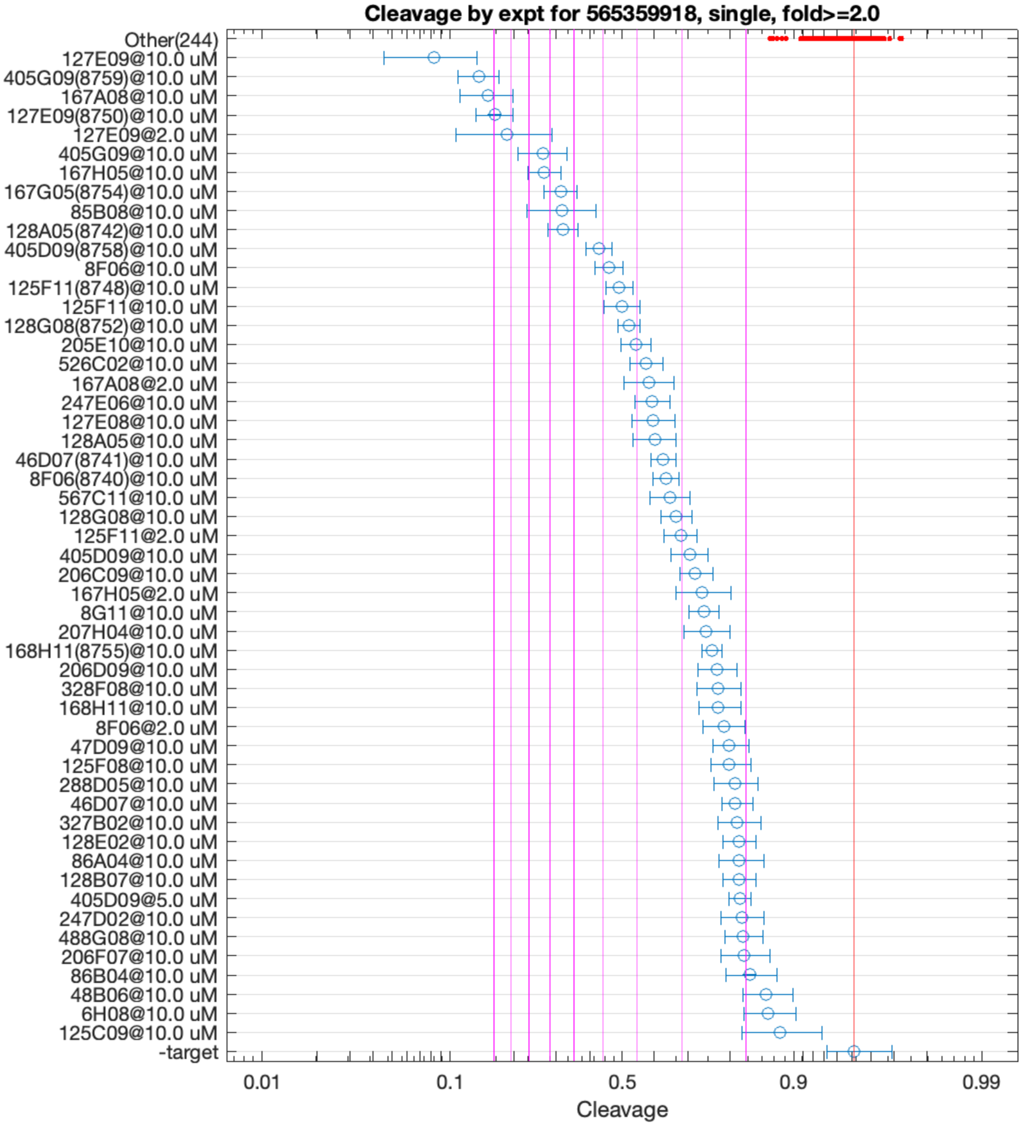


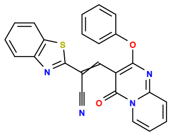

127E09

v>6.0 s=14.4

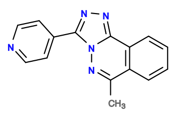

405G09

v>6.6 s=10.9

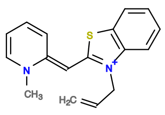

167A08

v>4.5 s=10.6

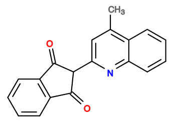

167H05

v>4.1 s=7.2

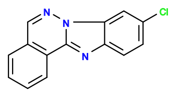

167G05

v>4.0 s=6.5

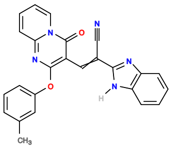

128A05

v>3.8 s=6.5

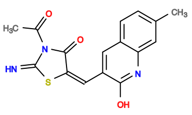

85B08

v>4.1 s=6.4

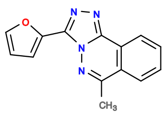

405D09

v>3.6 s=5.1

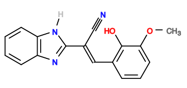

8F06

v>3.2 s=4.9

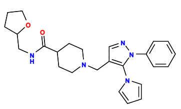

446C03

v>4.5

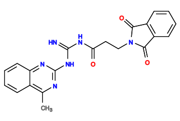

125F11

v>3.0 s=4.4

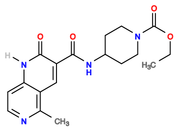

608B07

v>4.4

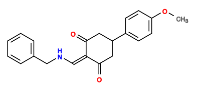

48D04

v>4.3

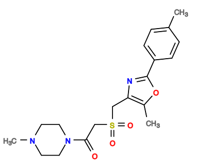

285A10

v>4.3

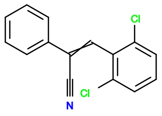

8G09

v>4.3

**565360115 GCTGTC ACTGGAA GAGCGCTTAACGTAAGTCACTGGATGA TTCCGGT CTGACGA GTCC AATGTCGGAGTGATTATCGTTGGCAGAGTG GGAC GAAACAGC**

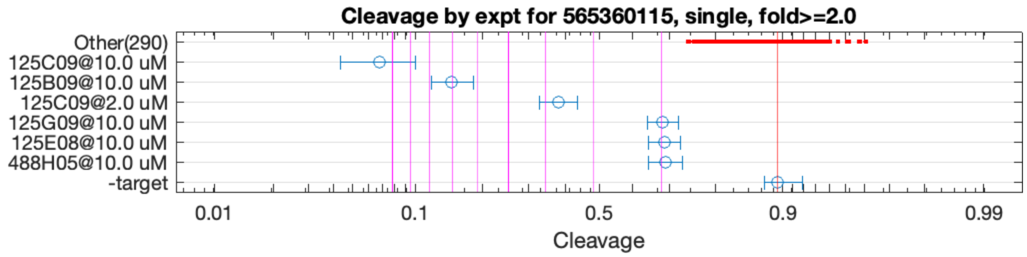


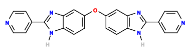

125C09

v>8.2 s=10.7

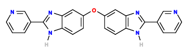

125B09

v>6.5 s=7.0

**565360436 GCTGTC ACTGGA AGGCATGTCTTTGAGTAGCCAAAACTGGAA TCCGGT CTGACGA GTCC GATACGTTGGTGAGTAGATGCATGTGCCTG GGAC GAAACAGC**

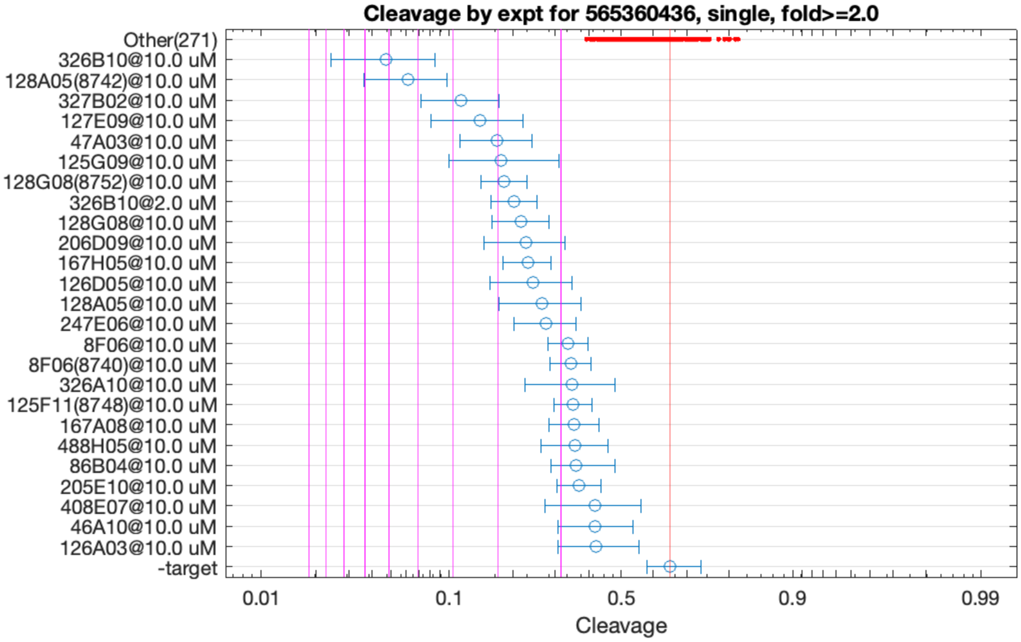


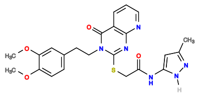

326B10

v>3.0 s=6.1

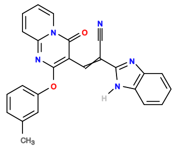

128A05

v>2.2 s=5.4

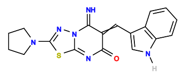

327B02

v>2.2 s=3.8

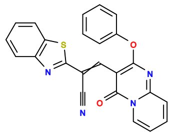

127E09

v>1.8 s=3.3

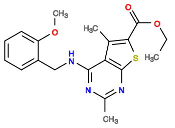

47A03

v>1.8 s=3.0

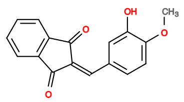

125G09

v>1.8 s=2.9

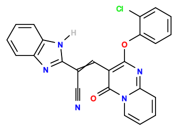

128G08

v>2.4 s=2.9

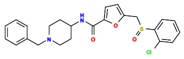

487E02

v>2.6

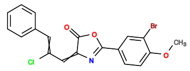

567G08

v>2.5

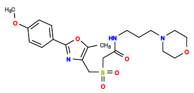

285B11

v>2.5

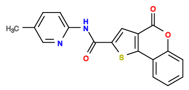

206D09

v>2.6 s=2.5

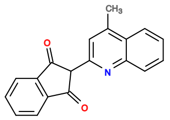

167H05

v>1.9 s=2.5

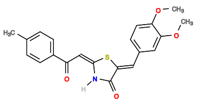

126D05

v>2.5 s=2.4

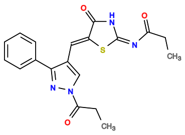

47A11

v>2.4

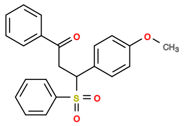

47A09

v>2.4

**565360460 GCTGTC ACTGGA AGGTACGTCTTTGAGTAGCCAAAACTGGAA TCCGGT CTGACGA GTCC GATACGTTGGTGAGTAGATGCATGTGCCTG GGAC GAAACAGC**

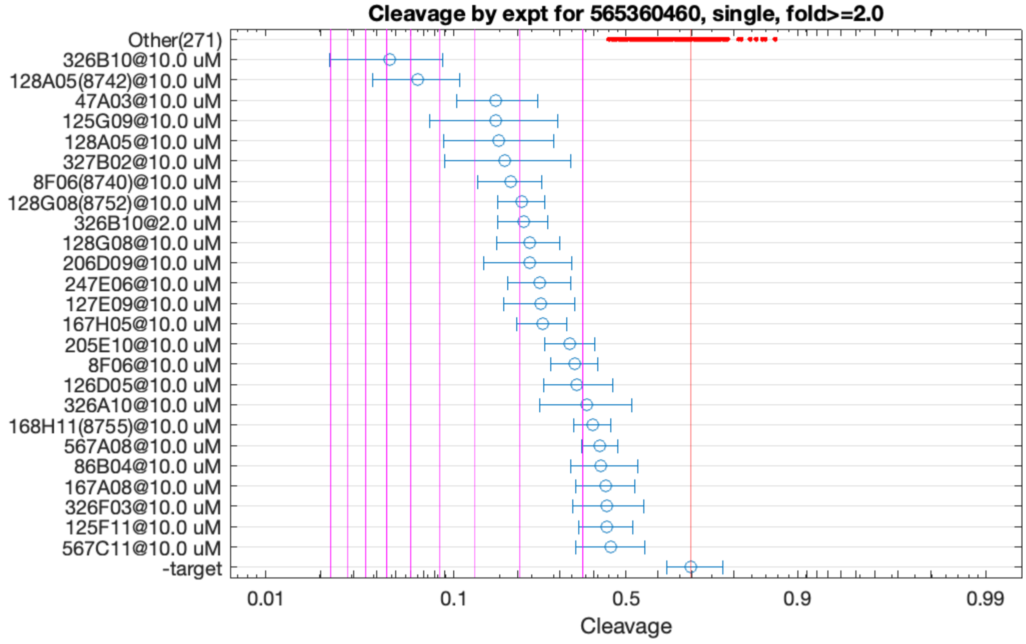


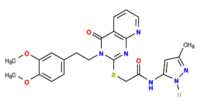
326B10
v>3.2 s=6.7

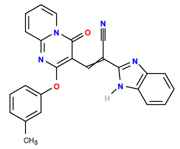
128A05
v>2.1 s=5.6

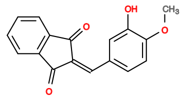
125G09
v>2.0 s=3.5

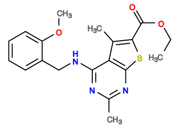
47A03
v>1.9 s=3.4

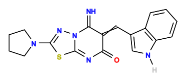
327B02
v>2.3 s=3.3

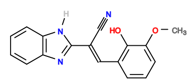
8F06
v>1.8 s=3.1

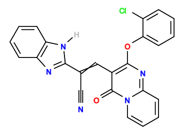
128G08
v>2.5 s=3.0

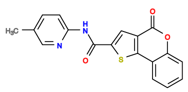
206D09
v>2.3 s=2.8

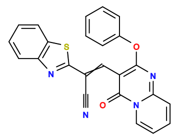
127E09
v>1.8 s=2.6

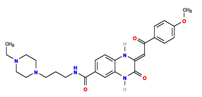
247E06
v>2.1 s=2.6

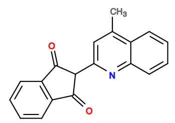
167H05
v>2.0 s=2.6

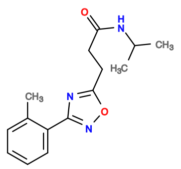
405C07
v>2.5

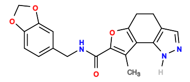
446E04
v>2.5

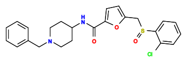
487E02
v>2.5

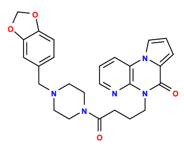
485A07
v>2.4

**565366116 GCTGTC ACTGGA GGAAGAATTCAAGTGTGAAAAGGTTGATCA TCCGGT CTGACGA GTCC GCAAACCTCAACGCATAACACTTGTGACTG GGAC GAAACAGC**

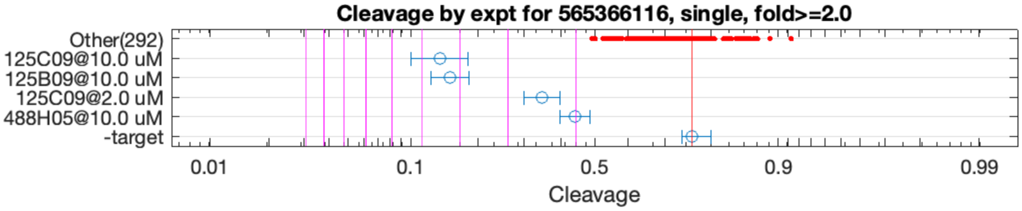


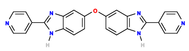

125C09

v>4.9 s=4.5

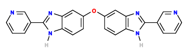

125B09

v>4.3 s=4.2

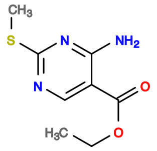

488H05

v>1.5 s=2.0

**565370758 GCTGTC AC TGGATTCTGGTTAGCTGCAACTGGAAATTCT GT CTGAAGA GTCC TTGGAAAGGTGGACGAAACAGCTGCGTGCC GGAC GAAACAGC**

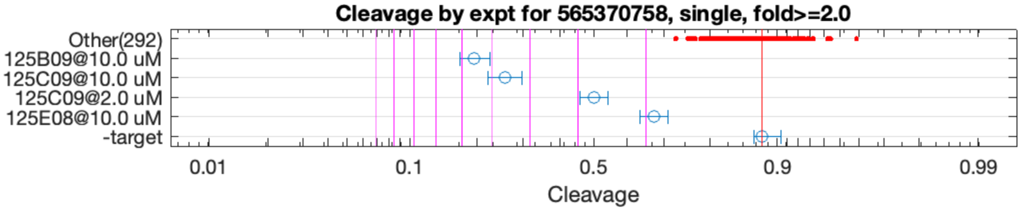


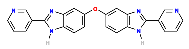

125B09

v>4.4 s=5.6

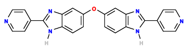

125C09

v>4.3 s=4.6

**565372067 GCTGTC ACTGG ATTTACTGGATTTTCGTG TCAGT CTGACGA GTCT TTGT GGAC GAAACAGC**

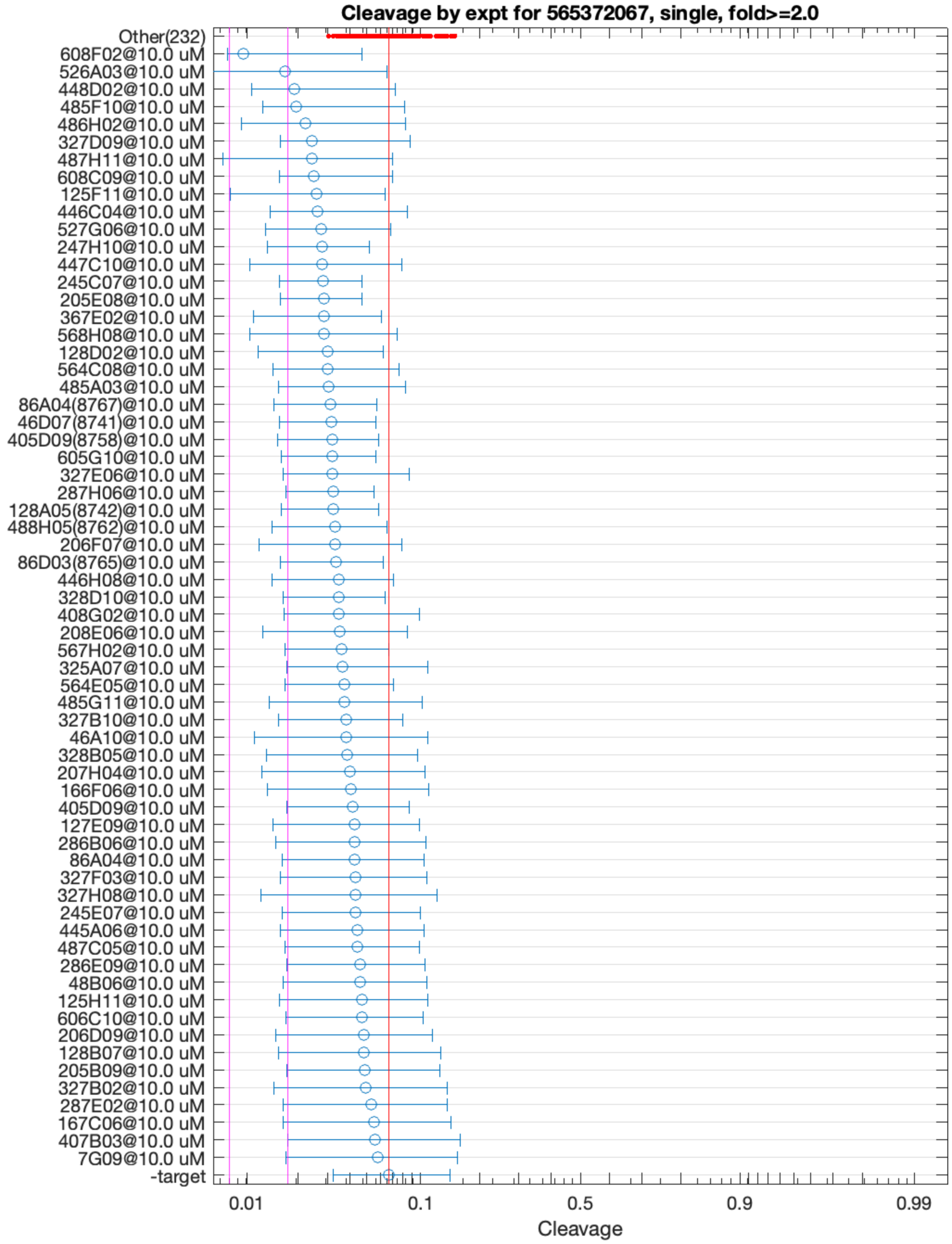


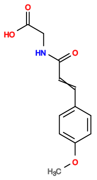

608F02

v>0.7 s=2.7

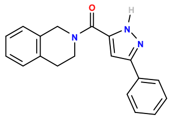

526A03

v>0.6 s=2.0

565378920 GCTGTC ACTG GATTTTCAGTTCGACGAACACTGGATACCGTAAA CAGT CTGAAGA GTCC AAATCTGCCGATTTTCGCGTGGGTGCGGG GGAC GAAACAGC

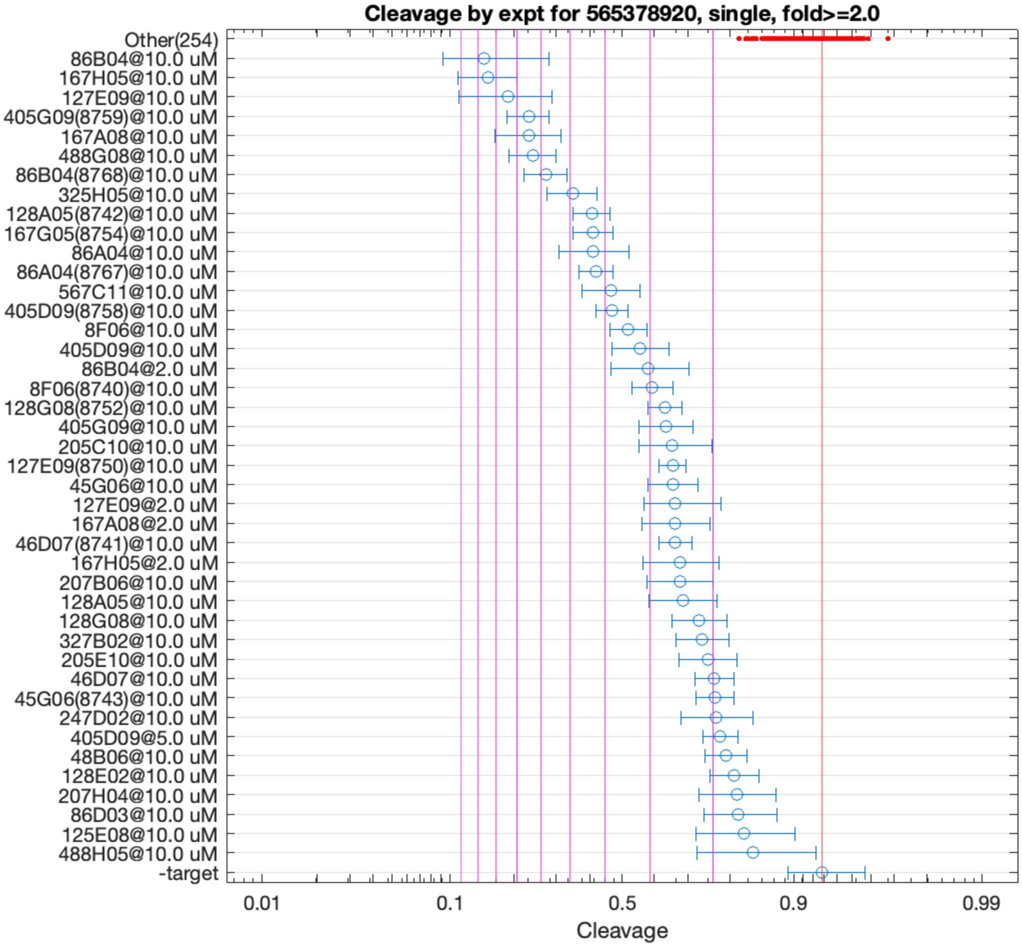


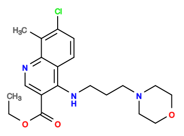

86B04

v>4.0 s=8.6

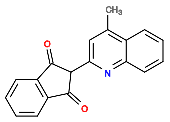

167H05

v>3.2 s=8.3

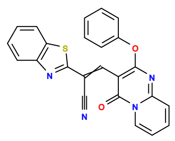

127E09

v>3.0 s=7.3

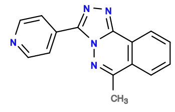

405G09

v>3.7 s=6.7

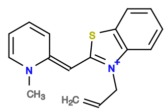

167A08

v>2.4 s=6.5

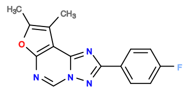

488G08

v>3.2 s=6.2

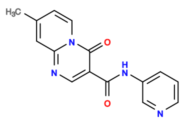

325H05

v>4.3 s=5.0

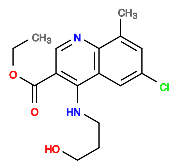

86A04

v>2.9 s=4.4

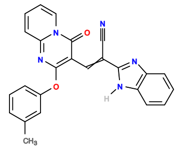

128A05

v>2.5 s=4.4

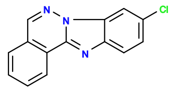

167G05

v>2.9 s=4.3

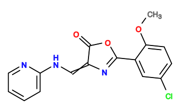

567C11

v>2.7 s=3.8

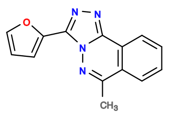

405D09

v>2.6 s=3.8

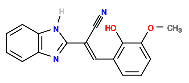

8F06

v>2.3 s=3.4

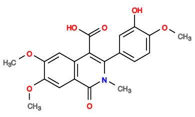

206H06

v>3.2

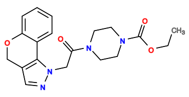

328C11

v>3.1

**565380623 GCTGTC ACTGG ATTTTCCAGGATCACTGGATGCATGACCATATTTGGTGTGCCGAGCGGCA CCAGT CTGACGA GTCT GTGGCG GGAC GAAACAGC**

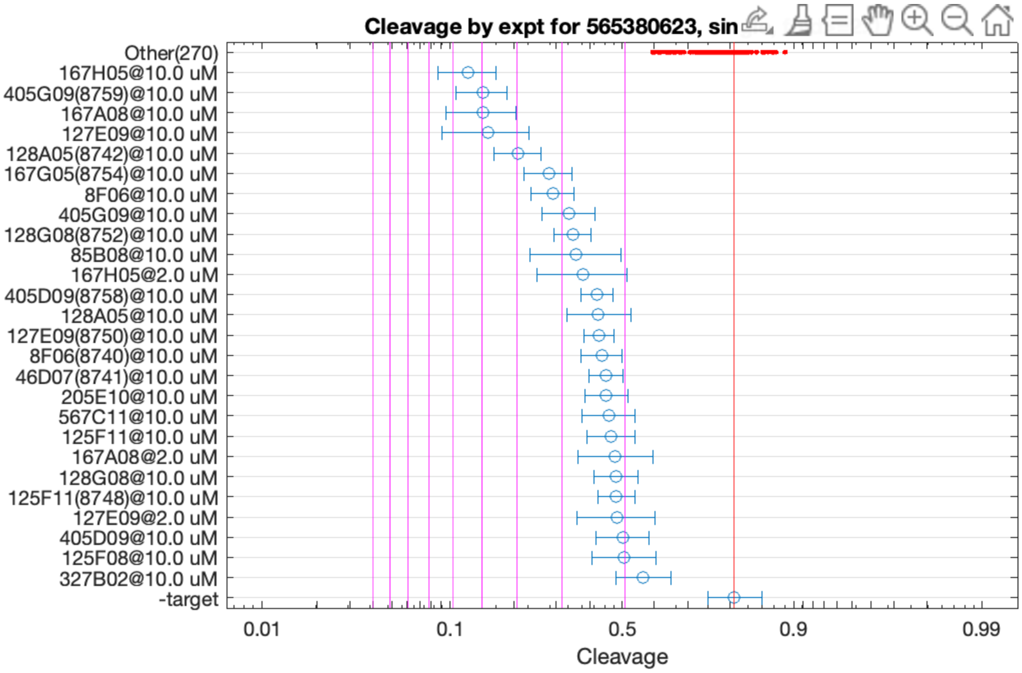


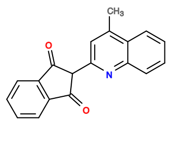

167H05

v>2.4 s=5.5

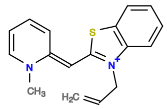

167A08

v>2.5 s=5.0

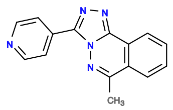

405G09

v>3.2 s=4.9

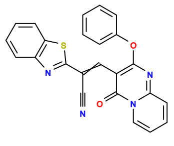

127E09

v>2.6 s=4.8

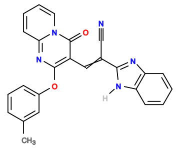

128A05

v>2.4 s=4.0

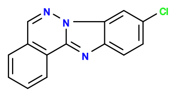

167G05

v>2.5 s=3.3

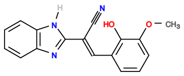

8F06

v>2.0 s=3.2

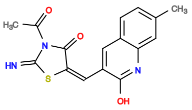

85B08

v>2.4 s=2.8

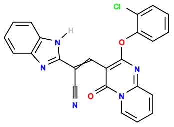

128G08

v>2.7 s=2.8

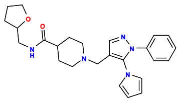

446C03

v>2.6

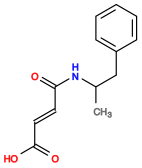

6E06

v>2.6

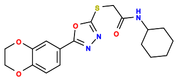

246A07

v>2.5

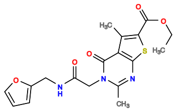

206G08

v>2.5

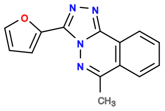

405D09

v>2.2 s=2.4

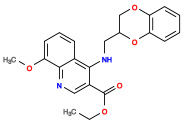

205E10

v>2.1 s=2.2

**565385129 GCTGTC ACTG GATTTTGGTGGATGACGAAACACTGGATACCGTAAA CAGT CTGAAGA GTCC AAATCTGCTCGATGTCGTGTGGGTGCGGG GGAC GAAACAGC**

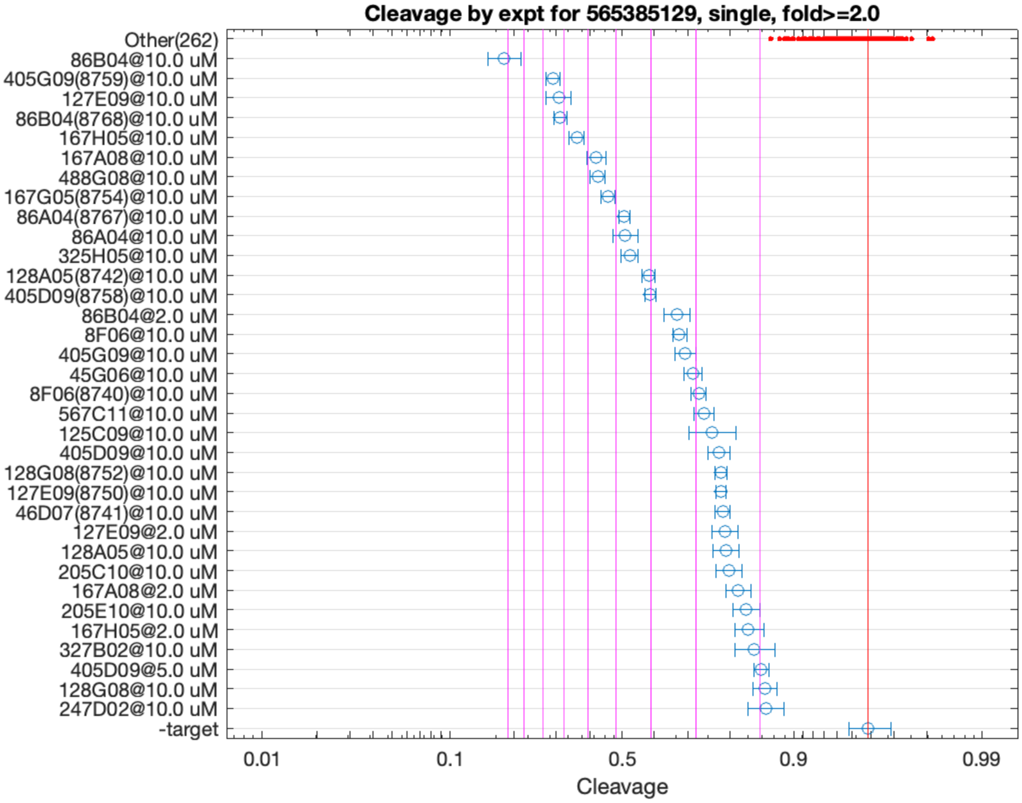


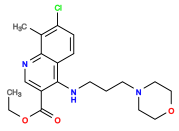

86B04

v>4.4 s=10.2

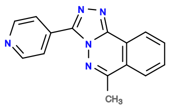

405G09

v>3.7 s=7.4

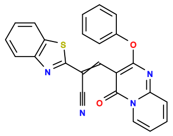

127E09

v>2.9 s=7.2

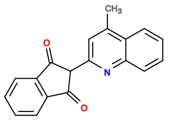

167H05

v>3.1 s=6.5

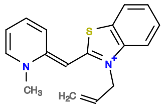

167A08

v>2.4 s=5.7

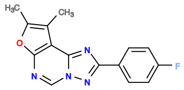

488G08

v>2.9 s=5.6

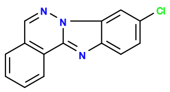

167G05

v>3.2 s=5.3

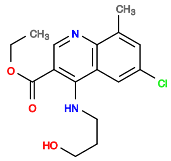

86A04

v>3.3 s=4.8

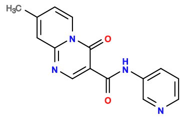

325H05

v>4.4 s=4.6

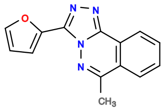

405D09

v>2.7 s=4.0

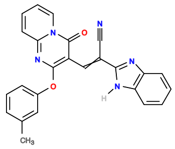

128A05

v>2.4 s=4.0

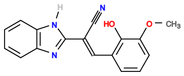

8F06

v>2.2 s=3.3

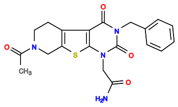

327A11

v>3.2

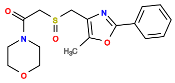

285E02

v>3.2

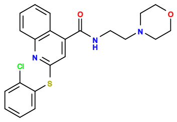

208H11

v>3.1

**565388300 GCTGTC ACTGGA ATGTGGCGTGAGTAAGGTGCGGTAAGATCA TCTGGT CTGATGA GTCC GTTGAACCGTGCTTGAGTAGCGTCGACTGGG GGAC GAAACAGC**

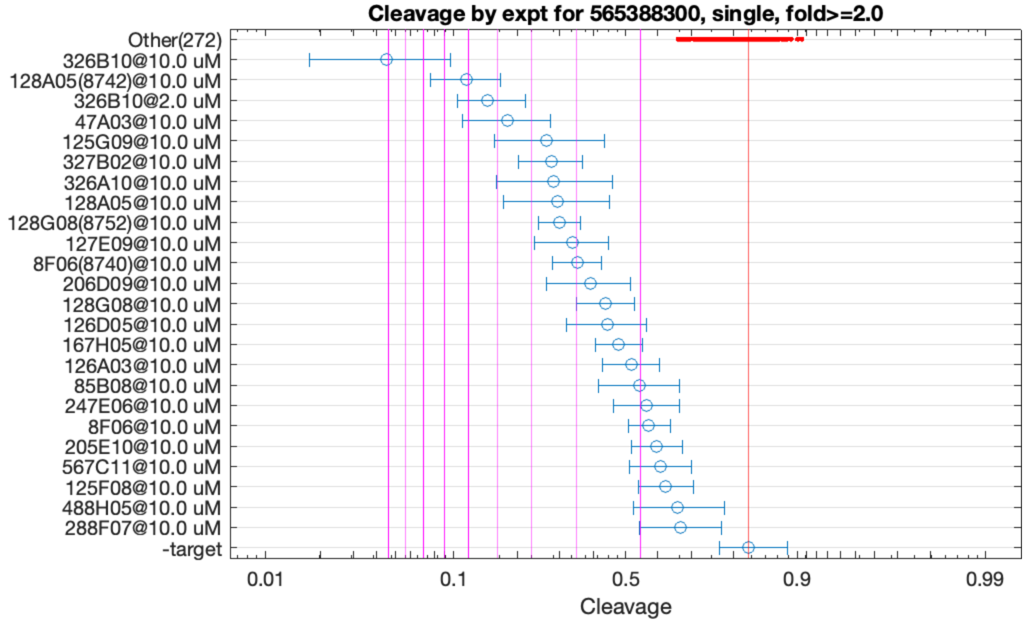


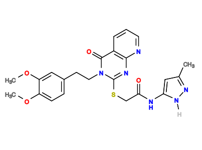

326B10

v>3.9 s=10.4

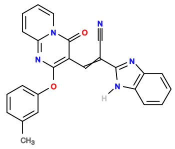

128A05

v>2.4 s=6.2

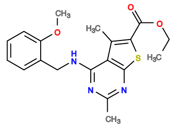

47A03

v>2.1 s=4.7

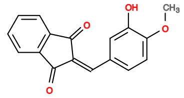

125G09

v>2.1 s=3.7

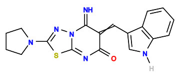

327B02

v>2.3 s=3.5

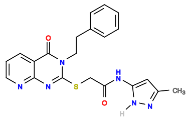

326A10

v>3.0 s=3.4

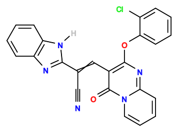

128G08

v>2.5 s=3.3

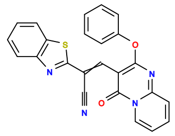

127E09

v>1.6 s=3.1

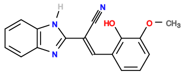

8F06

v>1.8 s=3.0

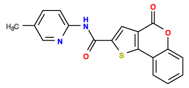

206D09

v>3.3 s=2.7

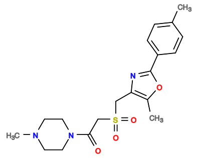

285A10

v>2.7

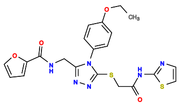

205E07

v>2.6

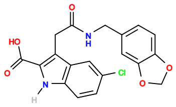

245E04

v>2.6

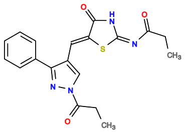

47A11

v>2.6

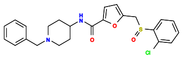

487E02

v>2.6

565411466 GCTGTC ACTGGA GACTGGTG TCCAGT CCGATGA GTCT ACTGCCTCTGCACTATCTGCAAGCGTACCG GGAC GAAACAGC

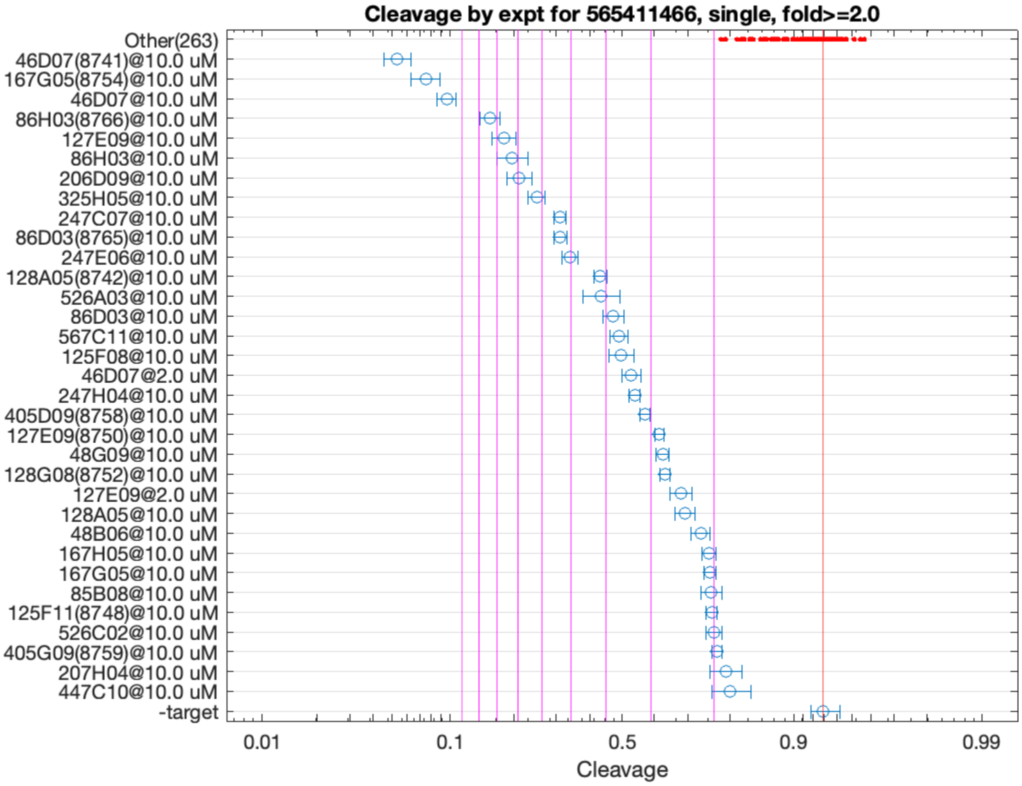


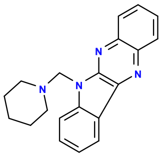

46D07

v>8.9 s=15.1

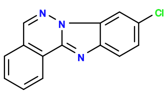

167G05

v>5.7 s=12.5

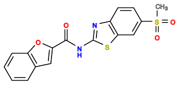

86H03

v>4.6 s=8.4

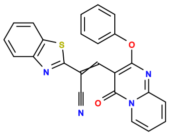

127E09

v>2.3 s=7.7

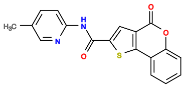

206D09

v>6.0 s=6.9

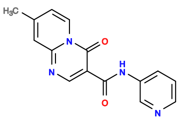

325H05

v>5.3 s=6.2

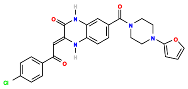

247C07

v>4.4 s=5.3

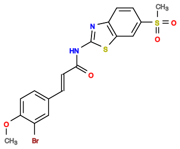

86D03

v>5.0 s=5.3

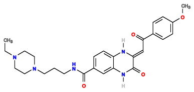

247E06

v>5.5 s=5.0

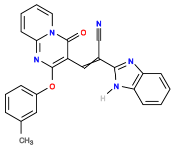

128A05

v>2.4 s=4.1

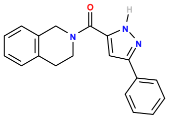

526A03

v>2.3 s=4.1

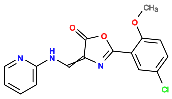

567C11

v>3.1 s=3.7

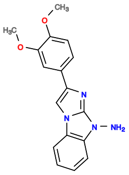

125F08

v>3.1 s=3.6

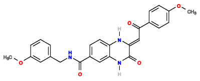

247H04

v>2.9 s=3.3

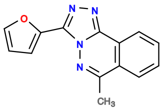

405D09

v>2.6 s=3.1

**565476652 GCTGTC ACTGGA GGAAGAGG TCCGGT CTGATGA GTCC GAAAACAAGTCAGCGCTAGGGAGATCGGTG GGAC GAAACAGC**

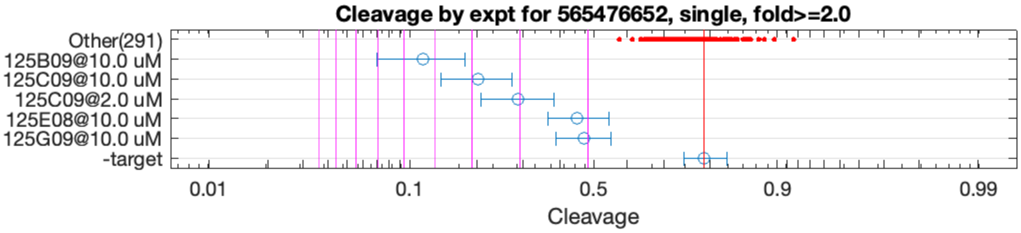


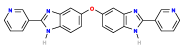

125B09

v>4.3 s=5.4

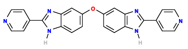

125C09

v>4.2 s=3.8

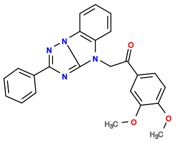

125E08

v>1.4 s=2.1

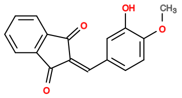

125G09

v>1.6 s=2.1

**565491775 GCTGTC ACTGGA AACTCTCTGAATGAAGCGGGTATCGCAATG TCTGGT CTGATGA GTCC GATGCGGTACAGGTCAGGGTTGGATGTG GGAC GAAACAGC**

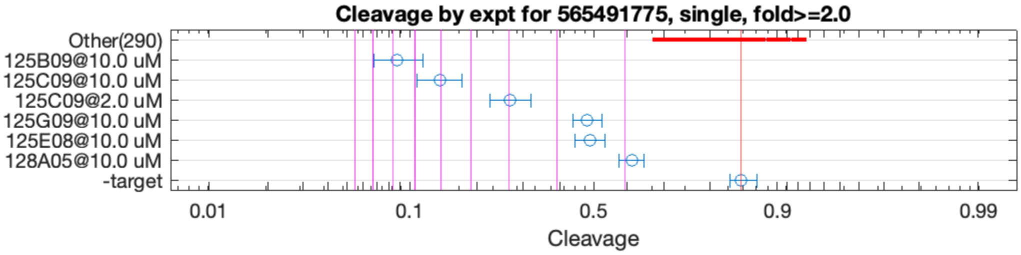


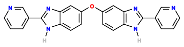

125B09

v>6.7 s=7.8

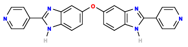

125C09

v>6.5 s=6.0

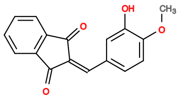

125G09

v>1.7 s=2.5

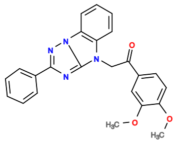

125E08

v>1.3 s=2.5

**565493161 GCTGTC ACTGGAA ATTGCAACCAAATGCAGAAAACCTCAGA TTCCGGT CCGATGA GTCCT TGGGGT GGGAC GAAACAGC**

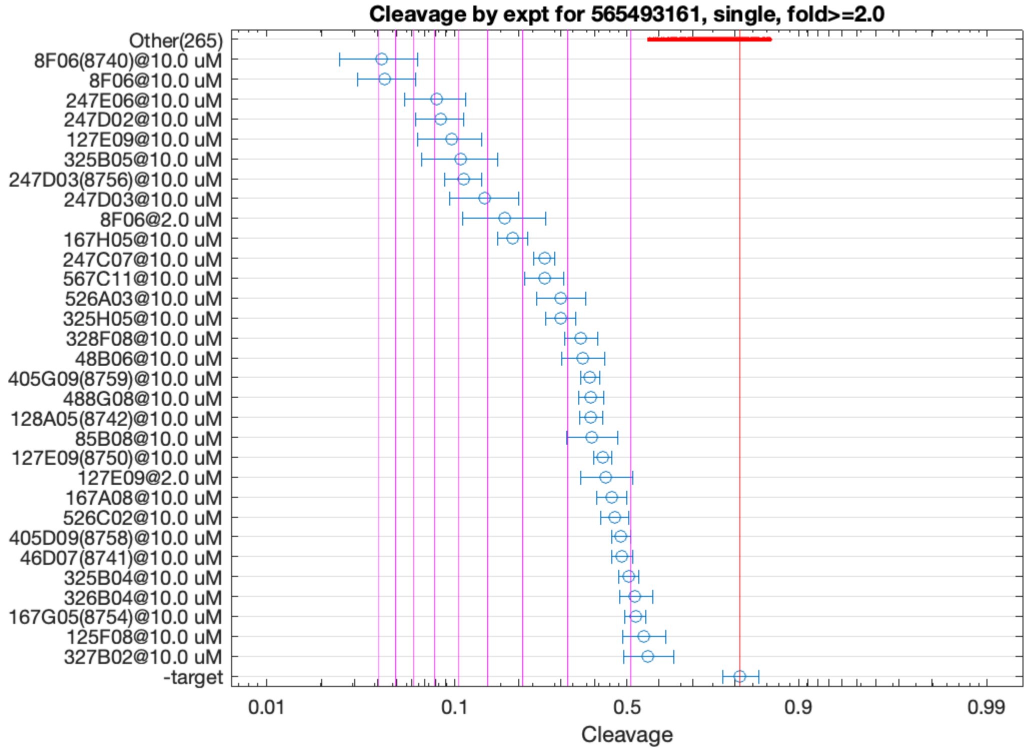


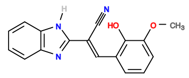

8F06

v>4.1 s=10.0

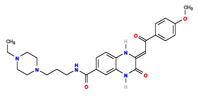

247E06

v>4.1 s=7.0

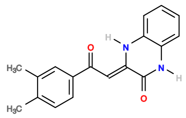

247D02

v>5.1 s=6.7

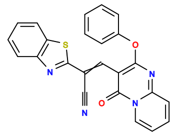

127E09

v>3.1 s=6.4

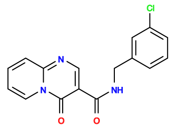

325B05

v>4.0 s=6.0

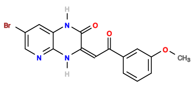

247D03

v>4.1 s=5.9

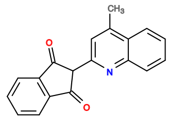

167H05

v>3.1 s=4.3

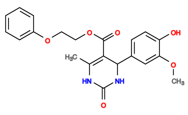

5A04

v>4.2

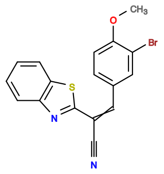

8C05

v>4.1

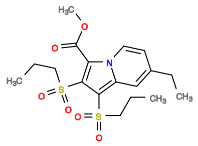

85G06

v>4.1

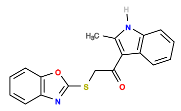

126E02

v>4.0

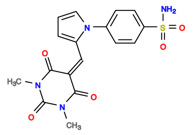

46A07

v>4.0

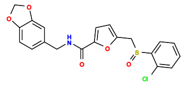

486G11

v>3.9

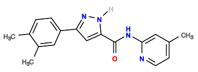

526H04

v>3.9

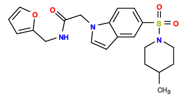

528C07

v>3.9

565513272 GCTGTC ACTGG AAAAACACTGGATGCATGACCATATTTGGTGTGCCGAGCTGCA CCAGT CTGATGA GTTC GTGGCG GGAC GAAACAGC

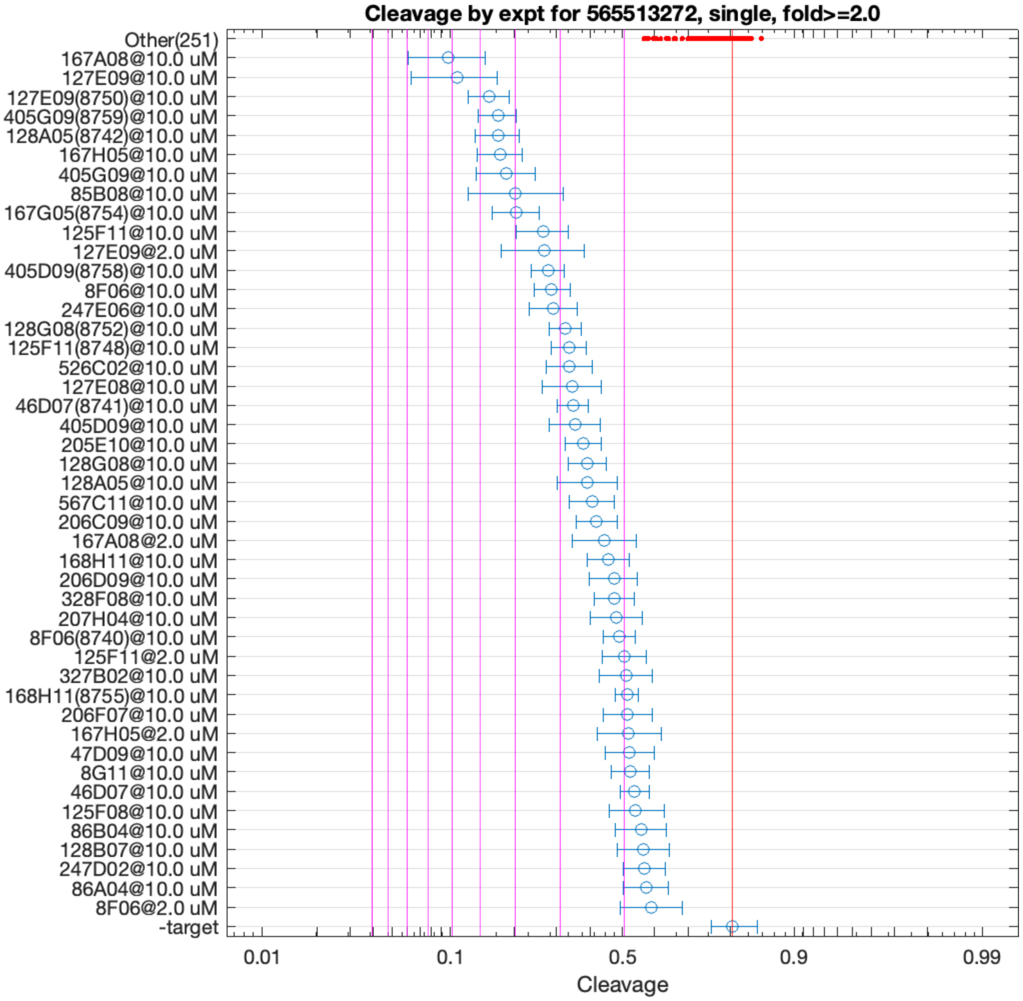


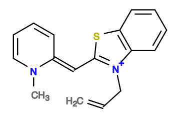

167A08

v>3.1 s=6.1

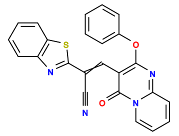

127E09

v>3.4 s=5.9

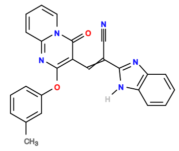

128A05

v>2.6 s=4.5

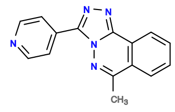

405G09

v>3.6 s=4.5

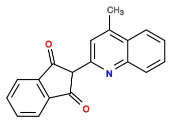

167H05

v>2.8 s=4.3

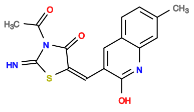

85B08

v>2.6 s=4.0

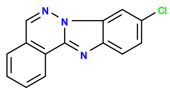

167G05

v>2.9 s=3.9

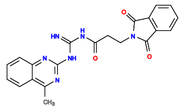

125F11

v>2.3 s=3.4

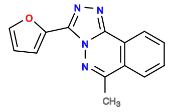

405D09

v>2.5 s=3.3

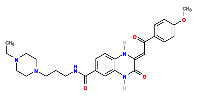

247E06

v>2.5 s=3.2

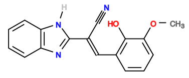

8F06

v>2.3 s=3.2

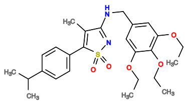

447C04

v>3.0

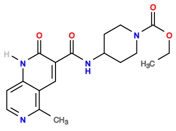

608B07

v>3.0

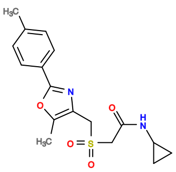

285D06

v>3.0

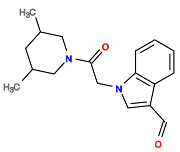

606E08

v>2.9

**565547677 GCTGTC ACTGGA GGGGACGA TCCGGT CTGATGA GTCC GTATCGAGGAAGAATTCGCGATATCCACAG GGAC GAAACAGC**

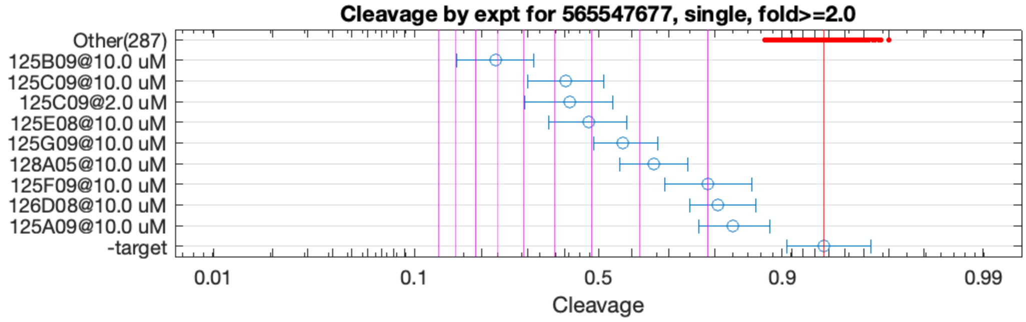


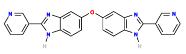

125B09

v>6.4 s=7.1

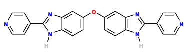

125C09

v>5.9 s=4.7

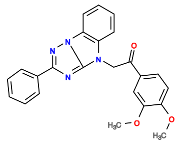

125E08

v>1.7 s=4.0

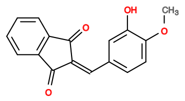

125G09

v>2.6 s=3.3

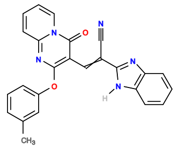

128A05

v>1.0 s=2.7

**565587233 GCTGTC ACTGGA TTTTCCTGCAATGATCTGTGGGGAAACGGC TCCGGT CTGATGA GTTC GTGGCG GGAC GAAACAGC**

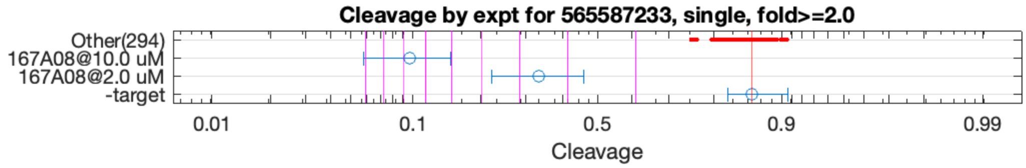


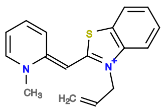

167A08

v>2.4 s=7.8

**565658419 GCTGTC ACTGGA CATTCTGAGCATTTGGGTTGAATTGGCGAA TCCGGT CTGACGA GTCCTGTA GTCGGTTACCGTGAAGCTCGGG TGTGGGAC GAAACAGC**

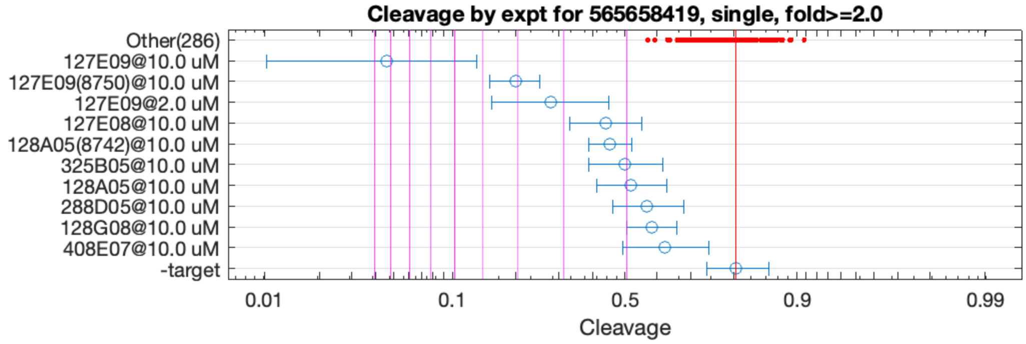


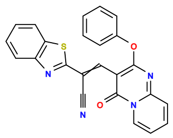

127E09

v>2.6 s=9.0

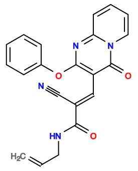

127E08

v>1.4 s=2.3

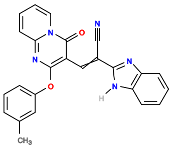

128A05

v>1.4 s=2.2

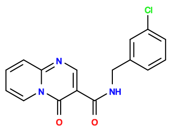

325B05

v>1.2 s=2.0

**565658927 GCTGTC ACTGGA ATTTTCTGGTTCAACTGGAATTATGCCTTCAAAGGGTGCCGATCTAA TCTGGT CTGATGA GTCT GTGGCG GGAC GAAACAGC**

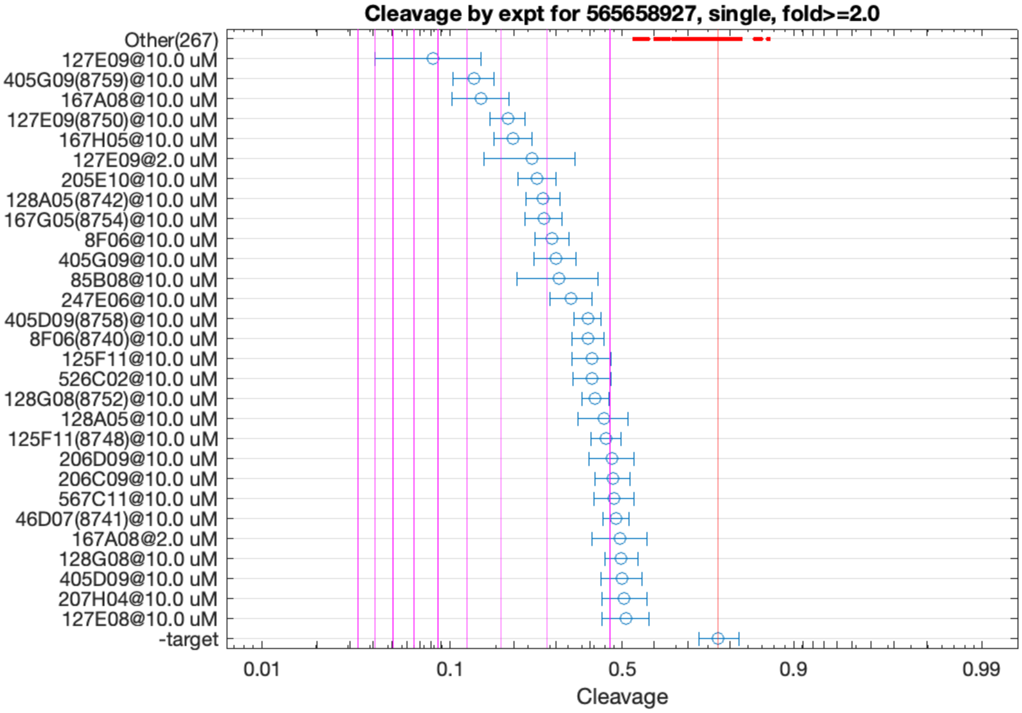


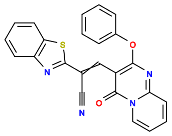

127E09

v>2.7 s=6.2

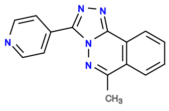

405G09

v>2.9 s=4.8

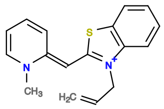

167A08

v>2.2 s=4.5

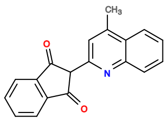

167H05

v>2.1 s=3.7

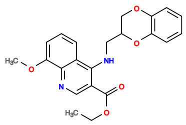

205E10

v>2.3 s=3.1

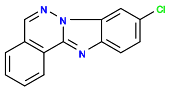

167G05

v>2.2 s=3.1

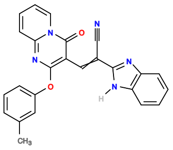

128A05

v>2.0 s=3.0

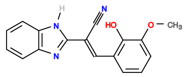

8F06

v>1.7 s=2.9

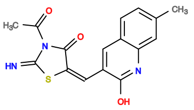

85B08

v>2.1 s=2.8

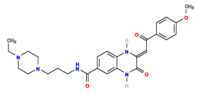

247E06

v>2.1 s=2.6

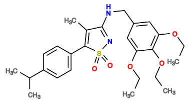

447C04

v>2.3

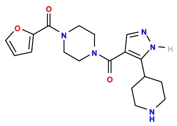

526D09

v>2.3

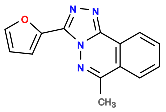

405D09

v>2.0 s=2.3

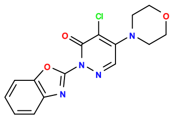

248D09

v>2.3

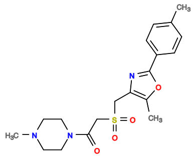

285A10

v>2.3

**565658937 GCTGTC ACTGGA ACTGGATCCAATCGCATGAGTGTGAATGGGTGTGCTGAGCTGC TCTAGT CTGACGA GTCC GTGGCG GGAC GAAACAGC**

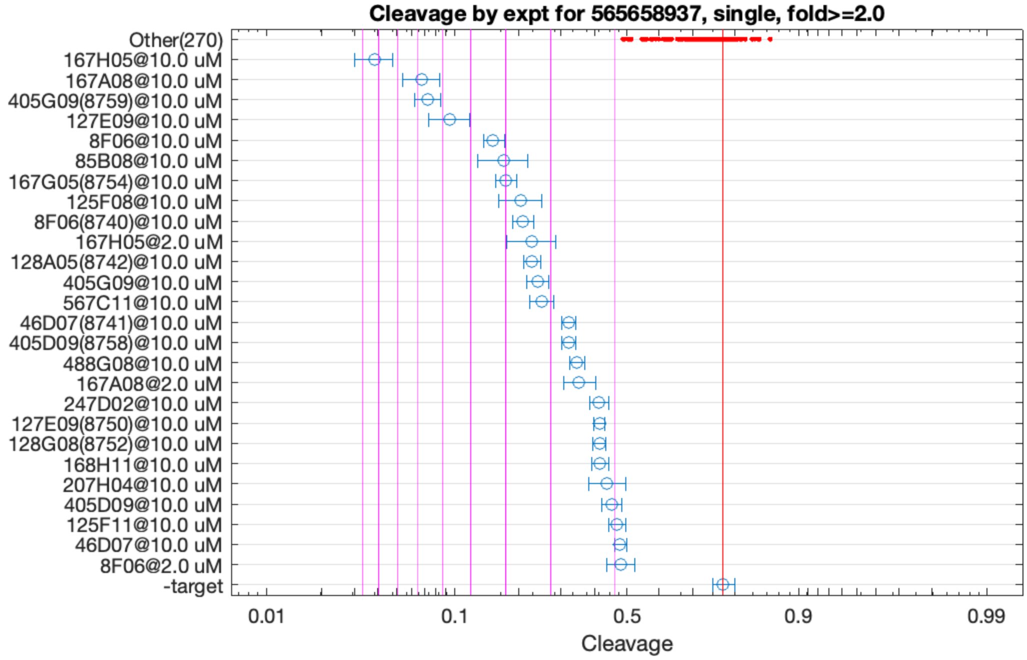


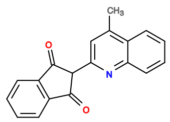

167H05

v>2.9 s=9.3

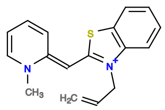

167A08

v>2.9 s=6.8

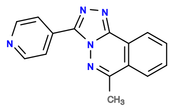

405G09

v>3.9 s=6.6

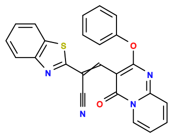

127E09

v>2.9 s=5.8

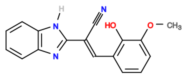

8F06

v>2.6 s=4.4

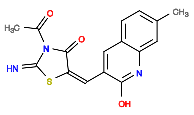

85B08

v>3.1 s=4.0

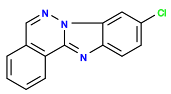

167G05

v>3.1 s=4.0

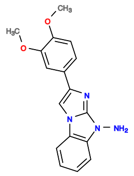

125F08

v>3.4 s=3.6

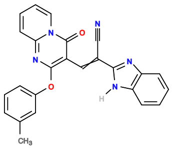

128A05

v>2.4 s=3.4

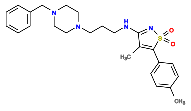

446E08

v>3.3

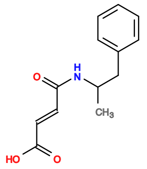

6E06

v>3.3

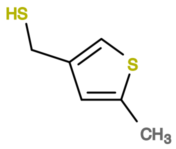

488C07

v>3.2

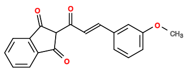

6C06

v>3.2

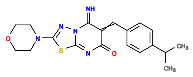

327H05

v>3.2

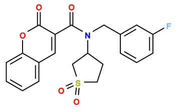

408E03

v>3.2

**565662496 GCTGTC ACTGGAC GTGGCGA GTTCGGT CTGACGA GTCC ATACATGTACTGTAAAGACAGGAGGTGGTG GGAC GAAACAGC**

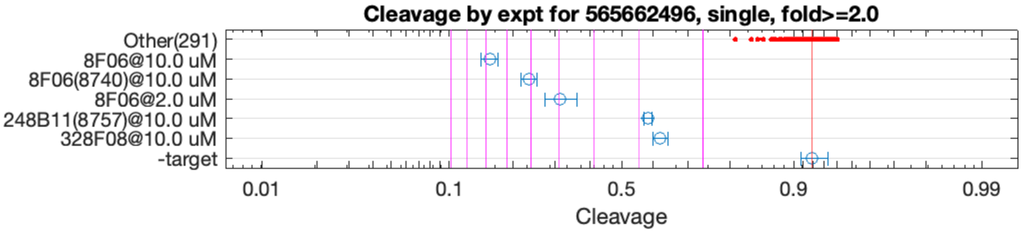


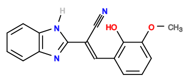

8F06

v>3.1 s=7.8

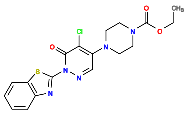

248B11

v>1.1 s=2.8

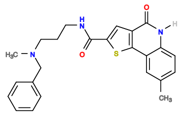

328F08

v>1.4 s=2.6

**565663129 GCTGTC ACTGGA ATTTTCTGGTTCAACTGGAATTATGCCTTTCGAGGGTTGCCGATCTAA TCTGGT CTGACGA GTCT GTGGCG GGAC GAAACAGC**

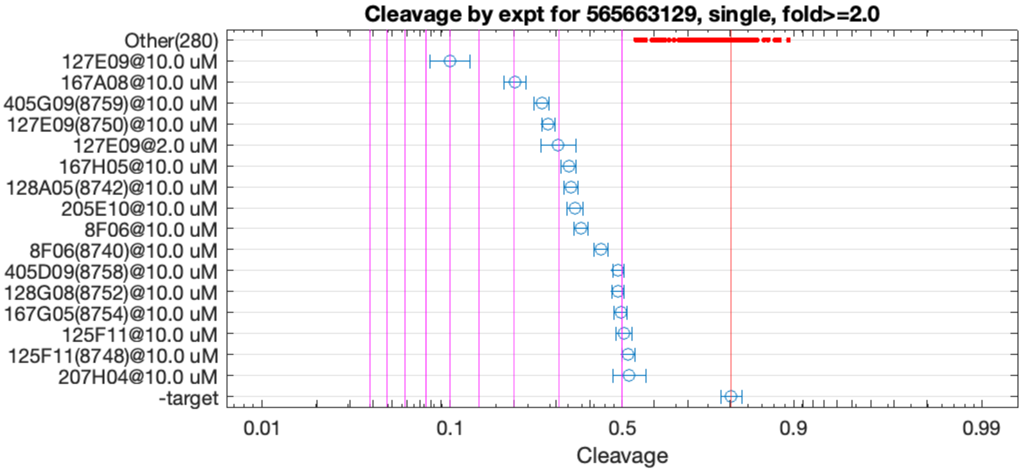


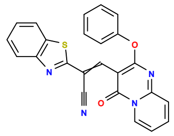

127E09

v>2.5 s=6.0

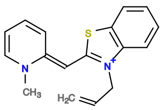

167A08

v>2.0 s=4.0

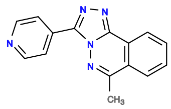

405G09

v>2.4 s=3.3

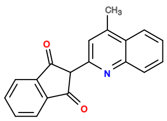

167H05

v>2.1 s=2.8

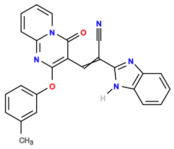

128A05

v>1.9 s=2.8

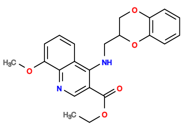

205E10

v>1.8 s=2.7

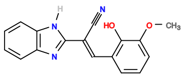

8F06

v>1.8 s=2.6

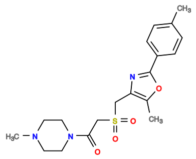

285A10

v>2.3

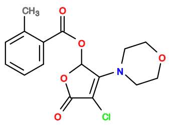

88E07

v>2.3

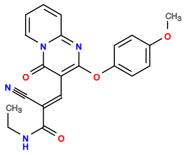

128G03

v>2.2

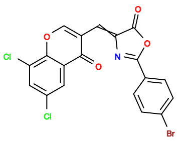

564H11

v>2.2

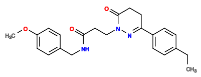

607A08

v>2.2

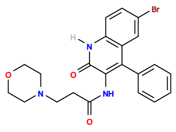

125A10

v>2.2

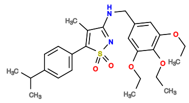

447C04

v>2.2

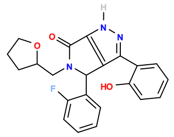

405B11

v>2.2

565675752 GCTGTC ACTG GAATGCGTAAA CAGT CTGACGA GTCC AAATCTGCTCGATGTCGTGTAGGTGCGTG GGAC GAAACAGC

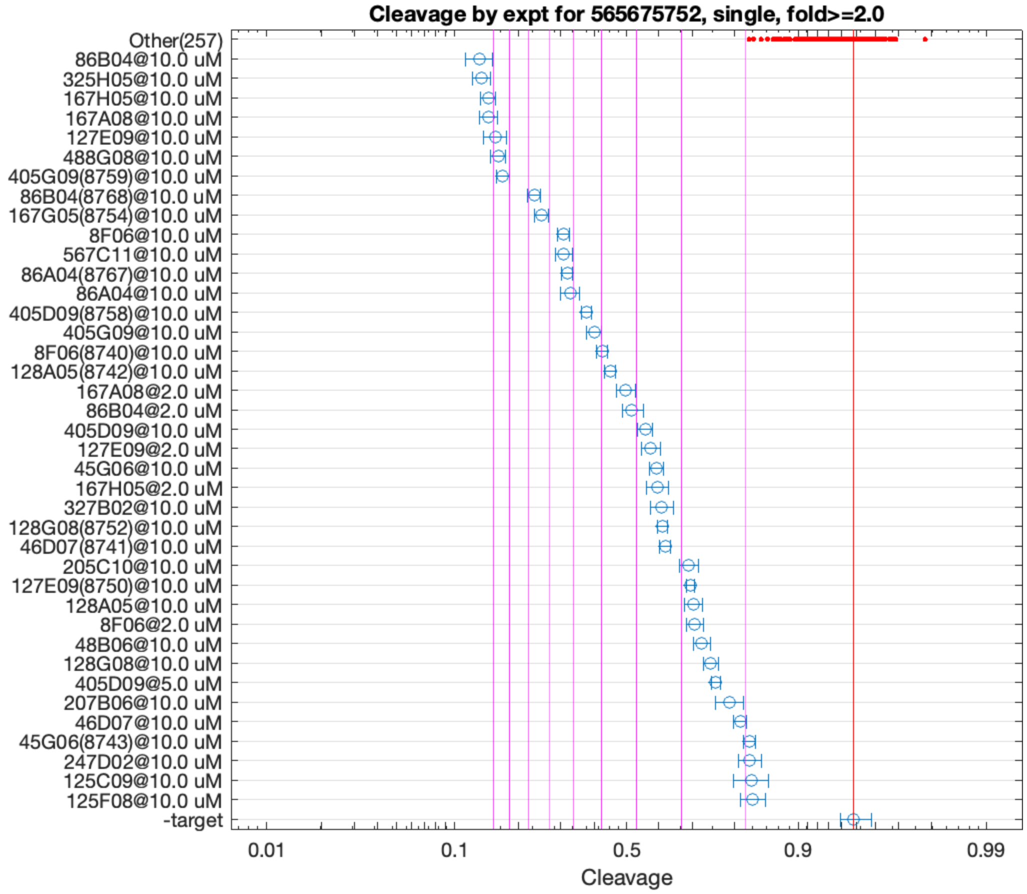


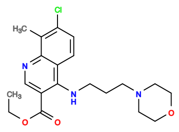

86B04

v>6.4 s=11.0

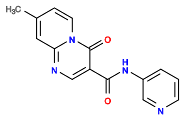

325H05

v>11.5 s=10.8

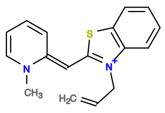

167A08

v>5.2 s=10.4

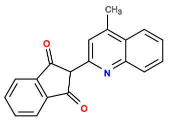

167H05

v>5.8 s=10.4

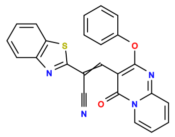

127E09

v>4.5 s=10.0

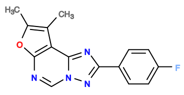

488G08

v>5.3 s=9.7

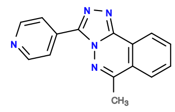

405G09

v>6.6 s=9.4

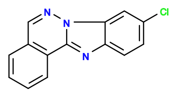

167G05

v>5.4 s=7.4

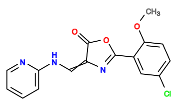

567C11

v>6.4 s=6.4

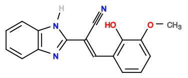

8F06

v>4.0 s=6.4

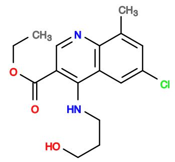

86A04

v>5.1 s=6.3

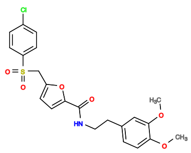

605E11

v>5.9

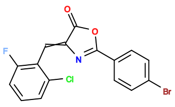

564H10

v>5.6

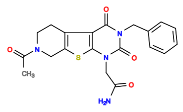

327A11

v>5.6

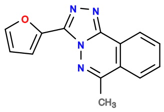

405D09

v>4.6 s=5.5

**565703067 GCTGTC ACTG GAATCCGTTAA CAGT CTGACGA GTCC AAATCTGCTCGATTTCGCGTGGGTGCGGG GGAC GAAACAGC**

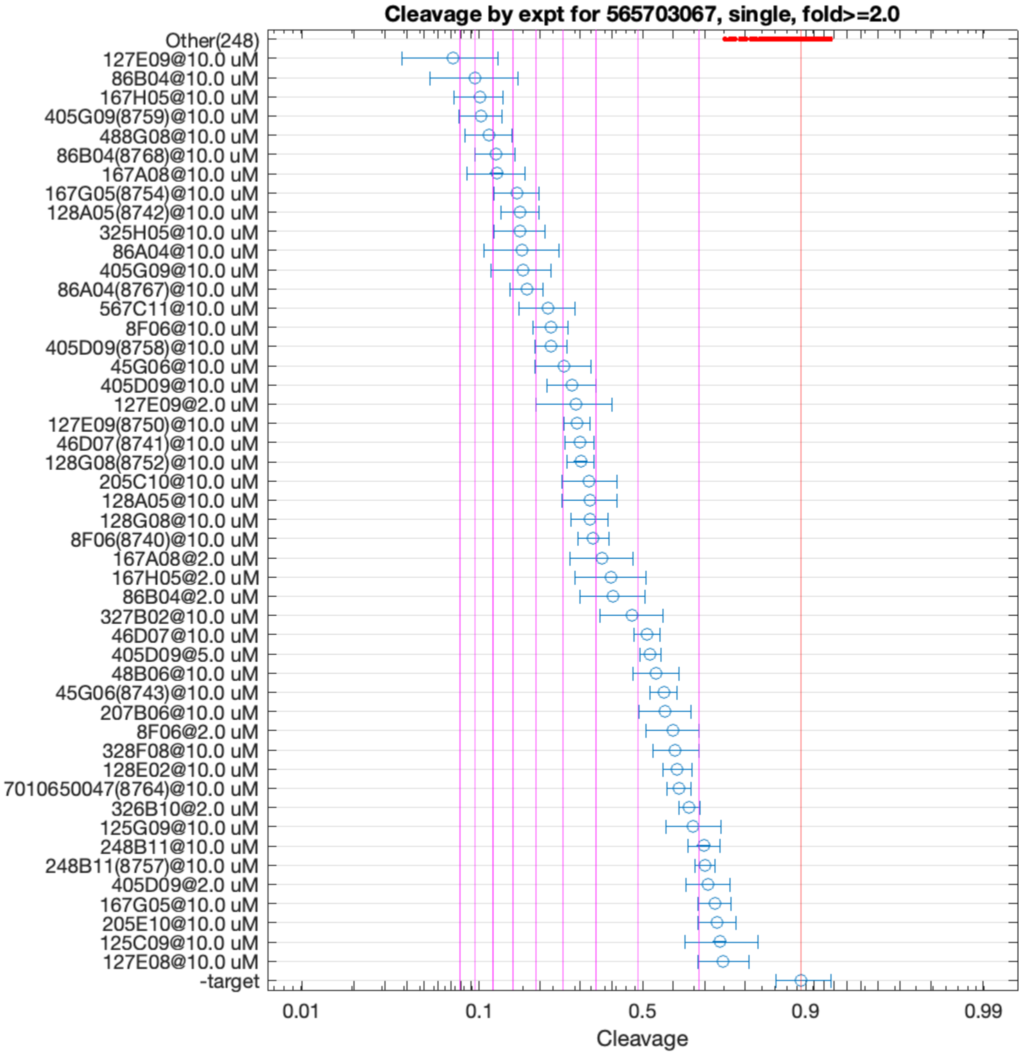


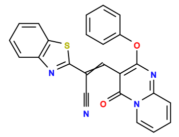

127E09

v>4.9 s=10.3

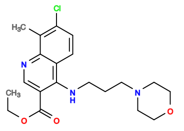

86B04

v>6.7 s=9.0

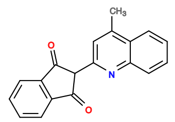

167H05

v>4.8 s=8.8

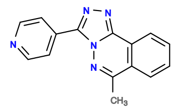

405G09

v>6.4 s=8.6

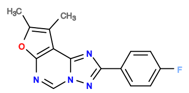

488G08

v>5.1 s=8.2

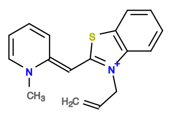

167A08

v>4.6 s=7.7

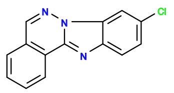

167G05

v>5.5 s=6.8

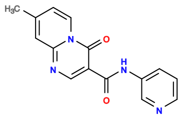

325H05

v>6.4 s=6.7

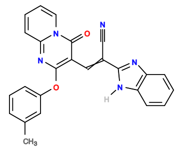

128A05

v>4.6 s=6.6

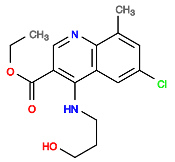

86A04

v>5.1 s=6.5

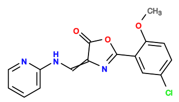

567C11

v>4.9 s=5.6

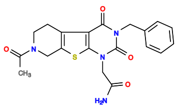

327A11

v>5.4

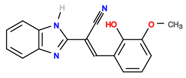

8F06

v>3.8 s=5.4

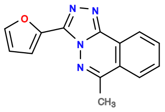

405D09

v>4.7 s=5.4

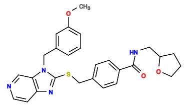

365A02

v>5.2

**565703478 GCTGTC ACTGGAA AACATTTTGCGCCGTTCTACTAGAACAC TTCCGGT CCGATGA GTCCTT GCGC GGGGAC GAAACAGC**

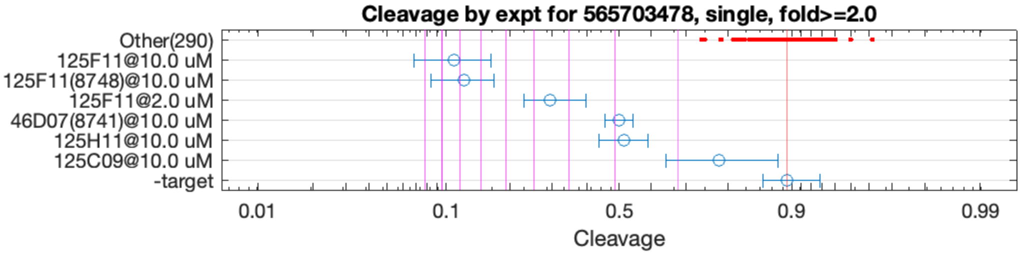


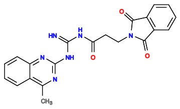

125F11

v>1.7 s=8.2

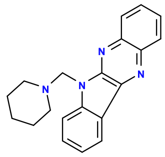

46D07

v>1.0 s=2.9

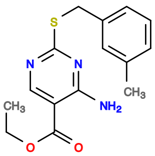

125H11

v>1.4 s=2.8

**565707921 GCTGTC ACTGG ATTTTCGACGAATGATGTTGAAAGGATGCATGACCATATTTGGTGTGCCGAGCTGCA CCAGT CTGATGA GTTC GTGGCG GGAC GAAACAGC**

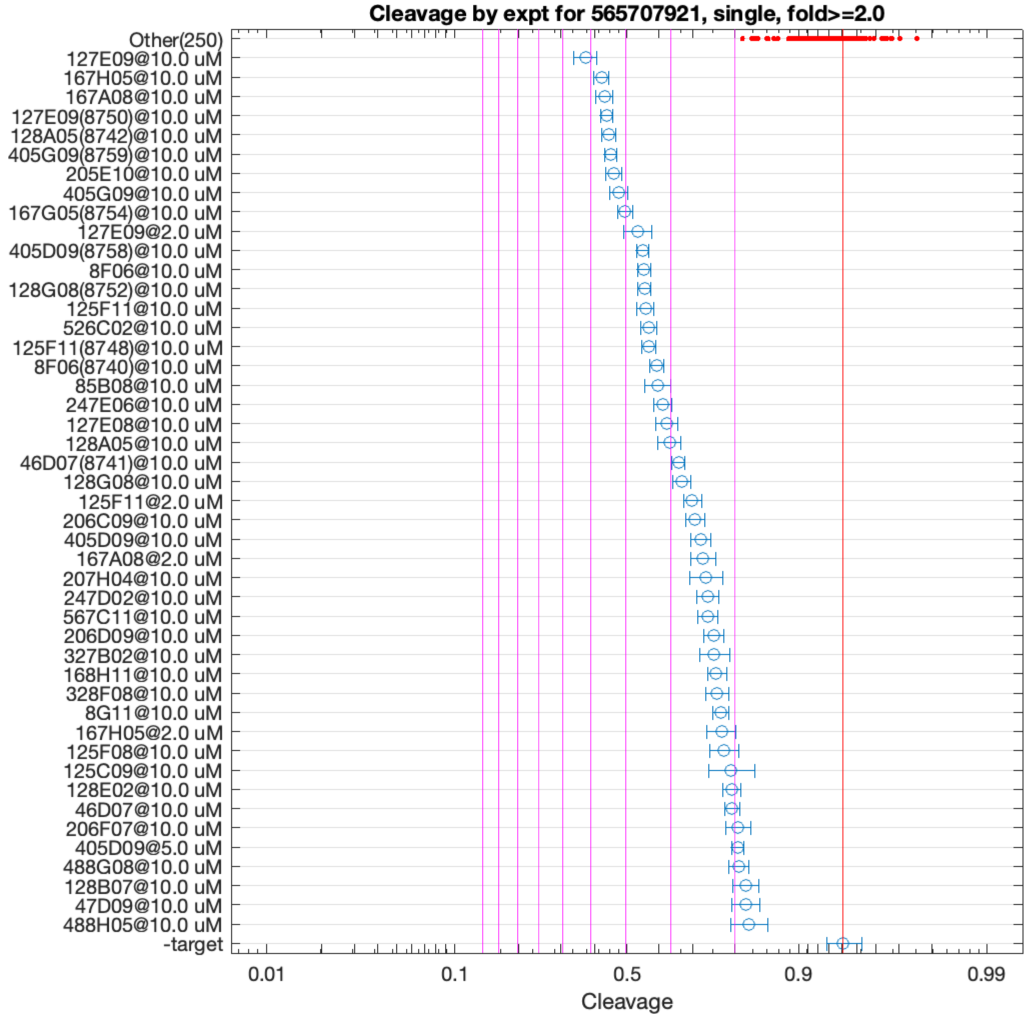


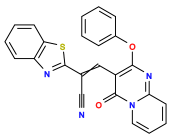

127E09

v>4.2 s=5.2

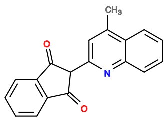

167H05

v>3.7 s=4.7

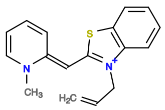

167A08

v>3.5 s=4.6

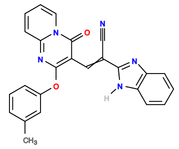

128A05

v>3.4 s=4.5

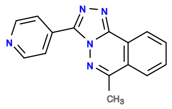

405G09

v>4.4 s=4.4

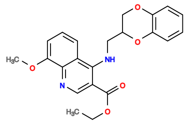

205E10

v>3.9 s=4.3

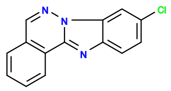

167G05

v>3.6 s=4.0

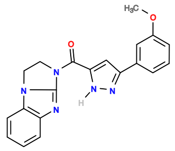

525D11

v>3.9

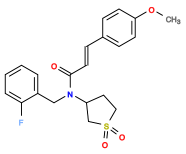

407E11

v>3.9

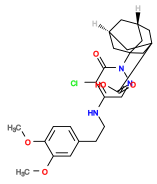

247F08

v>3.9

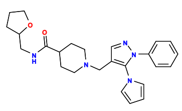

446C03

v>3.8

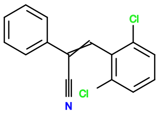

8G09

v>3.8

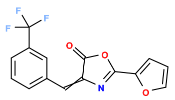

564A05

v>3.8

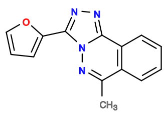

405D09

v>3.4 s=3.6

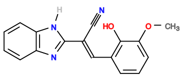

8F06

v>3.0 s=3.6

**565730172 GCTGTC ACTGGAA AAAGG TTTCGGT CTGATGA GTTC GATTGATGACGAAACAGCATTGTGCAGCCCG GGAC GAAACAGC**

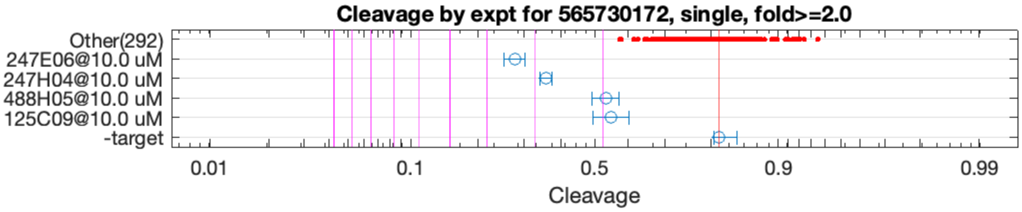


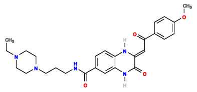

247E06

v>3.6 s=3.4

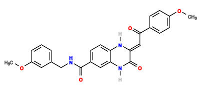

247H04

v>1.3 s=2.8

**565736657 GCTGTC ACTGGA AAAAGGACG TCTAGT CTGATGA GTCT ACTGCCTCTGCCCTATCTGCAAGCGTACCG GGAC GAAACAGC**

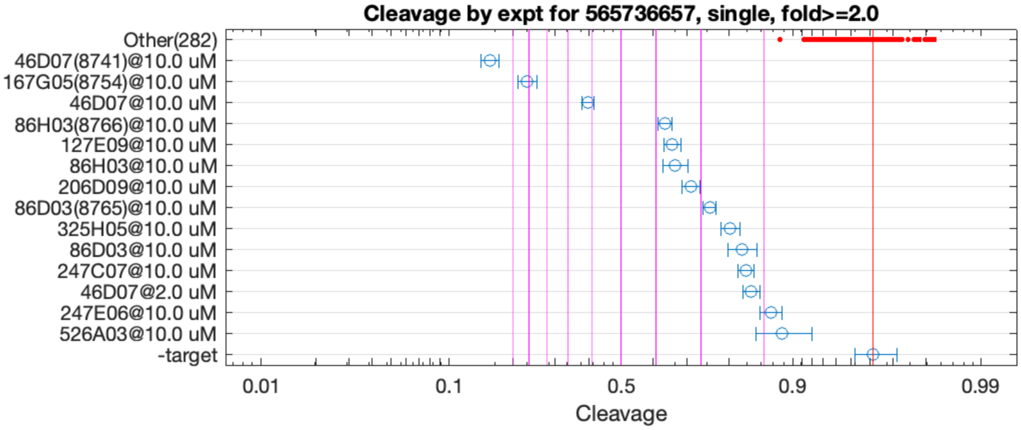


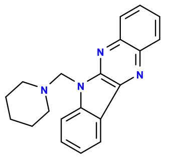

46D07

v>5.0 s=11.6

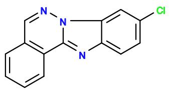

167G05

v>2.4 s=9.0

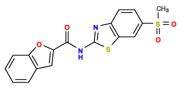

86H03

v>2.0 s=3.8

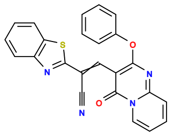

127E09

v>1.5 s=3.6

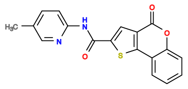

206D09

v>2.6 s=3.2

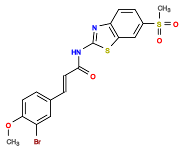

86D03

v>2.9 s=2.8

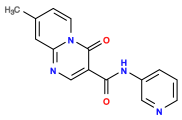

325H05

v>2.2 s=2.5

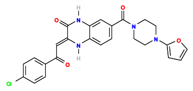

247C07

v>1.9 s=2.2

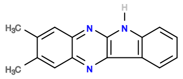

568D02

v>2.0

**565770089 GCTGTC ACTGGAT GGT GTCCGGT CCGATGA GTCC GTGTCGAAAAAGTTGATTGCTTCACACCCA GGAC GAAACAGC**

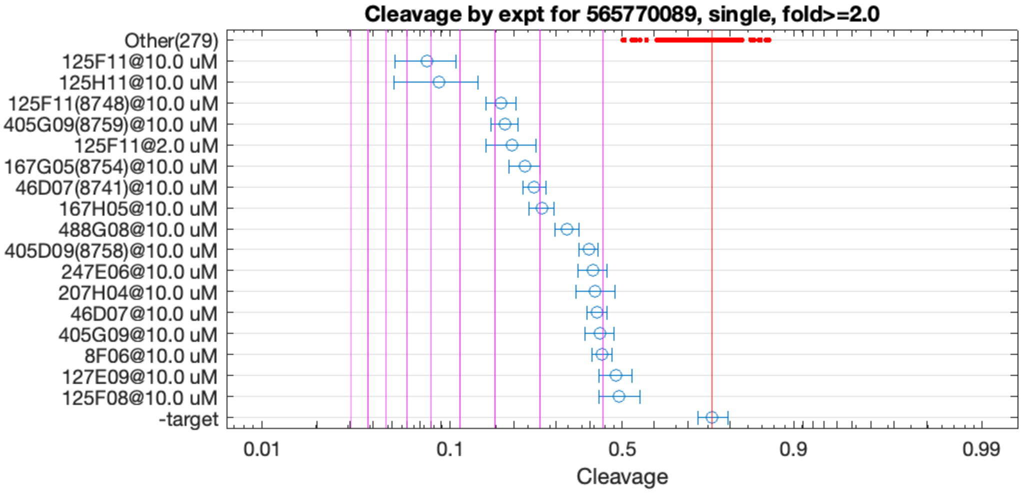


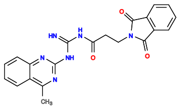

125F11

v>4.2 s=6.2

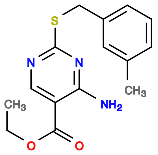

125H11

v>3.7 s=5.7

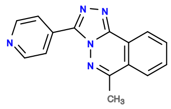

405G09

v>1.9 s=3.7

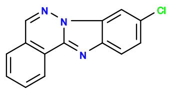

167G05

v>1.8 s=3.3

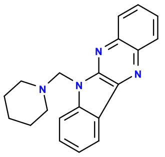

46D07

v>1.8 s=3.1

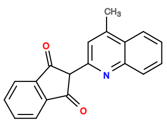

167H05

v>1.7 s=3.0

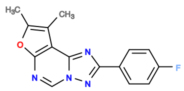

488G08

v>1.5 s=2.6

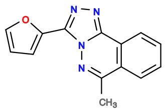

405D09

v>1.6 s=2.2

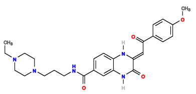

247E06

v>1.4 s=2.1

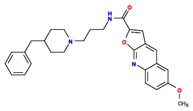

207H04

v>1.4 s=2.1

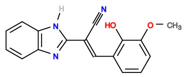

8F06

v>1.5 s=2.0

565872930 GCTGTC ACTG GAAAACGTAAA CAGT CTGAAGA GTCC AAATCTGCCGAGTTCGTGTGGGTGCGTG GGAC GAAACAGC

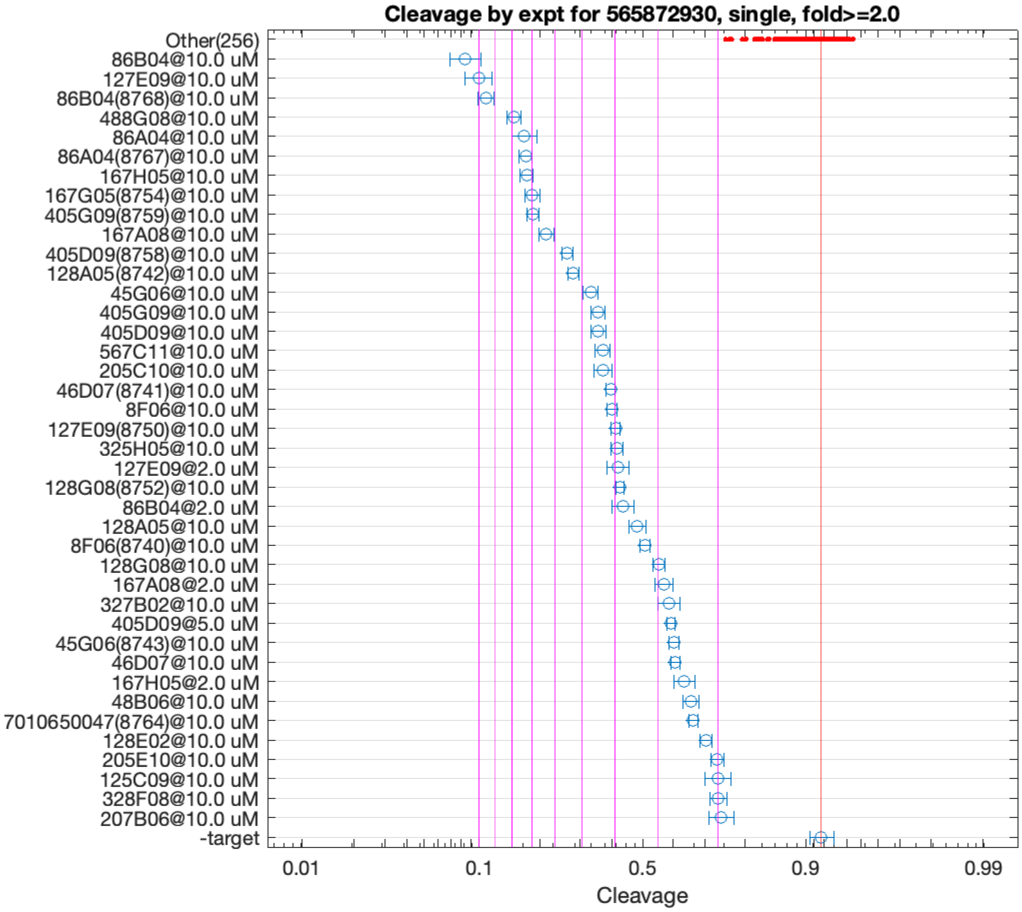


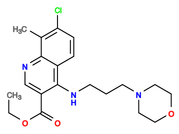

86B04

v>8.3 s=11.0

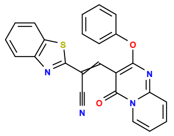

127E09

v>5.4 s=10.0

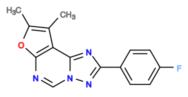

488G08

v>5.1 s=7.9

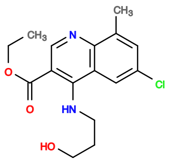

86A04

v>6.1 s=7.4

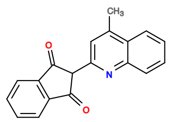

167H05

v>4.8 s=7.2

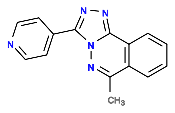

405G09

v>5.7 s=7.0

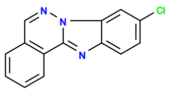

167G05

v>5.7 s=7.0

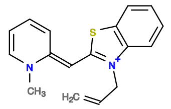

167A08

v>4.0 s=6.4

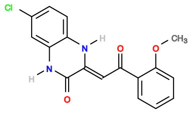

247C02

v>5.6

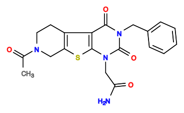

327A11

v>5.6

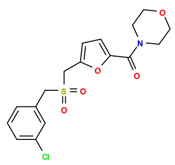

607A02

v>5.6

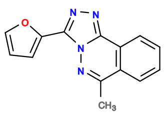

405D09

v>4.7 s=5.5

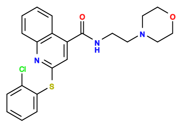

208H11

v>5.4

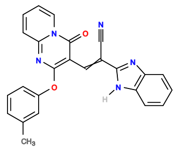

128A05

v>4.3 s=5.3

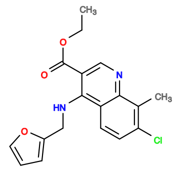

45G06

v>5.4 s=4.7

**565928760 GCTGTC ACTGGA AAAAGGTTGCAATTCAAATGTCACTGGACTTC TCTGGT CTGATGA GTC AGAATGCTGACGAAACGGCTCTTGTTGTCGCG GAC GAAACAGC**

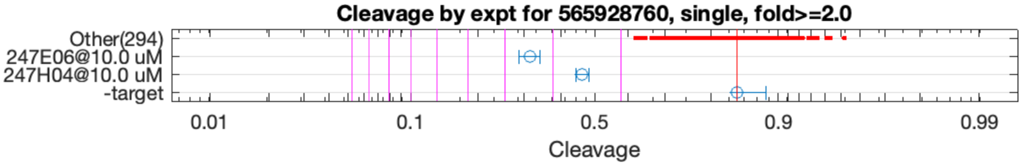


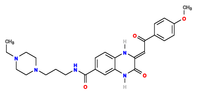

247E06

v>3.9 s=3.4

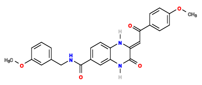

247H04

v>1.1 s=2.5

**565957580 GCTGTC ACTGGAAT AAG GTTTCGGT CTGATGA GTCC GATGACGAAACAGCGAGTTGTGCGAACCCG GGAC GAAACAGC**

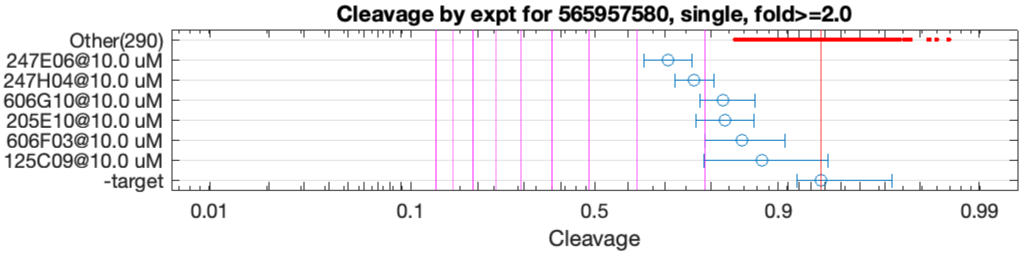


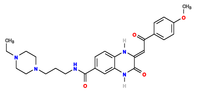

247E06

v>3.2 s=2.5

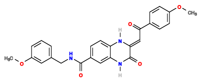

247H04

v>0.8 s=2.1

**565958337 GCTGTC ACTGGA CACTACT TCCGGT CTGATGA GTCC GGGATACGACAAGTATGCCATTAGCAGGTG GGAC GAAACAGC**

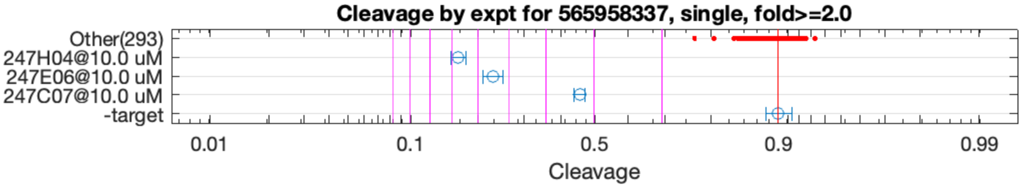


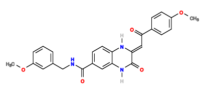

247H04

v>1.2 s=6.8

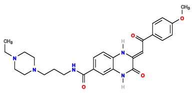

247E06

v>8.2 s=5.5

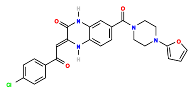

247C07

v>1.7 s=3.3

**566071971 GCTGTC ACTGGAA CGTAGGTACTAGCGGAGTCAACACAATA TTCCGGT CTGACGA GTCCTG ATATCGCAAAGCTGGCACATACTACG TGGGAC GAAACAGC**

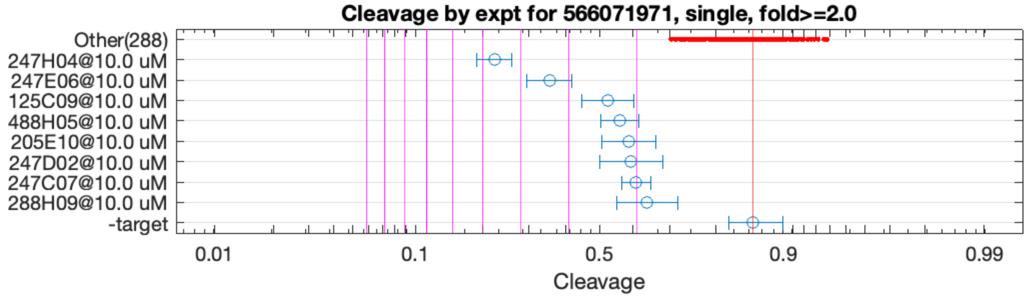


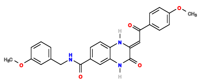

247H04

v>1.5 s=4.6

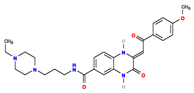

247E06

v>3.7 s=3.4

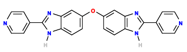

125C09

v>1.2 s=2.4

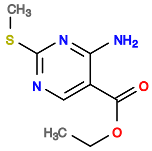

488H05

v>1.2 s=2.2

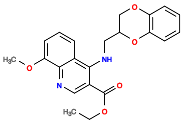

205E10

v>1.1 s=2.1

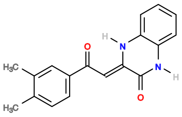

247D02

v>1.2 s=2.1

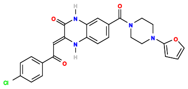

247C07

v>1.4 s=2.0

**566087733 GCTGTC ACTGG ATTTATGCCGTTGAAAGGGGTGCCGATCTAAA CTAGT CTGAAGA GTCT GTGGCG GGAC GAAACAGC**

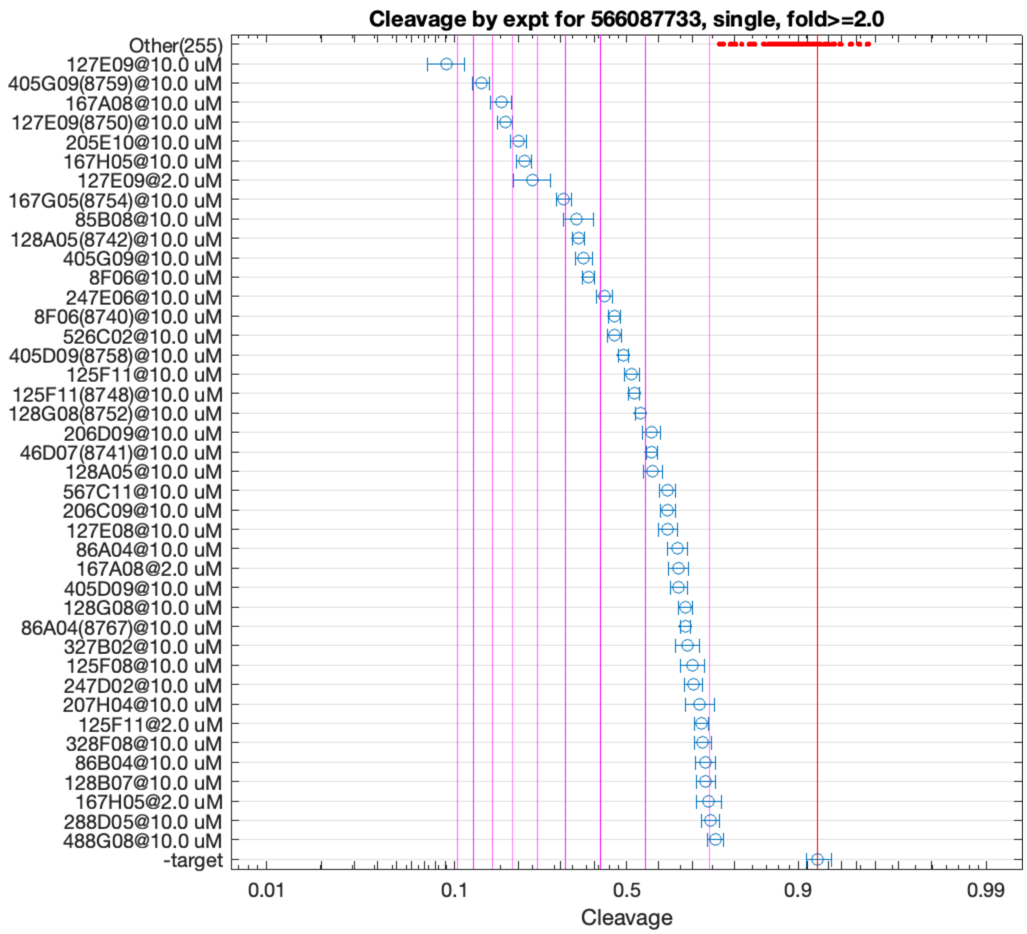


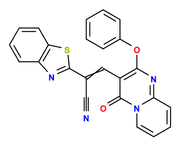

127E09

v>4.4 s=10.7

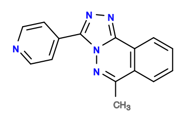

405G09

v>4.8 s=8.6

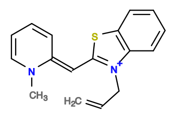

167A08

v>3.3 s=7.5

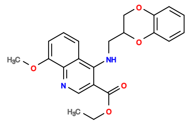

205E10

v>3.5 s=6.8

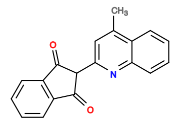

167H05

v>3.1 s=6.5

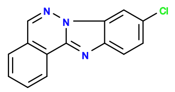

167G05

v>3.6 s=5.0

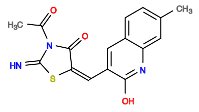

85B08

v>3.3 s=4.7

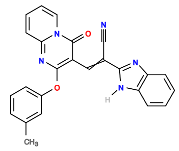

128A05

v>2.9 s=4.6

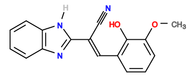

8F06

v>2.9 s=4.3

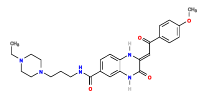

247E06

v>3.1 s=3.9

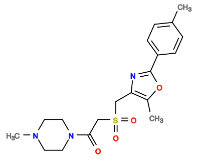

285A10

v>3.7

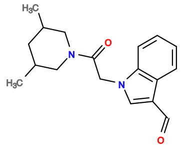

606E08

v>3.7

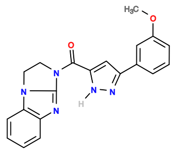

525D11

v>3.7

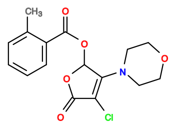

88E07

v>3.7

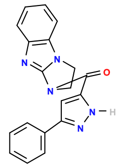

526C02

v>3.7 s=3.7

**566087783 GCTGTC ACTGGA TTTTCCTGCAATGGTCTGTGGGGAAACGGC TCTGGT CTGATGA GTTC GTGGCG GGAC GAAACAGC**

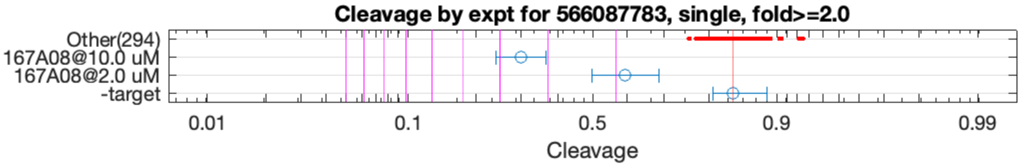


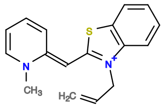

167A08

v>1.0 s=3.5

**566218011 GCTGTC ACTGGA AGGGGACGA TCCGGT CTGATGA GTCC GTATCGAGGAAGAATTCGCGATATCCACAG GGAC GAAACAGC**

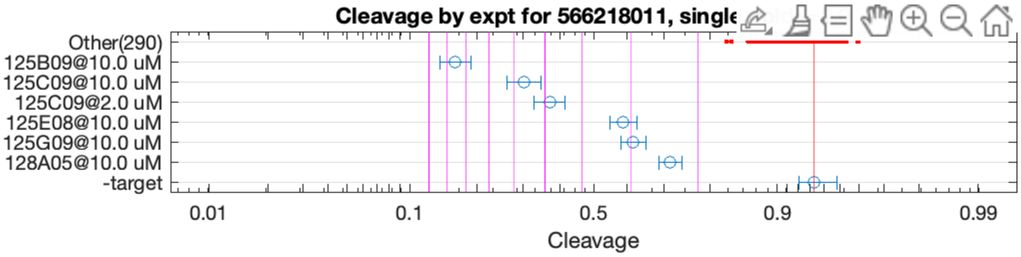


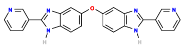

125B09

v>7.2 s=8.7

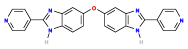

125C09

v>7.1 s=5.6

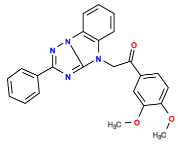

125E08

v>1.4 s=3.1

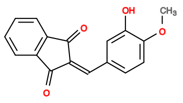

125G09

v>2.0 s=3.0

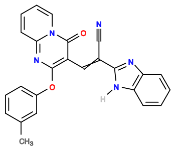

128A05

v>1.1 s=2.4

**566222400 GCTGTC ACTGGA GGAAGATG TCCGGT CTGATGA GTCC GAAAATGTTCGCGTAGACAGGAGTTCGGTG GGAC GAAACAGC**

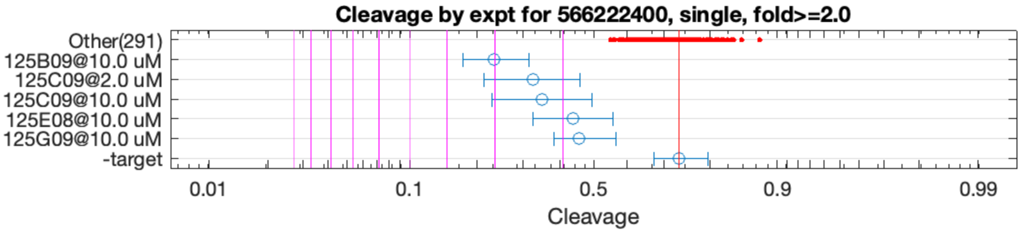


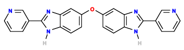

125B09

v>2.9 s=3.0

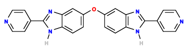

125C09

v>2.4 s=2.4

**566225843 GCTGTC ACTGGA TTCTCTTTTGGTTTGCAAATGGGAACGGC TCCGGT CTGATGA GTTC GTGGCG GGAC GAAACAGC**

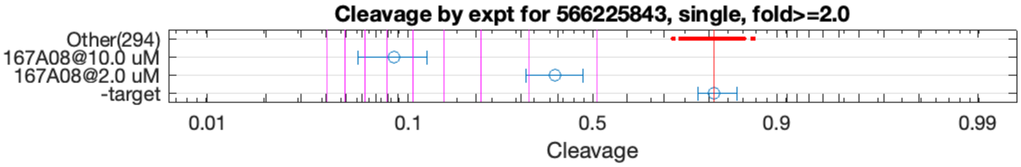


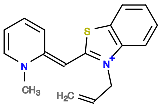

167A08

v>1.9 s=6.6

**566226123 GCTGTC ACTGGA TTGTCTTTCGGAATTACTGGAAGATGACGGC TCCGGT CTGATGA GTTC GTGGCG GGAC GAAACAGC**

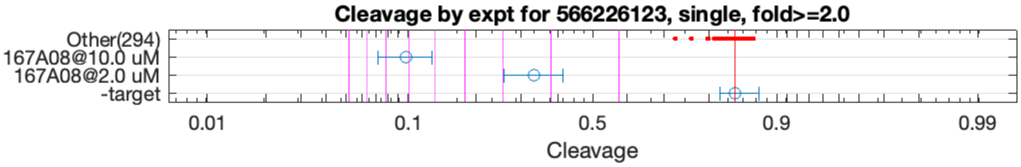


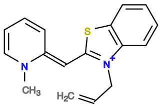

167A08

v>2.3 s=7.1

**566228712 GCTGTC ACTGGA TTTCGGTTAGCAATGCTACTGGAACGGC TCCGGT CTGATGA GTTC GTGGCG GGAC GAAACAGC**

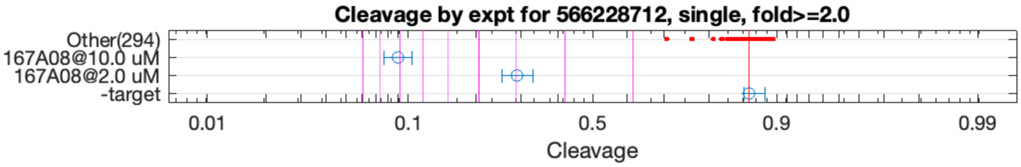


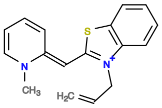

167A08

v>2.9 s=8.2

**566230038 GCTGTC ACTGGA TTTCTTGTTGGTAGCAGCAGGAAACGGC TCCGGT CTGAAGA GTTC GTGGCG GGAC GAAACAGC**

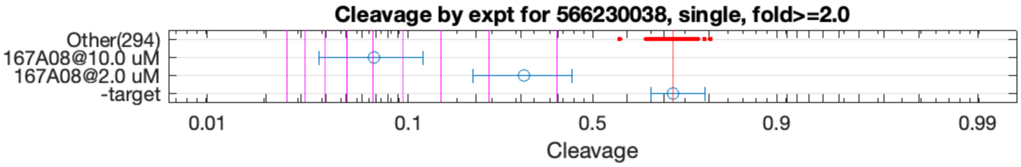


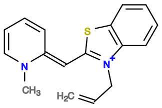

167A08

v>1.7 s=6.1

**566264584 GCTGTC ACTGGA TATCACCGCAGAAATCAAGCCATTTCGAAC TCCGGT CTGATGA GTTC ACGTTCGGGATGAGCGCCTGGGGATGGGTA GGAC GAAACAGC**

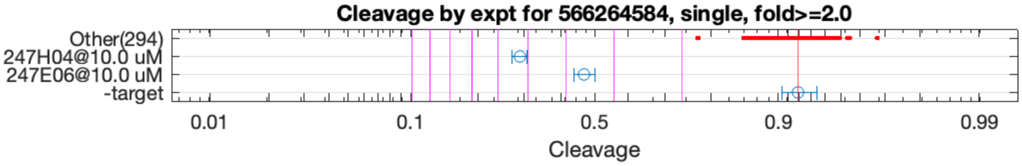


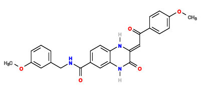

247H04

v>1.2 s=5.2

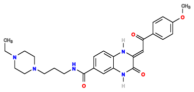

247E06

v>5.5 s=3.6

**566297247 GCTGTC ACTGGA ATCCAAGCCCTATGAGCAACATGGTATACC TCCGGT CTGATGA GTTCT AGTCGCCATGTAGCACGGTAGCTGAGGA GGGAC GAAACAGC**

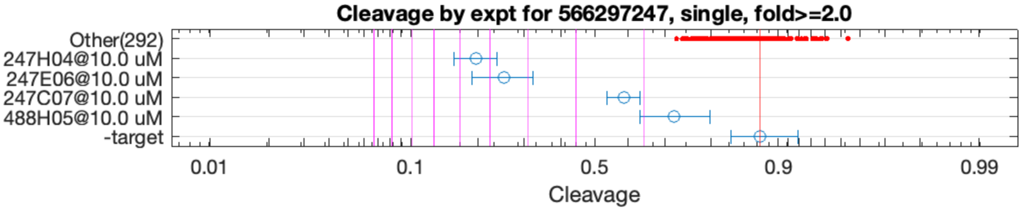


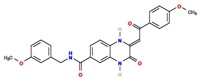

247H04

v>1.4 s=5.4

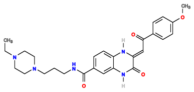

247E06

v>5.2 s=4.5

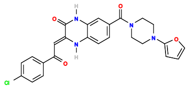

247C07

v>1.4 s=2.2

**566304100 GCTGTC ACTGGAAT AAG GTTTCGGT CTGATGA GTCC GATGACGAAACAGCGAGTTGCGCGAACCCG GGAC GAAACAGC**

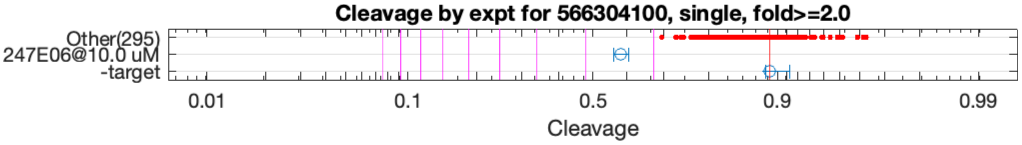


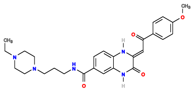

247E06

v>3.0 s=2.4

**566314690 GCTGTC ACTGGAT GGT GTCCGGT CCGATGA GTCC GTGTCGAAAAAGTTGATTACTTCACACCCA GGAC GAAACAGC**

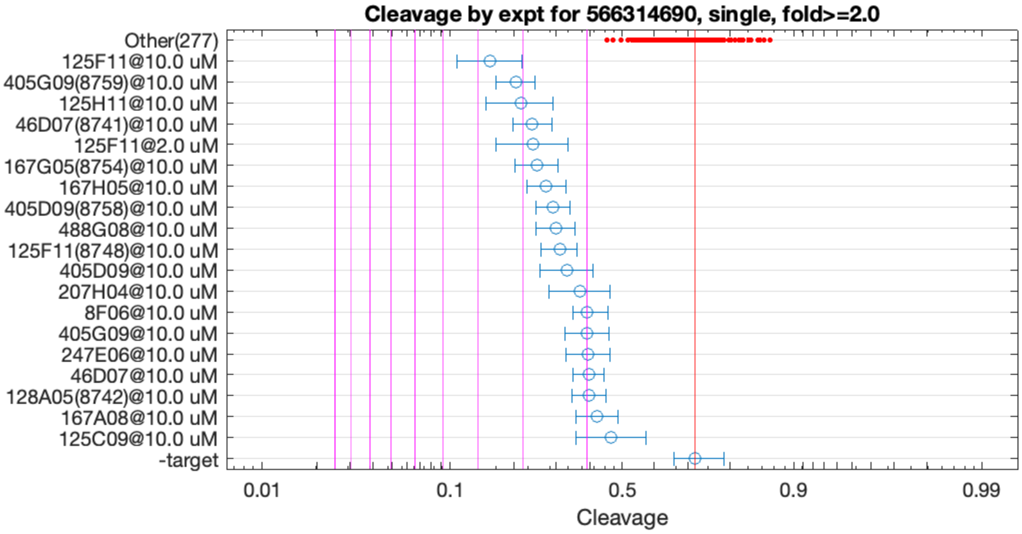


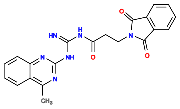

125F11

v>2.8 s=3.7

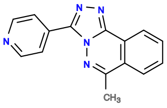

405G09

v>1.8 s=3.2

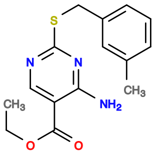

125H11

v>2.5 s=3.0

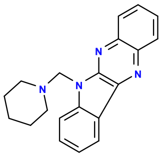

46D07

v>1.6 s=2.8

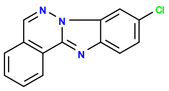

167G05

v>1.7 s=2.8

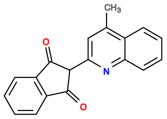

167H05

v>1.5 s=2.6

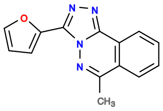

405D09

v>1.5 s=2.5

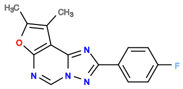

488G08

v>1.4 s=2.4

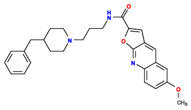

207H04

v>1.6 s=2.0

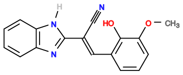

8F06

v>1.4 s=2.0

**566334018 GCTGTC ACTGGA TTTTGCGCCGTTCTACTTGAACACT TCCGGT CCGATGA GTCCTT GCGC GGGGAC GAAACAGC**

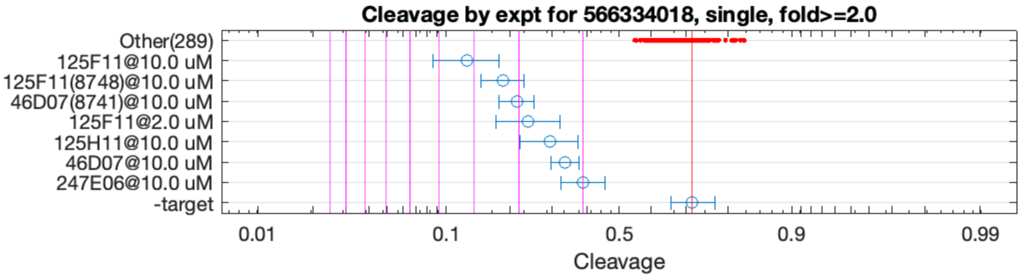


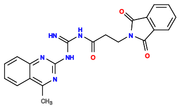

125F11

v>2.6 s=4.2

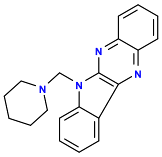

46D07

v>1.6 s=3.0

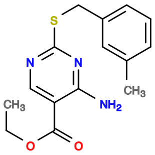

125H11

v>1.9 s=2.5

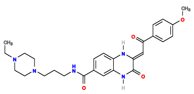

247E06

v>1.2 s=2.0

**566566059 GCTGTC ACTGGAG CTGGACA CTCTGGT CTGATGA GTCC ATGAAGAGTGTGCGCAGACAAACGCACGGG GGAC GAAACAGC**

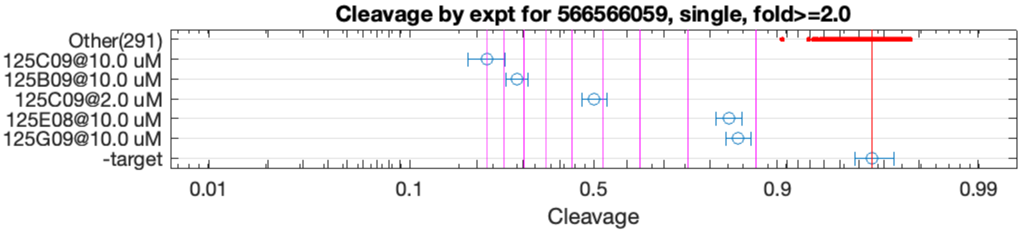


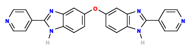

125C09

v>9.9 s=10.1

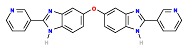

125B09

v>7.7 s=8.3

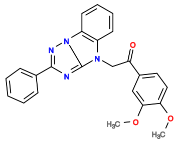

125E08

v>1.5 s=2.4

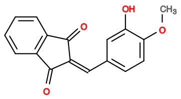

125G09

v>1.9 s=2.2

**566568470 GCTGTC ACTGGAC GTGGCGA GTTCGGT CTGATGA GTCC AAAAGGAAGAGTGTGTGTGAGCACAGAGGC GGAC GAAACAGC**

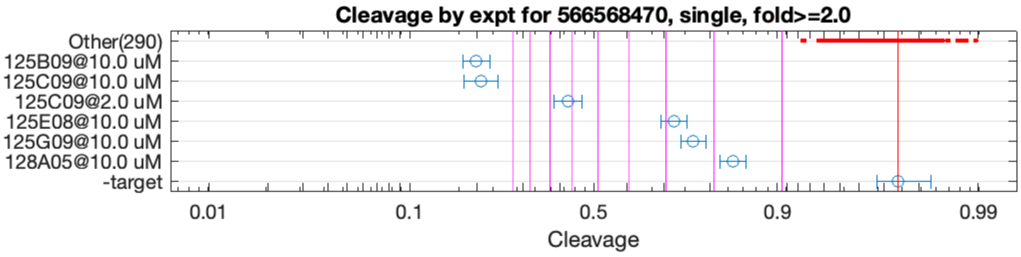


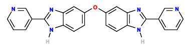

125B09

v>12.1 s=12.3

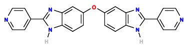

125C09

v>13.6 s=12.0

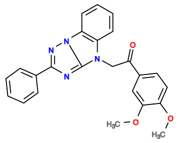

125E08

v>2.0 s=3.9

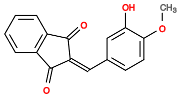

125G09

v>3.2 s=3.4

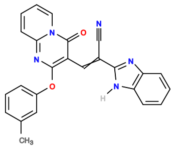

128A05

v>1.4 s=2.7

**566652019 GCTGTC ACTGGA TACGATTTGGTGATTTCGCGCATGCCAAAC TCCGGT CTGACGA GTCT CAGTTTCGCATGCCCTTGCCAATGAAGTA GGAC GAAACAGC**

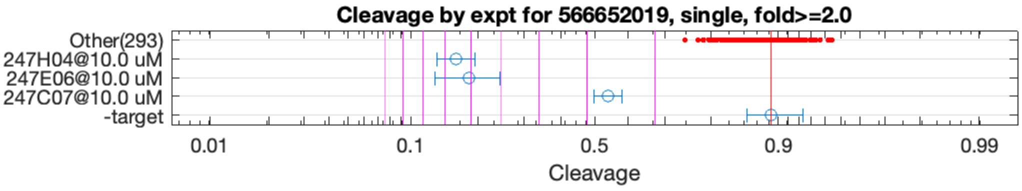


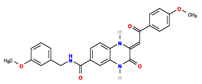

247H04

v>1.4 s=6.5

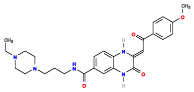

247E06

v>6.8 s=6.0

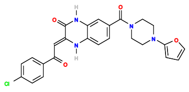

247C07

v>1.7 s=2.7

**566655862 GCTGTC ACTGGAT GGT GTCCGGT CCGATGA GTCC GTGTCGAAAAGGTTGATTGCTTCACACCCA GGAC GAAACAGC**

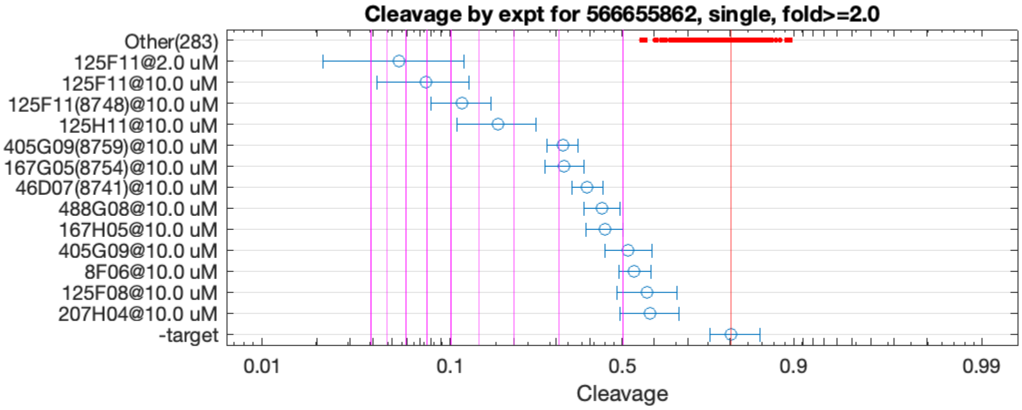


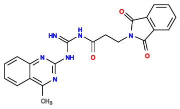

125F11

v>4.0 s=8.5

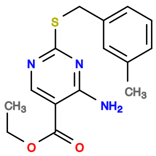

125H11

v>2.8 s=4.4

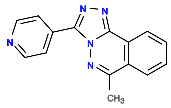

405G09

v>1.4 s=2.9

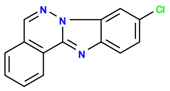

167G05

v>1.5 s=2.9

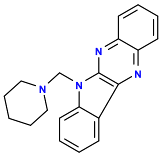

46D07

v>1.3 s=2.5

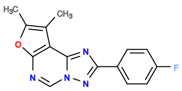

488G08

v>1.2 s=2.3

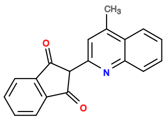

167H05

v>1.3 s=2.2

**566669595 GCTGTC ACTGGA CACTCCGTGGGCATTTGGGTTGAATTGGCGAA TCCGGT CTGACGA GTCCTGTA GTCGGTTACCGTGCAGCTCAGG TGTGGGAC GAAACAGC**

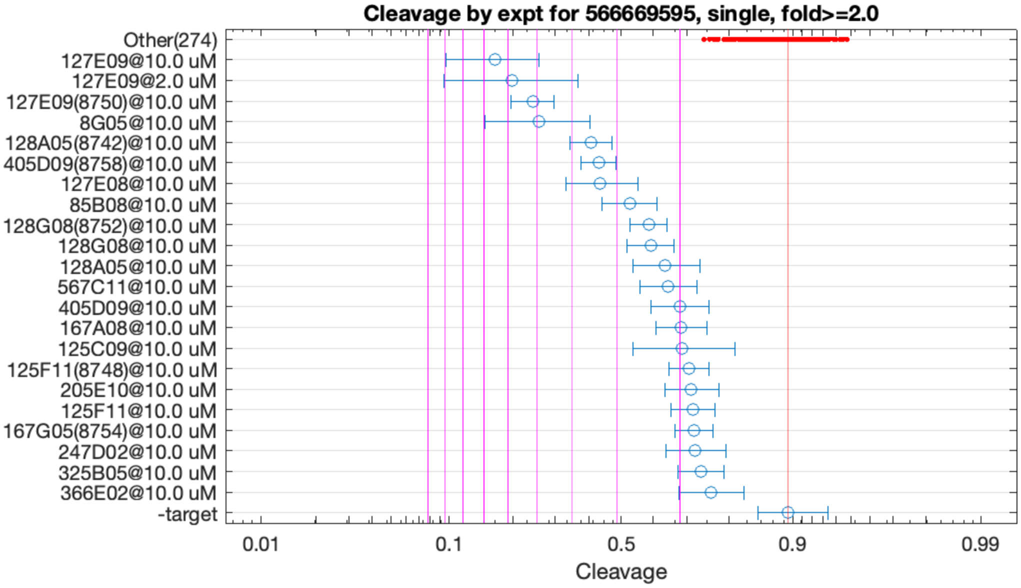


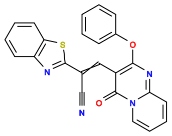

127E09

v>3.4 s=6.6

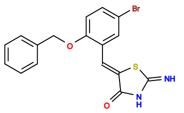

8G05

v>4.7 s=4.9

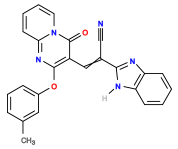

128A05

v>1.6 s=3.5

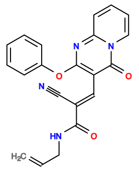

127E08

v>1.6 s=3.3

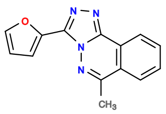

405D09

v>1.7 s=3.3

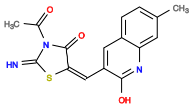

85B08

v>1.4 s=2.7

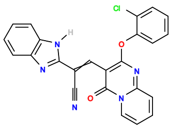

128G08

v>2.0 s=2.4

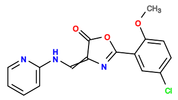

567C11

v>1.5 s=2.2

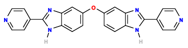

125C09

v>1.3 s=2.0
